# An optogenetic toolkit for robust activation of FGF, BMP, & Nodal signaling in zebrafish

**DOI:** 10.1101/2025.04.17.649426

**Authors:** Leanne E. Iannucci, Velanganni Selvaraj Maria Thomas, William K. Anderson, Micaela R. Murphy, Caitlin E.T. Donahue, Catherine E. Campbell, Matthew T. Monaghan, Allison J. Saul, Katherine W. Rogers

**Author notes:** These authors contributed equally (Leanne E. Iannucci, Velanganni Selvaraj Maria Thomas).

## Abstract

Cell signaling regulates a wide range of biological processes including development, homeostasis, and disease. Accessible technologies to precisely manipulate signaling have important applications in basic and translational research. Here, we present an optogenetic toolkit for signaling manipulation in zebrafish embryos. We introduce a zebrafish-optimized optogenetic FGF signaling activator and a single-transcript Nodal signaling activator, and assess them together with a previously established BMP signaling activator. We thoroughly characterize this suite of tools and demonstrate light-dependent spatiotemporal control of signaling *in vivo*. In response to ∼455 nm (blue) light, zebrafish receptor kinase domains fused to blue light-dimerizing LOV domains enable robust signaling activation with minimal inadvertent activity in the dark or at wavelengths over 495 nm. Each optogenetic tool initiates pathway-specific signaling and activates known target genes. Signaling is activated with rapid on/off kinetics, and activation strength can be tuned by adjusting light irradiance. Finally, we demonstrate spatially localized signaling activation *in vivo*. Together, our results establish this optogenetic toolkit as a potent experimental platform and provide guidelines for rapid, direct, and adjustable activation of FGF, BMP, and Nodal signaling in zebrafish embryos.

## INTRODUCTION

Cellular signaling dynamics (i.e., levels and durations) play a crucial role in orchestrating diverse biological processes including embryogenesis (Bosman and Sonnen, 2022; Hill, 2022; Kicheva and Briscoe, 2023; Sagner and Briscoe, 2017). During early vertebrate development, dynamic signaling gradients generated by fibroblast growth factor (FGF), bone morphogenetic protein (BMP), and Nodal are involved in embryonic patterning (Hill, 2022; Jones and Mullins, 2022; Liberali and Schier, 2024). These pathways signal via receptor kinases and remain at the center of many active developmental biology investigations. Established methods to manipulate FGF, BMP, and Nodal signaling such as mutating pathway genes, using heat shock promoters to drive genetically encoded agonists or antagonists, recombinant proteins, and small molecule drugs have provided foundational information about their roles in patterning. However, these strategies have limitations in spatiotemporal resolution, constraining experimental possibilities. The development and characterization of molecular technologies to precisely activate receptor kinase-based signaling *in vivo* opens new avenues of study.

Molecular optogenetic strategies use light to manipulate biological processes and have yielded novel insights across fields including developmental biology in multiple model systems (Johnson and Toettcher, 2018; Rogers and Müller, 2020). Optogenetics has been used to control processes as diverse as gene expression (LaBelle et al., 2021; Reade et al., 2017), apoptosis (Mruk et al., 2020), protein localization (Buckley, 2019; Buckley et al., 2016), and signaling (Benman et al., 2022; Čapek et al., 2019; Kainrath et al., 2017; McNamara et al., 2025; Rogers et al., 2020; Sako et al., 2016; Saul, Rogers et al., 2023). Research exploring signaling is well suited for molecular optogenetic approaches because this technology can enable precise spatiotemporal control over signaling (Farahani et al., 2021).

In the zebrafish embryo model, optogenetic tools have revealed key insights into the roles of signaling in development (Farahani et al., 2021; Rogers and Müller, 2020). The externally fertilized, translucent, microscopy-friendly zebrafish embryo is an ideal system to address developmental biology questions (Hill, 2022; Jones and Mullins, 2022; Liberali and Schier, 2024; Mullins et al., 2021) and is amenable to optogenetic approaches. Optogenetic manipulation of Nodal signaling in zebrafish has been used to define developmental windows during which Nodal is actively patterning the germ layers (Reade et al., 2017; Vopalensky et al., 2018), to investigate cellular responses to Nodal and BMP signaling (Rogers et al., 2020; Sako et al., 2016), and to examine the spatial requirements for Nodal-mediated patterning (McNamara et al., 2025). Optogenetic approaches have also provided insights into how Nodal and non-canonical Wnt signaling influence cell migration (Čapek et al., 2019; Emig et al., 2025). Finally, optogenetic tools that activate extracellular signal-regulated kinase (ERK) have been demonstrated (Benman et al., 2022; Kainrath et al., 2017; Patel et al., 2019) and used to investigate ERK’s influence on morphogenetic movements (Patel et al., 2019) and cell size/cycle regulation in the epidermis (Ramkumar et al., 2025). These compelling early studies showcase the novel research avenues that optogenetic signaling manipulators can enable. However, to address gaps in our understanding of dynamic signal interpretation (e.g., gradient-mediated patterning), optogenetic experiments introducing precise signaling inputs are ideal, but require in-depth knowledge of tool kinetics, responses to different light wavelengths and irradiances, and spatial activation capabilities. Therefore, a robustly characterized toolkit of optogenetic signaling activators with a unified design tested under identical conditions is valuable for use in developmental signaling investigations.

Toward these goals, here we present a comprehensive, direct comparison of an optogenetic toolkit to activate FGF, BMP, and Nodal signaling during zebrafish development. Our toolkit comprises a novel zebrafish-optimized FGF activator, a single-transcript Nodal activator, and an established BMP activator (Rogers et al., 2020). We show that light-mediated signaling activation by these tools *in vivo* is wavelength- and pathway-specific and leads to activation of known pathway target genes. We demonstrate that these tools activate signaling with distinct, rapid on/off kinetics and that signaling activation strength can be tuned by adjusting light irradiance. Finally, we demonstrate localized control of FGF and Nodal signaling via spatially restricted light exposure. Together, our work provides a resource for zebrafish developmental biologists (and beyond) in search of technology that enables straightforward, dynamic experimental activation of FGF, BMP, or Nodal signaling with high spatiotemporal control.

## RESULTS

### Optogenetic receptor kinase activator strategy

The optogenetic tools characterized here use the general activation strategy of light-mediated receptor kinase dimerization described in pioneering studies from 2014 (Chang et al., 2014; Crossman and Janovjak, 2022; Grusch et al., 2014; Kim et al., 2014). Endogenous FGF, BMP, and Nodal pathways signal via a similar mechanism: Ligand binding to extracellular receptor domains initiates interactions between intracellular receptor kinases, leading to phosphorylation of an effector that translocates to the nucleus and regulates target gene expression (Derynck and Budi, 2019; Heldin and Moustakas, 2016; Ornitz and Itoh, 2015). To couple signaling activation to light rather than ligand exposure, intracellular receptor kinase domains can be fused to light-responsive dimerizing proteins. Light exposure can now activate signaling by inducing receptor kinase interactions. Because these fusions lack the receptor extracellular ligand binding domain, they should be insensitive to extracellular ligand. The optogenetic tools characterized here are based on constructs developed by (Sako et al., 2016) and contain the flavin mononucleotide (FMN)-binding, blue light homodimerizing “light oxygen voltage-sensing” (LOV) domain AUREOCHROME variant from the algae *Vaucheria frigida* (VfAU1) (Takahashi et al., 2007; Toyooka et al., 2011) fused to zebrafish receptor kinase domains associated with a distinct pathway (FGF, BMP, or Nodal) (Fig. 1). The constructs used here contain a myristoylation motif for membrane targeting and a C-terminal HA or FLAG tag for immuno-detection. We refer to these constructs as “bOpto” tools for **<u>b</u>**lue light **<u>Opto</u>**genetic signaling activators.

**Figure 1:**
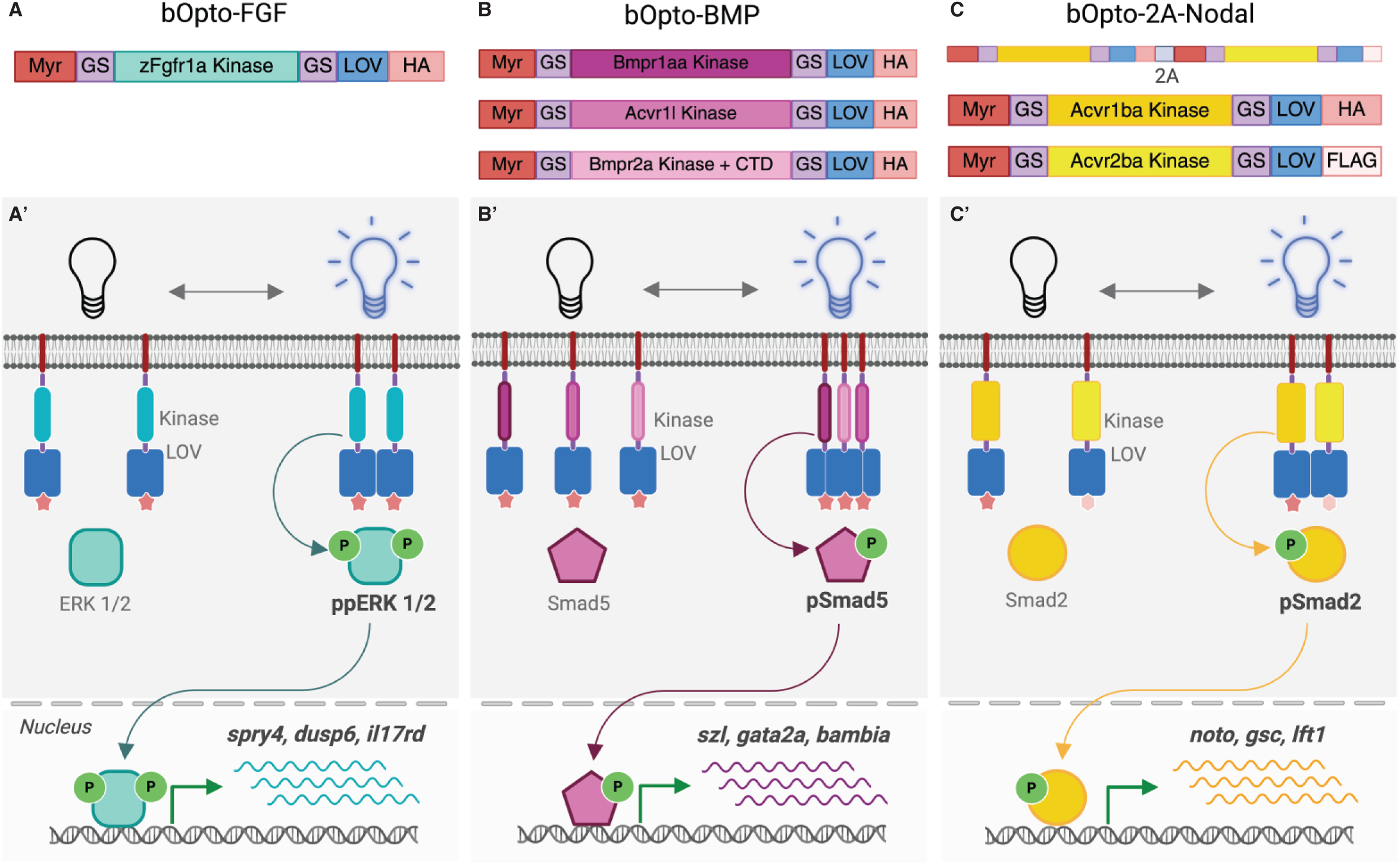
Optogenetic signaling activator toolkit. **A,B,C)** Schematic of constructs used here to activate FGF (A), BMP (B), and Nodal (C) signaling. Myr = myristoylation motif, GS = glycine/serine linker, LOV = light oxygen voltage-sensing domain, HA = hemagglutinin tag, FLAG = FLAG epitope, P = phosphate group. The single-transcript bOpto-2A-Nodal construct (C) follows the same design as -FGF and -BMP, except the type I (Acvr1ba) and type II (Acvr2ba) components are connected via a 2A peptide sequence (gray). **A’,B’,C’)** Optogenetic strategy to activate FGF (A’), BMP (B’), and Nodal (C’) signaling. Blue light-dimerizing LOV domains are fused to myristoylated receptor kinase domains. Blue light exposure should lead to receptor kinase interactions, signaling effector phosphorylation, and activation of target genes.

We first sought to create a zebrafish-optimized bOpto-FGF activator. Activation of FGF receptor kinases engages multiple intracellular signal transduction pathways, including a branch in which mediators SOS and MEK facilitate ERK phosphorylation (Ornitz and Itoh, 2015). Optogenetic ERK activators have been described in many systems (Choi et al., 2021; Csanaky et al., 2019; Dine et al., 2018; Grusch et al., 2014; Kim et al., 2014; Krishnamurthy et al., 2020; Pal et al., 2023; Rauschendorfer et al., 2021; Yadav et al., 2021). To our knowledge, three previous optogenetic ERK activators have been reported in zebrafish: 1) BcLOV-SOS_cat_, a fusion of the human RAS activator SOS_cat_ (Toettcher et al., 2013) with the temperature-sensitive blue light homodimerizing domain BcLOV4 (Benman et al., 2022), 2) psMEK^E203K^, a constitutively active hamster MEK1 fused to 400/500 nm photoswitchable Dronpa monomers modified from (Zhou et al., 2017) by the addition of an activating E203K substitution (Patel et al., 2019; Ramkumar et al., 2025), and 3) mFGFR1-MxCBD, a mouse FGF receptor intracellular domain fused to a cobalamin binding domain which homodimerizes in the dark and dissociates with green light exposure (Kainrath et al., 2017). These ERK activators feature human-, mouse-, or hamster-derived signaling components that may behave differently in different species. In addition, BcLOV-SOS_cat_ and psMEK^E203K^ act downstream of the receptor kinase (Benman et al., 2022; Patel et al., 2019; Ramkumar et al., 2025), which is useful for interrogating the roles of a specific pathway branch but may not provide information about the entire FGF receptor-mediated signaling cascade. The previous receptor-based tool, mFGFR1-MxCBD (Kainrath et al., 2017), requires constant green light exposure to prevent ectopic activation, complicating the implementation of this tool *in vivo*. Therefore, our objective was to design a bOpto tool that 1) uses a zebrafish receptor kinase to recapitulate species-specific pathway interactions, 2) functions at the level of the FGF receptor to mimic endogenous signaling, and 3) is activated by blue light (∼455 nm) and reverts to the inactive state when returned to dark. In zebrafish, the five FGF receptors function largely redundantly, although phenotypes are only observed when *fgfr1a* is knocked down in combination with any other receptor, suggesting that this receptor is a key FGF signaling mediator in the early embryo (Leerberg et al., 2019). We therefore fused the coding sequence downstream of the putative transmembrane domain from zebrafish Fgfr1a (zFgfr1a) to LOV (VfAU1) to generate bOpto-FGF (Fig. 1A,A’, Addgene #232639).

Next, we created a single-transcript version of a previously published optogenetic Nodal activator. The original LOV-based bOpto-Nodal activator is composed of two constructs: VfAU1 fused to the serine/threonine kinase domains of the zebrafish type I receptor Acvr1ba and the type II receptor Acvr2ba, respectively (Sako et al., 2016). (Note: We refer to this original two-component activator as bOpto-Nodal (Saul, Rogers et al., 2023); (Sako et al., 2016) refers to this as “Opto-acvr1b and Opto-acvr2b”, and (McNamara et al., 2025) refers to it as “optoNodal”.) To minimize the potential for dark leakiness (McNamara et al., 2025) we considered an alternative design. We created “bOpto-2A-Nodal”, a single-transcript construct that encodes both components separated by a viral PTV-1 2A peptide sequence leading to the production of two protein products (Fig. 1C,C’, Addgene #232640) (Kemmler et al., 2023; Ryan et al., 1991). Notably, this approach has been successful with other optogenetic systems (Humphreys et al., 2023; Krishnamurthy et al., 2020; Ueda et al., 2022). In addition, we replaced the HA tag on the type II-LOV component with a FLAG tag to facilitate immuno-detection of individual components (Fig. 1C). The one-transcript bOpto-2A-Nodal shows strong light-specific activity and minimal dark activity over a range of 2.5 – 40 pg mRNA; in contrast, the two-transcript system appears inactive at 2.5 pg, light-responsive at 10 pg, and dark leaky at 40 pg (Fig. 2, Supp. Fig. 1).

**Figure 2:**
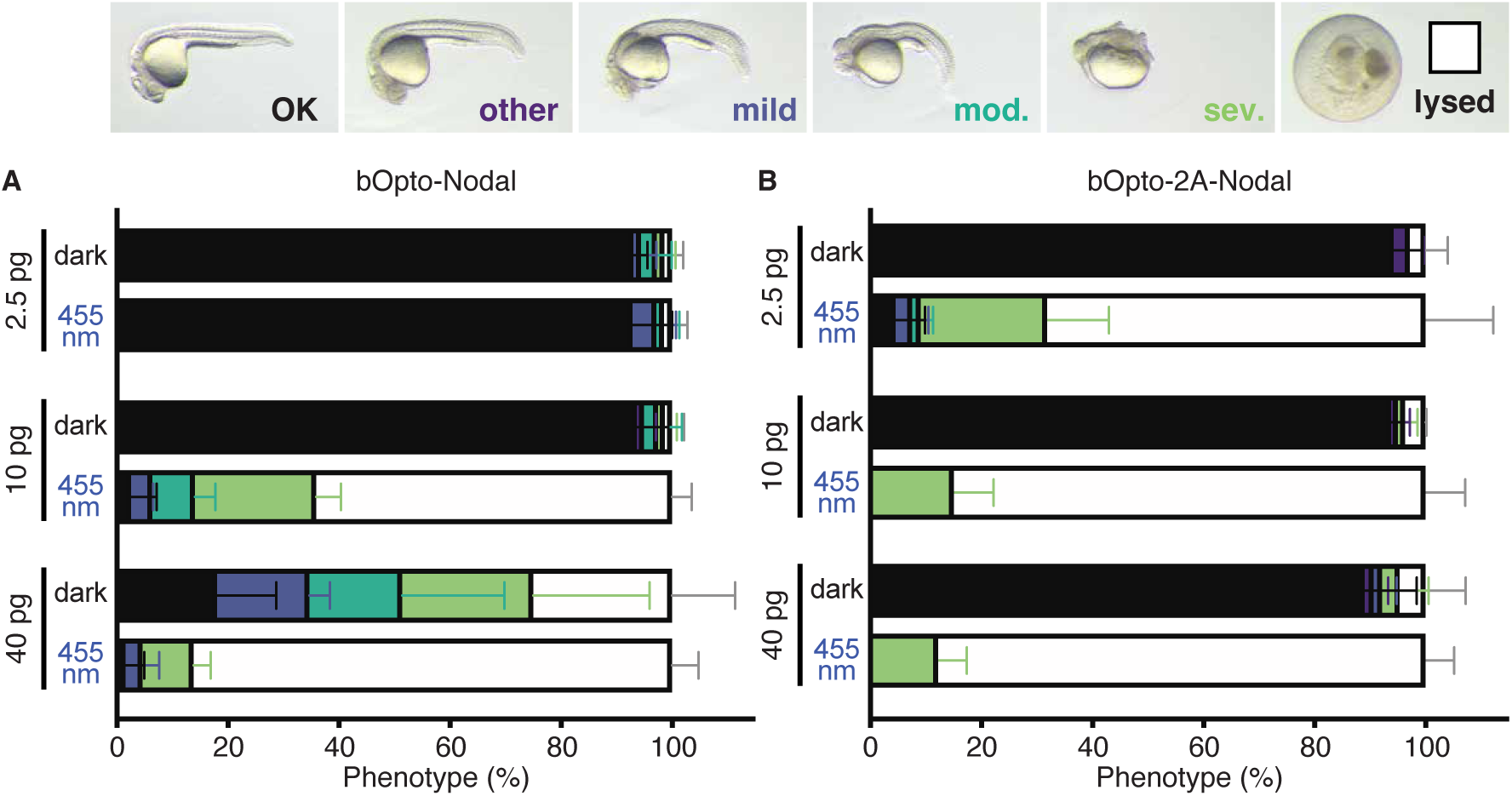
bOpto-Nodal & bOpto-2A-Nodal mRNA dosage test. Zebrafish embryos injected with the indicated amounts of “*bOpto-Nodal*” mRNA from Sako et al. 2016 **(A)** or *bOpto-2A-Nodal* mRNA **(B)** were exposed to dark or 455 nm light (50 W/m^2^) starting 1.5-2 hours post-fertilization. Note that *bOpto-Nodal* is comprised of 2 transcripts; each transcript was injected at the indicated amount. Phenotypes were scored at 1 day post-fertilization. Developmental defects and lysis are consistent with ectopic Nodal signaling. (N = 3; Supp. Fig. 1; Mean +/- SD).

The third tool in our kit is a previously established bOpto-BMP activator (Fig. 1B,B’) (Rogers et al., 2020; Saul, Rogers et al., 2023). This is a three-component system composed of LOV (VfAU1) fused to the cytoplasmic domains of the zebrafish BMP receptors Bmpr1aa, Acvr1l, and Bmpr2a. Previous work demonstrated robust BMP activation in zebrafish with minimal dark leakiness (Rogers et al., 2020). To add resolution to the previous characterization, assess additional features, and allow direct comparisons between tools, we include bOpto-BMP in these comprehensive analyses.

Together, bOpto-FGF, -BMP, and -2A-Nodal comprise the toolkit of zebrafish-specific receptor kinase-based optogenetic signaling activators that we characterize herein.

### Wavelength dependent optogenetic activators of FGF, BMP, and Nodal signaling

We first sought to determine whether bOpto tools activate their respective downstream signaling pathways specifically in response to blue light exposure (455 nm). To empirically test the wavelength dependence of bOpto tools and define “safe handling” conditions that do not inadvertently activate signaling, we injected zebrafish embryos at the one-cell stage with mRNA encoding *bOpto-FGF*, *bOpto-BMP*, or *bOpto-2A-Nodal*. We then either maintained embryos in the dark, exposed them to 455 nm light (50 W/m^2^, Supp. Fig. 2A), or exposed them to 495+ nm light (495-2200 nm, 18.51 W/m^2^, Supp. Fig. 2B) from 2 hours post-fertilization (hpf) to 1 day post-fertilization (dpf). Ectopic BMP signaling results in ventralization (V1-V4) at 1dpf (Kishimoto et al., 1997; Nguyen et al., 1998; Schmid et al., 2000), whereas embryos experiencing strong ectopic FGF or Nodal signaling have developmental defects and often lyse by 1 dpf (Feldman et al., 2002; Fürthauer et al., 1997; Fürthauer et al., 2004; Rebagliati et al., 1998; Rogers et al., 2017). At 1 dpf uninjected controls appeared morphologically normal in all conditions (Fig. 3A, Supp. Fig. 3), suggesting minimal phototoxicity. Injected embryos maintained in the dark or exposed to 495+ nm light were morphologically indistinguishable from uninjected siblings. In contrast, injected embryos exposed to 455 nm light had phenotypes consistent with strong ectopic activation of the corresponding pathway. Our phenotype data indicates that exposure to 455 nm light robustly activates bOpto tools, while dark or 495+ nm light conditions do not.

**Figure 3:**
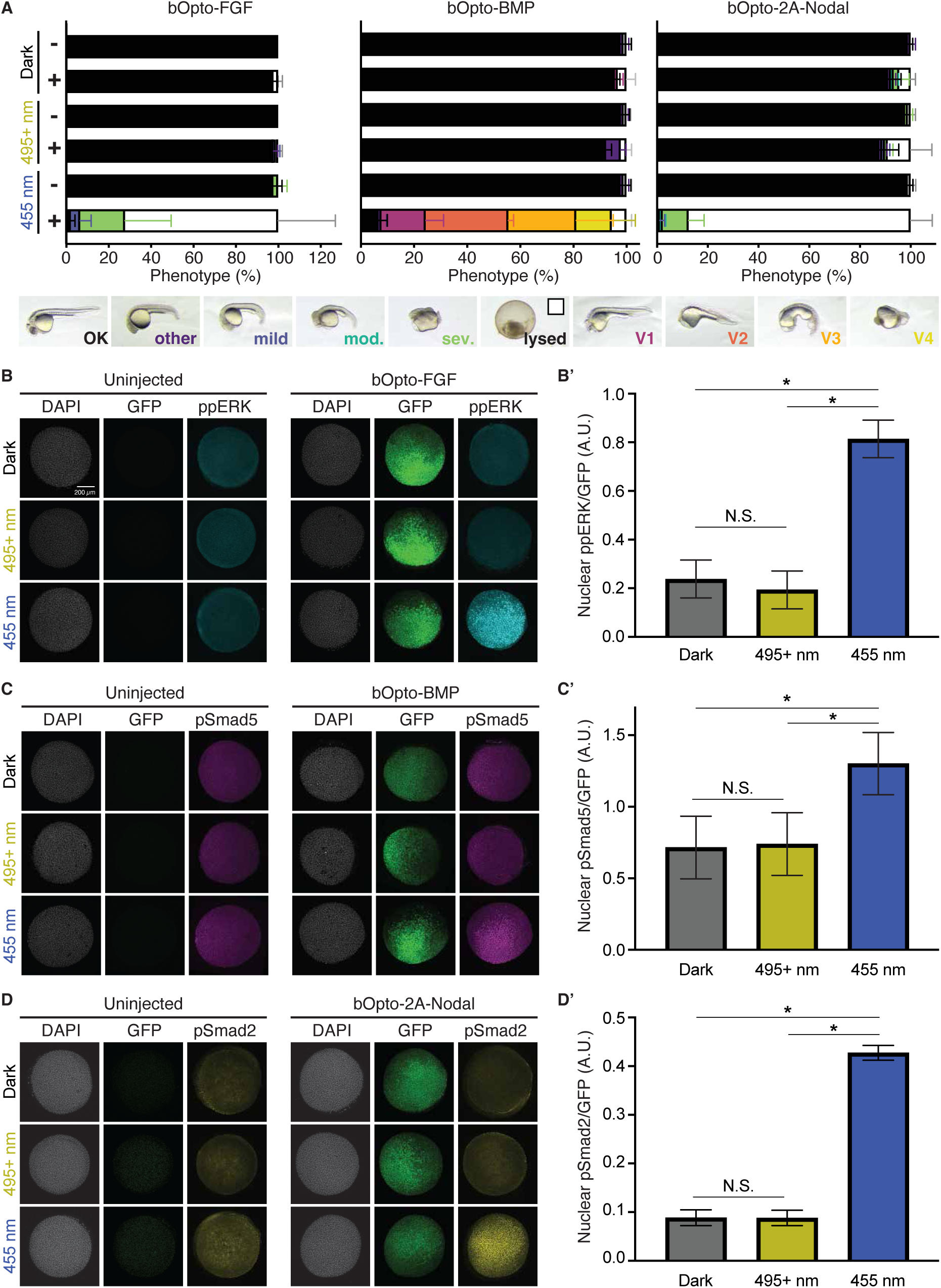
Wavelength-dependent activation of FGF, BMP, & Nodal signaling. **A)** Uninjected (-) embryos and embryos injected (+) at the one-cell stage with the indicated mRNA were exposed to dark, 495+ nm light (18.51 W/m^2^), or 455 nm light (50 W/m^2^) starting ∼2 hours post-fertilization (hpf). Pheno-types were scored at 1 day post-fertilization (dpf) (N = 3; Supp. Fig. 3; Mean +/- SD). **B,C,D)** Uninjected embryos and embryos injected with the indicated *bOpto* + *GFP* mRNA were exposed to dark, 495+ nm light, or 455 nm light starting at early gastrulation (50% epiboly - shield) for 30 min. HCR-IF was used to detect phosphorylated signaling effectors (ppERK1/2, pSmad1/5/9, & pSmad2/3 reflect FGF, BMP, & Nodal signaling, respectively). Scale bar is 200 µm. **B’,C’,D’)** Quantification of experiments shown in B,C,D. Linear-mixed model-predicted least squared means of GFP-normalized nuclear phosphorylated effector signal +/- SEM. (N = 3, * indicates *p* < 0.0001; Supp. Fig. 5).

To directly assay bOpto-mediated activation of each pathway at an earlier developmental stage, we then used hybridization chain reaction immunofluorescence (HCR-IF) (Schwarzkopf et al., 2021) to assess the phosphorylated effector corresponding to each tool’s pathway: ppERK1/2 (“ppERK”) for FGF, pSmad1/5/9 (“pSmad5”) for BMP, and pSmad2/3 (“pSmad2”) for Nodal. We assessed signaling in *bOpto*-injected and uninjected embryos exposed to dark, 455 nm light, or 495+ nm light for 30 minutes starting at early gastrulation (50% epiboly – shield). To account for non-uniform *bOpto* mRNA levels and distribution which could result in signaling heterogeneity within and between embryos, we include mRNA encoding GFP (Lim et al., 2009) in our *bOpto* injection mixes. Phosphorylated effector fluorescence intensity measured by HCR-IF was then normalized pixel-wise to GFP fluorescence intensity. Separately, to validate the co-localization of co-injected mRNAs, we co-injected *GFP* and *mScarlet-I3* (Gadella et al., 2023) mRNA and observed robust fluorescence co-localization based on structural similarity index analysis (Wang et al., 2004), supporting our normalization strategy (Supp. Fig. 4).

There were no obvious differences within ppERK, pSmad5, or pSmad2 signal in uninjected embryos across conditions (Fig. 3B-D, Supp. Fig. 5), indicating that light exposure does not affect these signaling pathways in this context. For all tools, there was a significant effect of wavelength on normalized phosphorylated signaling effector levels (*p* < 0.0001) (Fig. 3B’-D’, Supp. Fig. 5, Supplementary Materials). bOpto-FGF embryos exposed to 455 nm light showed a 3.4-fold increase in normalized ppERK levels compared to dark (*p* < 0.0001), and a 4.2-fold increase when compared to 495+ nm (*p* < 0.0001) (Fig. 3B,B’). In contrast, there were no differences between bOpto-FGF embryos exposed to 495+ nm light versus dark (*p* = 0.4744). Similarly, bOpto-2A-Nodal embryos exposed to 455 nm light showed a 4.9-fold increase in normalized pSmad2 levels compared to dark (*p* < 0.0001), and a 4.9-fold increase when compared to 495+ nm (*p* < 0.0001) (Fig. 3D,D’). There were no differences between bOpto-2A-Nodal embryos exposed to 495+ nm light versus dark (*p* = 0.9997). Finally, bOpto-BMP embryos exposed to 455 nm light similarly showed a 1.8-fold increase in normalized pSmad5 levels compared to dark (*p* < 0.0001), and a 1.8-fold increase compared to 495+ nm (*p* < 0.0001) (Fig. 3C,C’). Signaling levels in embryos exposed to 495+ nm were not different from dark exposed embryos (*p* = 0.9592). However, bOpto-BMP showed a smaller fold increase in signaling in response to blue light compared to -FGF and -2A-Nodal, possibly due to the presence of the endogenous ventral-to-dorsal pSmad5 signaling gradient which sets a higher baseline (Rogers et al., 2020; Tucker et al., 2008).

Together, our phenotype and HCR-IF results demonstrate robust bOpto-mediated signaling activation with 455 nm light exposure and negligible activation when exposed to wavelengths above 495nm (at 18.51 W/m^2^) or when maintained in the dark.

### bOpto tools activate expression of known pathway target genes

To determine whether bOpto-FGF, -BMP, and -2A-Nodal activate downstream target genes, we used hybridization chain reaction fluorescence *in situ* hybridization (HCR-FISH) (Choi et al., 2018) to assess expression of known pathway target genes. We exposed *bOpto*-injected and uninjected embryos to dark or 455 nm light (50 W/m^2^) for two hours starting before gastrulation (dome - 30% epiboly stage) to allow enough time for ample transcript accumulation. We then fixed embryos and performed multiplexed HCR-FISH (Fig. 4). We selected three well-characterized target genes per pathway: FGF = *spry4*, *dusp6*, and *il17rd* (Fig. 4A) (Fürthauer et al., 2002; Fürthauer et al., 2001; Molina et al., 2007); BMP = *sizzled*, *gata2a*, and *bambia* (Fig. 4B) (Greenfeld et al., 2021; Rogers et al., 2020); Nodal = *lefty1*, *goosecoid*, and *noto* (Fig. 4C) (Bennett et al., 2007; Bisgrove et al., 1999). In all cases, gene expression was similar when comparing uninjected dark embryos, uninjected 455 nm light exposed embryos, and injected dark embryos, suggesting negligible bOpto leakiness in the dark. In contrast, 455 nm light exposure activated gene expression in *bOpto*-expressing embryos. Together, our results demonstrate robust optogenetic activation of known target genes by bOpto-FGF, -BMP, and -2A-Nodal in zebrafish embryos, establishing that this suite of optogenetic activators provides experimental access to major signaling pathways during zebrafish development.

**Figure 4:**
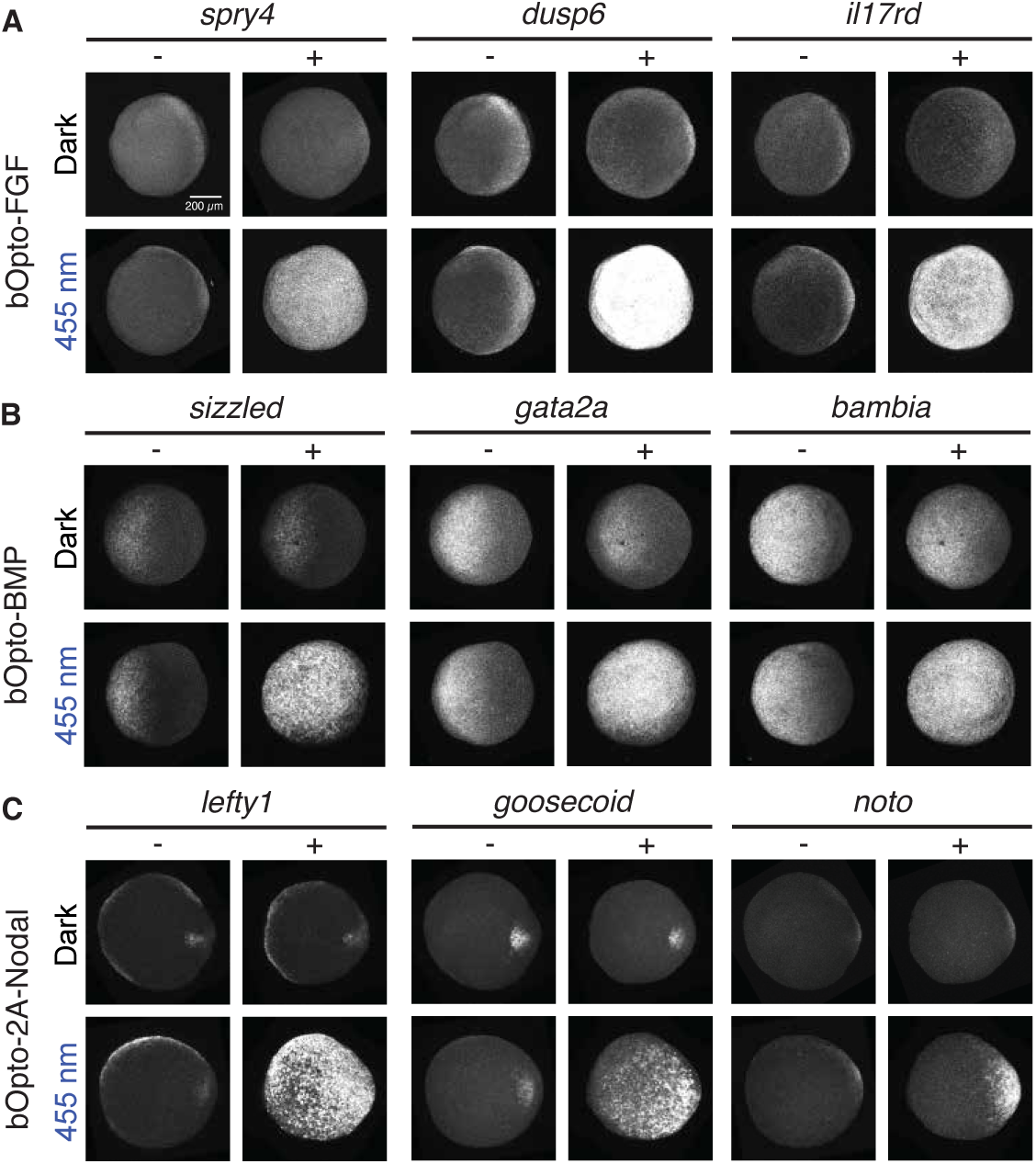
bOpto tools activate target gene expression. Uninjected embryos (-) and embryos injected (+) at the one-cell stage with mRNA encoding *bOpto-FGF* **(A)**, *bOp-to-BMP* **(B)**, or *bOpto-2A-Nodal* **(C)** were exposed to 455 nm light (50 W/m^2^, bottom rows) or dark (top rows) for 2 hours starting before gastrulation (dome - 30% epiboly). Multiplexed HCR-FISH was used to detect expression of the indicated pathway target genes. (N = 3; Scale bar is 200 µm).

### bOpto tools are pathway-specific

Receptors can engage in cross-talk, in which pathway components canonically associated with one signaling pathway are involved in transducing signaling from one or more additional pathways (Derynck and Budi, 2019; Ramachandran et al., 2018). We therefore designed an experiment to investigate whether this suite of optogenetic signaling activators exhibit cross-talk. We injected zebrafish embryos at the one-cell stage with mRNA encoding *bOpto-FGF*, -*BMP*, or *-2A-Nodal* and reared them in the dark until early gastrulation (50% epiboly – shield). We then exposed half of the injected embryos to 455 nm light (50 W/m^2^) for 30 minutes, and performed HCR-IF staining to detect ppERK, pSmad5, and pSmad2 levels for each tool (Fig. 5, Supp. Fig. 6).

**Figure 5:**
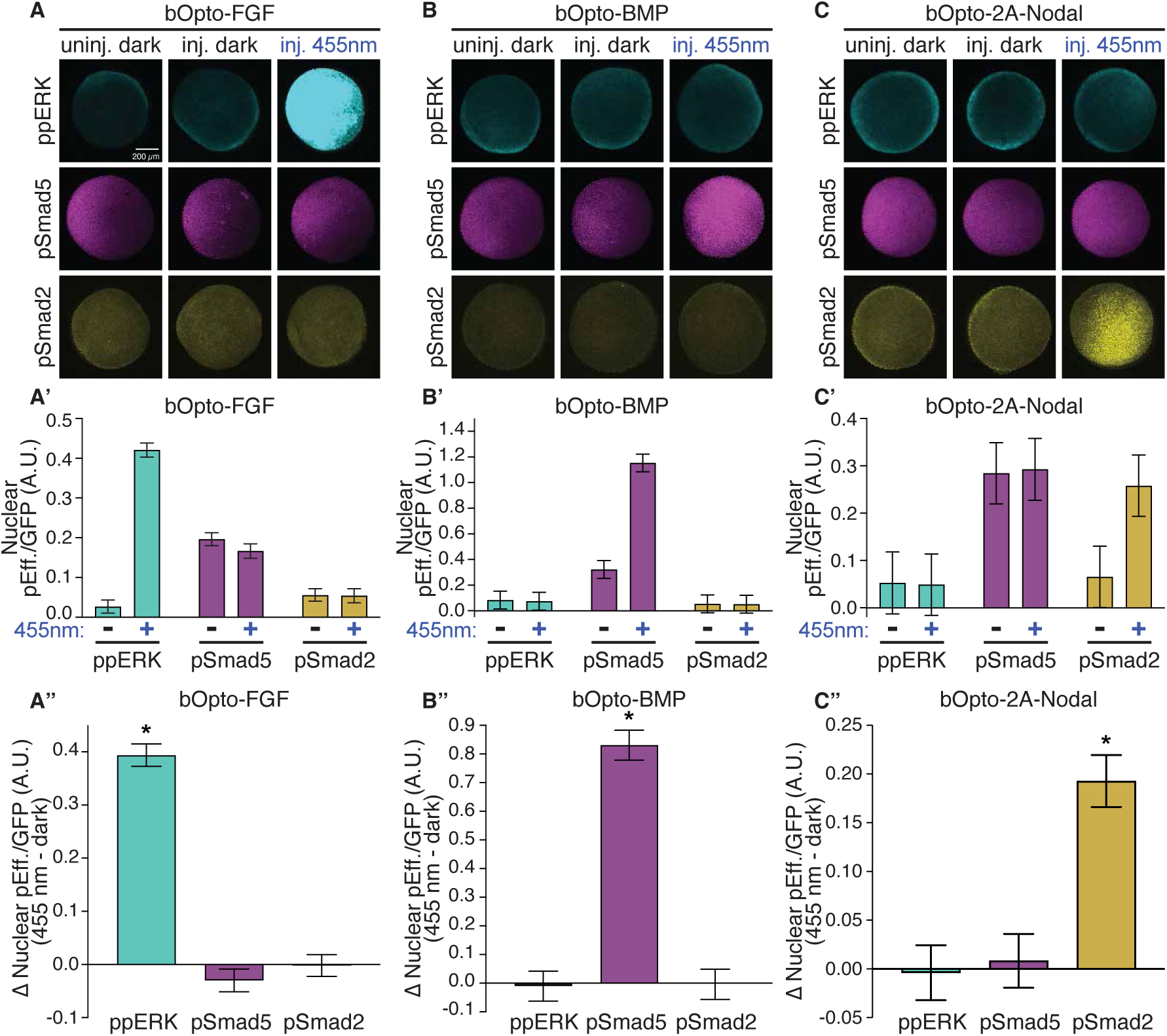
Pathway-specific optogenetic activation of FGF, BMP, & Nodal signaling. Uninjected embryos and embryos injected at the one-cell stage with mRNA encoding *GFP* & *bOpto-FGF* **(A)**, *bOpto-BMP* **(B)**, or *bOpto-2A-Nodal* **(C)** were exposed to dark or 455 nm light (50 W/m^2^) for 30 minutes starting at gastrulation (50% epiboly - shield). HCR-IF was used to detect phosphoryl-ated signaling effectors (ppERK1/2, pSmad1/5/9, & pSmad2/3 reflect FGF, BMP, & Nodal signaling, respectively). **A’,B’,C’)** Linear-mixed model-predicted least squared means of GFP-normalized nuclear phosphorylated effector signal (pEff.) +/- SEM in injected embryos. **A”,B”,C”)** Post-hoc contrast analysis of 455 nm - dark GFP-normalized nuclear phosphorylated effector signal (pEff.) +/- SEM in injected embryos. (N = 3; Supp. Fig. 6; Scale bar is 200 µm).

We noted minimal differences between uninjected embryos and injected embryos maintained in the dark, again suggesting negligible leaky dark activation consistent with our earlier wavelength dependence HCR-IF experiments (Fig. 3). bOpto-FGF embryos exposed to blue light had strongly elevated ppERK levels (15.7-fold increase) compared to dark (*p* < 0.0001); in contrast, pSmad5 (*p* = 0.5004) and pSmad2 (*p* > 0.9999) levels were unchanged between blue and dark conditions (Fig. 5A-A”, Supp. Fig. 6A). Similarly, blue light-exposed bOpto-BMP embryos had higher levels of pSmad5 compared to dark controls (3.6-fold increase, *p* < 0.0001), but no significant changes in ppERK (*p* > 0.9999) or pSmad2 (*p* > 0.9999) levels between blue and dark conditions (Fig. 5B-B”, Supp. Fig. 6B). Finally, blue light-exposed bOpto-2A-Nodal embryos showed a 4.0-fold increase in pSmad2 levels compared to dark controls (*p* < 0.0001), but no significant differences in ppERK (*p* > 0.9999) or pSmad5 (*p* > 0.9999) levels between blue and dark conditions (Fig. 5C-C”, Supp. Fig. 6C). Overall, bOpto-FGF, -BMP, and -2A-Nodal generate robust, pathway-specific optogenetic activation at the level of signaling in gastrulating zebrafish.

### bOpto tools have rapid on/off kinetics

Speed and reversibility are two benefits of optogenetic strategies (Rogers and Müller, 2020). To determine the on/off kinetics of bOpto-BMP, -FGF, and -2A-Nodal, we injected mRNA encoding each respective tool together with *GFP* mRNA into zebrafish embryos at the one cell stage. At early gastrulation stage (50% epiboly – shield) we exposed embryos to 455 nm light (50 W/m^2^) for 30 minutes, then removed light for a tool-specific dark “recovery period” that was determined empirically. We fixed embryos at multiple time points throughout the exposure and recovery phases, together with time-matched injected controls that were maintained in the dark. We then used HCR-IF to quantify phosphorylated signaling effector levels corresponding to the stimulated pathway (Fig. 6). GFP signal was used to normalize *bOpto* mRNA amount and distribution (Supp. Figs. 4 & 7). In addition, normalized effector signal was background subtracted against dark controls (Supp. Fig. 8). We observed clear signaling activation within the 30 min light exposure phase and a return to baseline after light removal for all tools (Fig. 6A-C, A’-C’).

**Figure 6:**
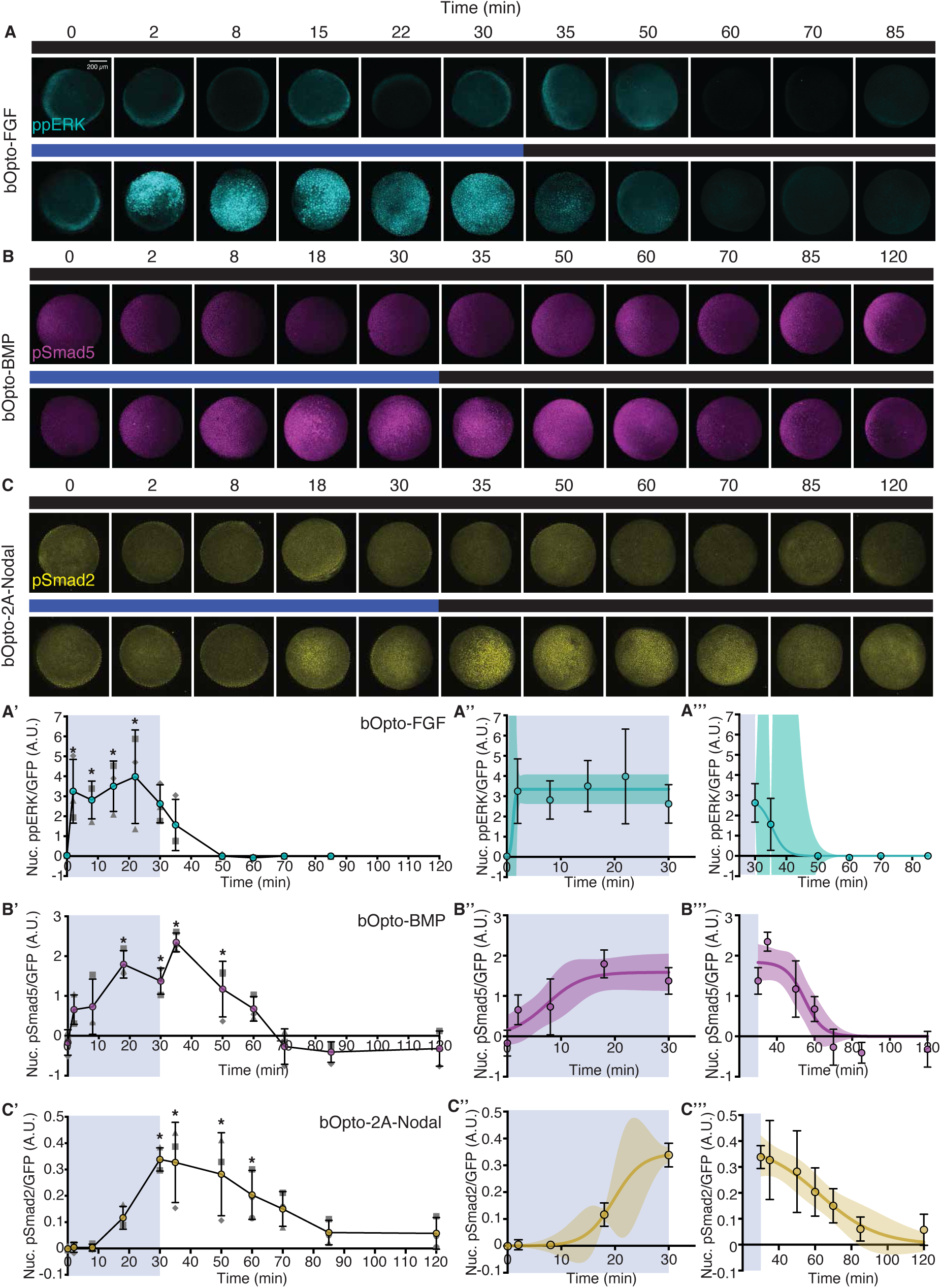
On/off kinetics of the optogenetic signaling activator toolkit. Embryos were injected at the one-cell stage with mRNA encoding *GFP* and either *bOpto-FGF*, *-BMP*, or -*2A-Nodal*. **A,B,C)** Starting at early gastrulation (50% epiboly - shield), embryos were either kept in the dark (top row) or exposed to 455 nm light (50 W/m^2^) light for 30 min (bottom row) and fixed during and after exposure. HCR-IF was used to detect phosphorylated signaling effectors (ppERK1/2, pSmad1/5/9, & pSmad2/3 reflect FGF, BMP, & Nodal signaling, respectively). (N = 3; Scale bar is 200 µm). **A’,B’,C’)** Quantification of experiments shown in A-C. Signal was GFP-normalized and subtracted against dark time-matched controls (N = 3; each N indicated by matched shape; Supp. Fig. 7 & 8; Mean +/- SD; * indicates *p* < 0.05 when compared to time = 0 min). **A’’- C’’, A’’’-C’’’)** A three-parameter logistic regression was fit to the 0-30 min data (A’’-C’’) or the >= 30 min data (A’’’-C’’’). (N = 3; Mean +/- SD; Supplementary Materials; solid line represents predicted curve fit +/- 95% CI).

First, to discern differences in “on” kinetics of the tools, we analyzed the light exposure phase of the time course (Fig. 6A”-C”). Three-parameter logistic regressions were fit to the data. From the model, three values were calculated approximating the time to reach a percentage of the curve’s upper asymptote: tON_20_ predicting initial activation, tON_50_ predicting half-maximum, and tON_90_ predicting saturation (Supplementary Materials). bOpto-FGF (curve fit R^2^ = 0.490) had the fastest on kinetics, reaching saturation within 2 minutes (Fig. 6A”). It had a tON_20_ of 0.79 min (CI: -6.27-7.86 min), a tON_50_ of 1.14 min (CI: -4.33-6.61 min), and tON_90_ of 1.68 min (CI: -2.20-5.57 min) (Fig. 6B”). bOpto-BMP (curve fit R^2^ = 0.643) had a tON_20_ of 2.76 min (CI: -2.94-8.47 min), a tON_50_ of 7.31 min (CI: 2.99-11.62 min), and tON_90_ of 14.66 min (CI: 3.42-25.91 min) (Fig. 6B”). Of the three constructs, bOpto-2A-Nodal (curve fit R^2^ = 0.966) had the slowest on kinetics: it had a tON_20_ of 16.12 min (CI: 12.25-19.99 min), a tON_50_ of 19.78 min (CI: 16.13-23.43 min), and tON_90_ of 25.59 min (CI: 14.04-37.14 min) (Fig. 6C”). Comparing initiation times (tON_20_), bOpto-2A-Nodal is ∼15 min slower than bOpto-FGF; bOpto-BMP and bOpto-FGF both initiate activation within 3 minutes of light exposure. Comparing the tools’ time to saturation (tON_90_), bOpto-FGF is ∼13 minutes faster than bOpto-BMP and ∼24 minutes faster than bOpto-2A-Nodal. It also appears that the tools that clearly reach saturation prior to the end of the 30 minute light exposure (bOpto-FGF and -BMP) maintain maximum signaling levels until light removal.

Next, to discern differences in “off” kinetics of the tools, we analyzed the dark recovery phase of the time courses (Fig. 6A’’’-C’’’). In this phase, embryos that had been exposed to the 30 minute, 455 nm light pulse were incubated in the dark for an additional 55 (bOpto-FGF) or 90 (bOpto-BMP and -2A-Nodal) minutes. The same curve fitting approach that was used to determine the on kinetics was also used here to assess the off kinetics with one conceptual alteration: We calculated the time to a defined percentage *decrease* from the upper asymptote to reflect the progressive return to baseline (Supplementary Materials). For each tool we calculated tOFF_20_ predicting time to initial deactivation, tOFF_50_ predicting time to half-maximum, and tOFF_90_ predicting approximate time to return to baseline. The off kinetics of bOpto-FGF (curve fit R^2^ = 0.791) were the most rapid of the tested tools, with a tOFF_20_ of 2.51 min (CI: -42.67-47.70 min), tOFF_50_ of 5.45 min (CI: -6.77-17.68 min), and a tOFF_90_ of 10.10 min (CI: -147.52-167.72 min). bOpto-BMP (curve fit R^2^ = 0.761) was determined to have a tOFF_20_ of 17.91 min (CI: 8.46-27.36 min), a tOFF_50_ of 24.69 min (CI: 18.71-30.67 min), and a tOFF_90_ of 35.63 min (CI: 22.54-48.72 min). Finally, bOpto-2A-Nodal (curve fit R^2^ = 0.626) had the slowest off kinetics, with a tOFF_20_ of 10.63 min (CI: -4.34-25.61 min), a tOFF_50_ of 32.67 min (CI: 22.22-43.12 min), and a tOFF_90_ of 67.60 min (CI: 33.56-101.63 min). Comparing the tools’ time to initial deactivation (tOFF_20_), bOpto-FGF began to decrease first, 8 min before bOpto-2A-Nodal and 15 min before bOpto-BMP. In contrast to bOpto-FGF and -2A-Nodal-induced signaling, which started to decrease relatively soon after light removal, bOpto-BMP-induced signaling was sustained into the dark period and then rapidly progressed towards baseline. Comparing time to return to baseline (tOFF_90_), bOpto-2A-Nodal had the slowest return time, ∼32 minutes slower than bOpto-BMP and ∼58 minutes slower than bOpto-FGF.

Each tool’s time to reach saturation in the light exposure phase (tON_90_) was faster than its time to return to baseline after the light had been removed (tOFF_90_). Additionally, the relative kinetics between the tools (e.g., bOpto-FGF fastest, bOpto-2A-Nodal slowest) was consistent for both on and off kinetics. In summary, these data demonstrate that all tools in the bOpto toolkit activate signaling within minutes and are reversible.

### Optogenetic signaling activation is dependent on irradiance

Another key advantage of optogenetic tools is the ability to adjust activation strength by modulating the amount of light energy delivered (Rogers and Müller, 2020), measured here by irradiance (W/m^2^). To characterize the irradiance dependence of bOpto tools, we injected embryos with mRNA encoding *bOpto* constructs and *GFP* at the one-cell stage, then at early gastrulation (50% epiboly – shield) exposed embryos to 455 nm light (Supp. Fig. 2A) at eight different irradiances ranging from 0.02 to 50 W/m^2^, changing roughly 3-fold between steps (Fig. 7A). Because bOpto-BMP and -2A-Nodal have slower times to saturation than bOpto-FGF (Fig. 6), we chose exposure durations of 5 minutes for bOpto-FGF and 25 minutes for bOpto-BMP and -2A-Nodal. We fixed embryos immediately after exposure and used HCR-IF to quantify the relevant phosphorylated signaling effector (Fig. 7), normalizing for mRNA level and distribution using GFP signal and background subtracting based on dark controls (Supp. Figs. 4, 9, & 10).

**Figure 7:**
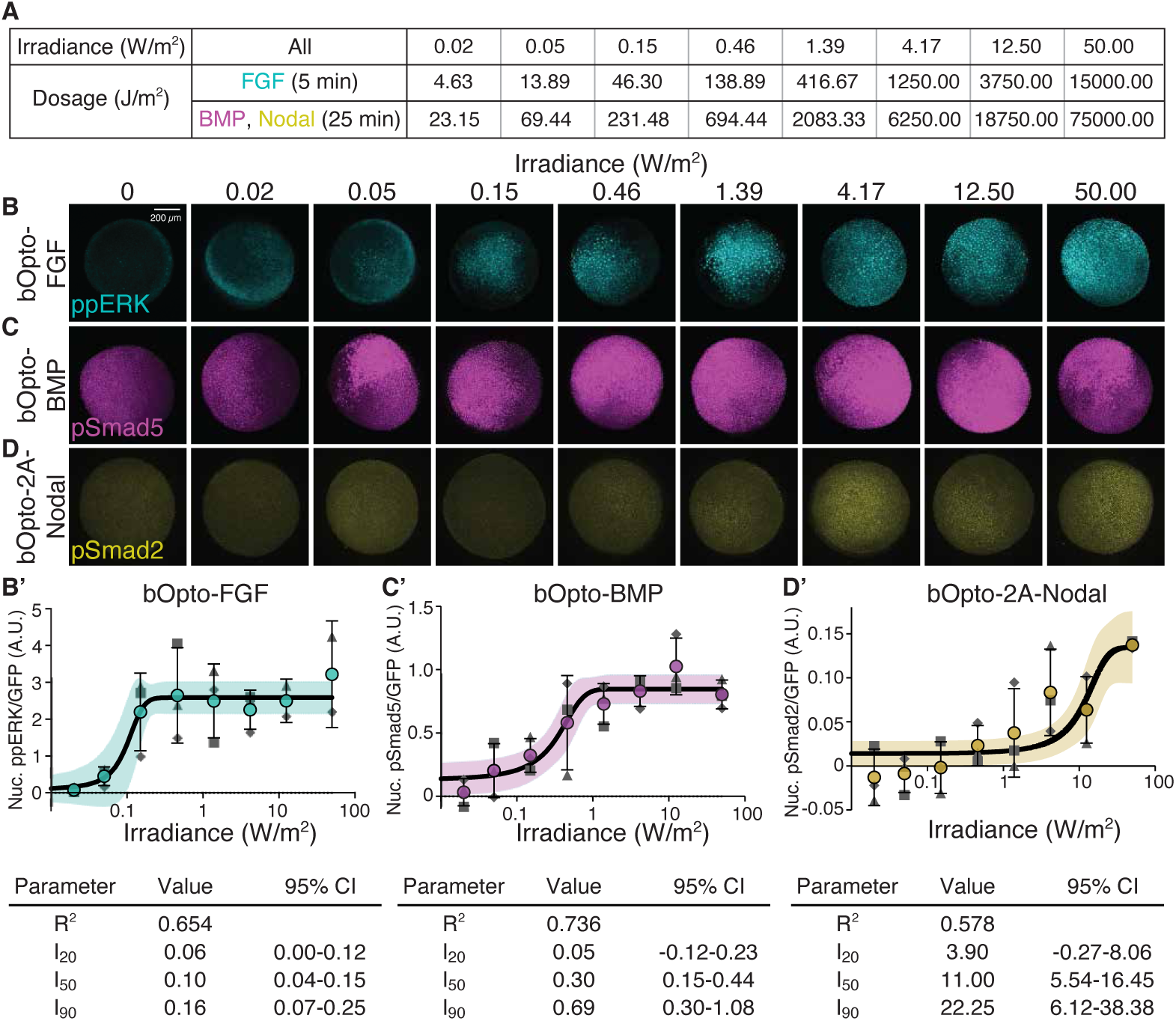
Irradiance sensitivity of optogenetic toolkit. **A)** Table of irradiance and corresponding light dosage values used in B-D’. **B,C,D)** Embryos were injected at the one-cell stage with mRNA encoding *GFP* and either *bOpto-FGF* (B), *bOpto-BMP* (C), or *bOpto-2A-Nodal* (D). Starting at early gastrulation (50% epiboly - shield), embryos were exposed to 455 nm light at the indicated irradiance for 5 min (bOp-to-FGF, B) or 25 min (bOpto-BMP and -2A-Nodal, C and D). HCR-IF was used to detect phosphorylated signaling effectors (ppERK1/2, pSmad1/5/9, & pSmad2/3 reflect FGF, BMP, & Nodal signaling, respectively). Scale bar is 200 µm. **B’,C’,D’)** Quantification of experiments in B-D. Nuclear signal was GFP-normalized and subtracted against dark controls. A three-parameter logistic regression was fit to the data. (N = 2-3; each N indicated by matched shape; Mean +/- SD; Supp. Fig. 9 & 10; solid line represents predicted curve fit +/- 95% CI; D’ shows nuclear signal only). Tables indicate goodness of fit (R^2^), the predicted irradiance (with 95% CI) at which 20 (I_20_), 50 (I_50_), and 90% (I_90_) of the curve’s upper asymptote is reached (Supplementary Materials).

For all three constructs, levels of phosphorylated signaling effectors increased with increasing light irradiance (Fig. 7). To characterize each tool’s irradiance dependence, three-parameter logistic regressions were fit to the data. From the model, three values were calculated approximating the irradiance at which a percentage of the curve’s upper asymptote is reached: I_20_ predicting initial response, I_50_ predicting half-maximum, and I_90_ predicting saturation (Supplementary Materials). bOpto-FGF (curve fit R^2^ = 0.654) was most sensitive to irradiance with an I_20_ of 0.06 W/m^2^ (CI: 0-0.12 W/m^2^), an I_50_ of 0.10 W/m^2^ (CI: 0.04-0.15 W/m^2^), and an I_90_ of 0.16 W/m^2^ (CI: 0.07-0.25 W/m^2^). bOpto-BMP (curve fit R^2^ = 0.736) had an I_20_ of 0.05 W/m^2^ (CI: -0.12-0.23 W/m^2^), an I_50_ of 0.30 W/m^2^ (CI: 0.15-0.44 W/m^2^), and an I_90_ of 0.69 W/m^2^ (CI: 0.30-1.08 W/m^2^). bOpto-2A-Nodal (curve fit R^2^ = 0.578) was the least sensitive to irradiance with an I_20_ of 3.90 W/m^2^ (CI: -0.27-8.06 W/m^2^), an I_50_ of 11.00 W/m^2^ (CI: 5.54-16.45 W/m^2^), and an I_90_ of 22.25 W/m^2^ (CI: 6.12-38.38 W/m^2^). We speculate that the bOpto-2A-Nodal curve plateaus over the range of tested irradiances; however, additional coverage between 10-100 W/m^2^ is needed to make a definite conclusion.

To directly compare the light sensitivity of these tools, we first need to account for the fact that exposure durations differed by tool. We therefore converted the I_20_ and I_90_ to dosages (J/m^2^) by multiplying them by the duration of exposure (bOpto-FGF = 5 min; -BMP and -2A-Nodal = 25 min) and calculating fold changes between tools. Based on their dosage-adjusted I_20_ values, bOpto-FGF was activated at a 5-fold lower light dosage than -BMP and a 332-fold lower light dosage than -2A-Nodal. Similarly, based on dosage-adjusted I_90_ values, bOpto-FGF saturates at a 35-fold lower light dosage than -BMP and a 1128-fold lower light dosage than -2A-Nodal. Together, our results demonstrate tunable, light irradiance-dependent activation of FGF, BMP, and Nodal signaling with this bOpto toolkit in zebrafish embryos.

### Spatially localized optogenetic signaling activation

A key advantage of optogenetic tools is the ability to manipulate biological processes with spatial precision (Rogers and Müller, 2020). Spatially resolved Nodal activation was recently achieved in zebrafish using a CRY2-based optogenetic activator (McNamara et al., 2025) and, at the level of gene expression, with the two-transcript LOV-based Nodal activator (Sako et al., 2016). Localized signaling activation has also been demonstrated for bOpto-BMP in zebrafish (Rogers et al., 2020). To determine whether bOpto-FGF and bOpto-2A-Nodal can enable spatially precise signaling activation, we co-injected *nls-Kaede* mRNA and either *bOpto-FGF* or *bOpto-2A-Nodal* mRNA at the one-cell stage. Kaede is a green-to-red photoconvertible fluorescent protein that photoconverts at 405 nm (Ando et al., 2002). bOpto-FGF embryos were reared in the dark, then starting around 50% epiboly were irradiated simultaneously with 405 + 455 nm LED light for 3 minutes using a digital micromirror device coupled to a laser scanning confocal over a square region (170 x 170 μm) (Fig. 8A-A”). A similar paradigm was followed for bOpto-2A-Nodal with the following changes in timing: 3 minutes 405 + 455 nm, followed by 24 minutes 455 nm only, followed by 3 minutes 405 + 455 nm again in an identically sized region (Fig. 8B-B”). A longer exposure was used for Nodal to account for its slower on kinetics, and 405 nm exposure was only used at the start and end to minimize UV damage while ensuring full Kaede conversion. HCR-IF was then performed to detect the FGF signaling effector ppERK or the Nodal effector pSmad2. Ectopic signaling effector activity overlapped with the photoconverted Kaede signal, demonstrating spatially localized signaling activation (Fig. 8). In summary, we demonstrate local optogenetic activation of FGF and Nodal signaling in gastrulating zebrafish using spatially restricted light exposure.

**Figure 8:**
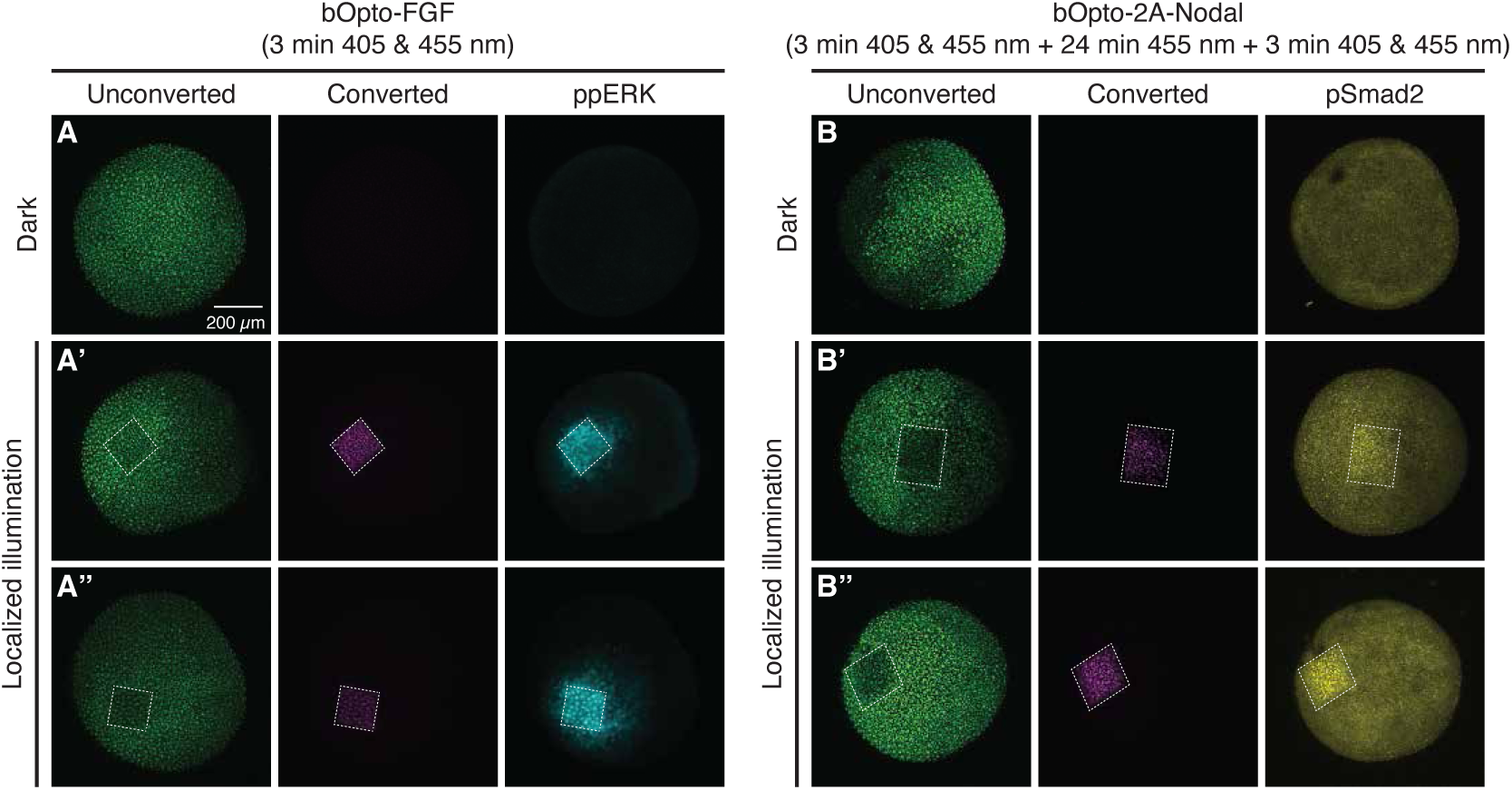
Spatially localized signaling activation. Embryos were injected at the one-cell stage with mRNA encoding the green-to-red photoconvertible fluorescent protein *nls-Kaede* and *bOpto-FGF* **(A-A”)** or *bOpto-2A-Nodal* **(B-B”)**. Starting at early gastrulation (50% epiboly) embryos were either kept in the dark (A,B) or illuminated locally (A’-A”, B’-B”) with a digital micromirror device using 405 and 455 nm LEDs for the indicated durations. HCR-IF was used to detect ppERK1/2 (FGF signaling) or pSmad2/3 (Nodal signal-ing). Dotted white lines outline photoconverted Kaede. Images from two biological replicates are shown per tool. Scale bar is 200 µm.

## DISCUSSION

### Rapid, spatially resolved optogenetic signaling activation in zebrafish

Here we introduce a zebrafish-optimized optogenetic toolkit and provide the systematic in-depth characterization necessary to apply this suite of tools. The toolkit is comprised of a novel FGF activator, a modified single-transcript Nodal activator with an improved dynamic range (Fig. 2), and a previously established BMP activator. This head-to-head *in vivo* characterization enables direct performance comparisons between the optogenetic signaling activators. We performed these characterizations in a relevant developmental context—the gastrulating zebrafish—where all three pathways are acting synergistically to pattern the embryonic axes. We show that bOpto tools enable wavelength- and pathway-specific activation of FGF, BMP, and Nodal signaling (Figs. 3 & 5) and target genes (Fig. 4) with the simple, easily controlled input of 455 nm light. Signaling can be activated with rapid on/off kinetics (Fig. 6), ectopic signaling strength can be tuned by modulating light irradiance (Fig. 7), and activation can be localized with spatially restricted illumination (Fig. 8). This optogenetic toolkit can therefore be a highly useful resource for investigations using zebrafish as a model system.

### bOpto tools do not cross-activate off-target pathways

We did not observe obvious, direct cross-activation of non-target pathways with bOpto tools using HCR-IF to assay ppERK, pSmad5, and pSmad2 (Fig. 5), but it is possible that other pathways could be inadvertently targeted by bOpto tools. Here we assessed direct effects by focusing on a relatively short 30 minute light exposure. However, with sufficiently long signaling durations, we would expect to observe inter-pathway effects because FGF, BMP, and Nodal are known to engage in many downstream interactions. For example, transcription of the BMP inhibitor Chordin is activated by Nodal and FGF signaling (Bennett et al., 2007; Koshida et al., 2002; Kudoh et al., 2004; Londin et al., 2005; Varga et al., 2007). Nodal promotes transcription of FGF ligand (Bennett et al., 2007; Mathieu et al., 2004; van Boxtel et al., 2018). FGF inhibits BMP transcription (Fürthauer et al., 1997) and can block nuclear translocation of the BMP effector pSmad1 (Kretzschmar et al., 1997). In summary, although bOpto tools are pathway-specific, experiments involving long-term signaling activation may affect non-targeted pathways due to downstream pathway interactions, though this caveat is not exclusive to optogenetic strategies.

### bOpto tools have distinct on/off kinetics and irradiance sensitivities

Our measurements revealed the temporal activation and deactivation properties of bOpto tools in zebrafish (Fig. 6, Supp. Figs. 7 & 8). All constructs in this toolkit have relatively fast on kinetics in the order of minutes. Signaling activation is reversible, decreasing to baseline after light is removed. In addition, each tool takes longer to deactivate than to activate. Despite these similarities, the tools have distinct kinetic properties: bOpto-FGF is the fastest signaling activator and bOpto-2A-Nodal is the slowest. bOpto-BMP on kinetics fall in between the other two tools, although we observed a unique feature in the off phase: Signaling was briefly sustained after light removal before rapidly returning to baseline. In contrast, there was a shorter lag between light removal and the decrease in ectopic signaling for bOpto-FGF and -2A-Nodal.

Overall, the signaling kinetics we measured for bOpto-BMP and bOpto-2A-Nodal in zebrafish are consistent with prior literature (McNamara et al., 2025; Rogers et al., 2020). As our zebrafish-optimized bOpto-FGF tool is novel, no literature comparisons are available. The kinetics we measured for bOpto-BMP are roughly consistent with those measured in the establishing study (Rogers et al., 2020). bOpto-2A-Nodal signaling kinetics appeared to be similar to those measured for the two-component bOpto-Nodal parent constructs (McNamara et al., 2025). bOpto-2A-Nodal on kinetics also appear roughly similar to those measured for a recent CRY2-based Nodal activator, though the CRY2-based construct may have faster off kinetics (Emig et al., 2025; McNamara et al., 2025). However, differences in exposure conditions (e.g., wavelength, irradiance) complicate direct comparisons.

We observed similar trends with responses to irradiance (Fig. 7, Supp. Figs. 9 & 10). Signaling activation by all tools increased as irradiance increased. bOpto-FGF activity was initiated with very low light dosages, whereas activation from bOpto-2A-Nodal required much higher light dosages, and bOpto-BMP was responsive to dosages between those that activated the FGF and Nodal tools. Previous literature also reported light dosage-dependence for bOpto-BMP (Rogers et al., 2020), the two-component bOpto-Nodal parent constructs, and the CRY2-based Nodal activator (McNamara et al., 2025). The striking differences in light dosage sensitivity we demonstrate here illustrate the importance of controlling and reporting light irradiances when using optogenetic tools.

Although further molecular characterization would be required to explain the differences in on/off kinetics and irradiance sensitivity between the bOpto tools, we speculate that several factors may contribute. First, the phosphorylation kinetics of the receptor kinase domains may differ, as well as the kinetics of phosphatases targeting pathway effectors. The kinase domain in the FGF activator is a receptor tyrosine kinase, whereas the BMP and Nodal activators are serine/threonine kinases. Receptor tyrosine kinase activity can be rapid (Farahani et al., 2023), and zebrafish embryos treated with a MEK inhibitor show strong reduction in ERK phosphorylation within just 10-20 minutes (Wilcockson et al., 2023). Another possible factor is the number of components in each bOpto activator: bOpto-FGF is a one-component system, whereas bOpto-2A-Nodal is composed of a type I and type II construct, and bOpto-BMP comprises two type I and one type II constructs (Fig. 1). This may influence the fraction of productive interactions formed upon light exposure. Finally, in the case of bOpto-2A-Nodal, the use of the 2A peptide might contribute to its observed properties. bOpto-2A-Nodal consists of two Myr-tagged constructs (Sako et al., 2016) separated by a 2A peptide (Fig. 1C). However, the localization of these components has not been directly tested. A previous report showed that a Myr-tagged construct downstream of a 2A peptide failed to localize to the membrane in HEK293T cells (Hadpech et al., 2018). If the type II-LOV construct is in fact located in the cytoplasm, activation kinetics could be delayed due to the larger search space of the cytoplasm compared to the cell membrane. Conversely, other optogenetic receptor kinase-based activators in which one component is cytoplasmic have been demonstrated to robustly activate signaling with an improved signal to noise ratio (Krishnamurthy et al., 2020; McNamara et al., 2025). We observed minimal dark leakiness and a wide dynamic range with bOpto-2A-Nodal (Fig. 2, Supp. Fig. 1). If the type II-LOV component of bOpto-2A-Nodal is cytoplasmic, this might explain the apparent improvement in dynamic range compared to previous observations from parent constructs (McNamara et al., 2025).

Overall, our measurements demonstrate that signaling activation with the bOpto toolkit is rapid, reversible, and tunable in gastrulation-stage zebrafish. This characterization therefore provides a range of useful parameters that can be experimentally manipulated to generate signaling pulses with different durations and amplitudes.

### Strategies to optimize use of bOpto signaling activators

Based on our work, we suggest four key practical recommendations for zebrafish researchers planning to incorporate bOpto signaling activators into their studies. First, mRNA injections can lead to heterogeneous distribution of bOpto tools. To mitigate these effects, we added GFP mRNA to injection mixes and used GFP signal to identify embryos injected with appropriate mRNA amounts and to normalize for differences in distribution or levels (Supp. Fig. 4). We found that GFP normalization improved the signal to noise ratio and aided interpretability (Supp. Figs. 5, 8, and 10).

Second, we show that signaling was not detectibly activated by light over 495 nm (Fig. 3, Supp. Figs. 3 & 5). Our 495+ nm light source (18.51 W/m^2^) emitted light at a lower irradiance than our 455 nm source (50 W/m^2^) (Supp. Fig. 2). However, >20 hours of 495+ nm light exposure at this relatively lower irradiance was insufficient to activate signaling based on phenotypes (Fig. 3A, Supp. Fig. 3). In contrast, as little as 0.15 W/m^2^ 455 nm light for 5 minutes robustly activated ectopic FGF signaling (Fig. 7, Supp. Fig. 10). We therefore recommend using light sources well above 495 nm to safely handle embryos (Supp. Fig. 2).

Third, our characterization of on/off kinetics and irradiance responses indicate that bOpto tools have distinct properties that should be considered when determining light exposure conditions in experimental designs. For example, bOpto-FGF is several hundred-fold more sensitive to light dosage and is also activated 10-20 minutes faster than bOpto-2A-Nodal (Figs. 6 & 7). Therefore, light exposure conditions that saturate signaling from bOpto-FGF may not even initiate signaling from bOpto-2A-Nodal.

Fourth, a potential confounding factor in optogenetic experiments is light-induced dysregulation of development. Circadian oscillations have been characterized in many organisms including zebrafish (Dekens et al., 2003; Froland Steindal and Whitmore, 2019; Krylov et al., 2021; Laranjeiro and Whitmore, 2014; Tamai et al., 2004; Whitmore et al., 2000; Ziv and Gothilf, 2006). Of the four circadian genes that we assayed in uninjected gastrulating zebrafish, three showed significant light-mediated increases in gene expression by HCR-FISH (*cry2*, *per2*, *tefa*; Supp. Fig. 11). It is unclear whether these changes in circadian gene expression affect development. However, we did not detect obvious impacts of 455 nm light exposure on signaling (Figs. 3B-D, B’-D’), gene expression (Fig. 4), or morphology (Fig. 3A, Supp. Fig. 3). We therefore speculate that the effects of light exposure on circadian gene expression did not confound the interpretation of our results. However, additional studies are needed to probe the effects of circadian gene responses in optogenetic experiments. We suggest that blue light exposed uninjected controls should be included in optogenetic experiments as a precaution.

### Prospects for optogenetic technology in developmental biology

We foresee at least three major uses of bOpto tools to address outstanding questions. First, to examine how cells decode these signaling features in a variety of contexts, our characterization of bOpto on/off kinetics and light irradiance dependence can help researchers create bespoke signaling inputs with desired amplitudes and durations. Second, it may be possible to multiplex these tools and jointly manipulate multiple pathways. This could facilitate studies examining how cells interpret combinatorial signaling (Briscoe and Small, 2015; Kicheva and Briscoe, 2023). Third, it will be valuable to use bOpto tools to impose tailored experimental signaling inputs in pathway loss-of-function mutants, as in (McNamara et al., 2025; Vopalensky et al., 2018). This could help define which features of signaling gradients are important for their patterning functions—e.g., slope, shape, dynamics—among other applications.

Finally, several technical advances would make optogenetic strategies more accessible. A current challenge with bOpto tools is that they are introduced by mRNA injection, which is transient and can lead to inconsistent distribution. Transgenic lines expressing bOpto constructs would obviate these challenges. In addition, we demonstrated local signaling activation via spatially restricted light exposure in gastrulation-stage zebrafish on the order of ∼100 μm. With further experimental refinements (e.g., focusing optics) it may be possible to activate with even finer spatial resolution. For example, cellular precision has been achieved in intact *Drosophila* embryos (Guglielmi et al., 2015; Izquierdo et al., 2018) and cell culture (Benedetti et al., 2018; Donahue et al., 2021; Passmore et al., 2025). Finally, tools to manipulate additional pathways, inhibit signaling, and orthogonal tools activated by different wavelengths would unlock further experimental possibilities.

In summary, the suite of optogenetic FGF, BMP, and Nodal activators characterized here represents a powerful experimental platform for developmental biologists and beyond. This toolkit provides a path for exploring key developmental biology questions such as how signaling dynamics and spatial gradients regulate embryogenesis. We anticipate that this work will motivate a fruitful expansion of optogenetic technology in developmental biology research.

## Acknowledgements

We thank the NIH zebrafish facility staff for maintaining our fish stocks. We also thank Jeffrey A. Farrell and his lab, Pedro P. Rocha, Sarah E. Sheppard, Leah F. Rosin, Harry A. Burgess, Patrick Müller, and the NICHD Aquatics Affinity Group for helpful discussions. Fig. 1 and Supp. Fig. 4A were created in BioRender (Rogers, K. (2025) https://BioRender.com/uybppkk).

## MATERIALS & METHODS

### Zebrafish husbandry and mRNA injections

Zebrafish husbandry and research protocols were approved by the NICHD Animal Care and Use Committee in accordance with the Guide for the Care and Use of Laboratory Animals of the National Institutes of Health. The AB zebrafish wild type strain was used in all experiments. Embryos were incubated at 28-28.5 °C in embryo medium (reverse osmosis water, Instant Ocean Sea Salt (0.25 g/L), and NaHCO_3_ (∼0.06 g/L, depending on initial pH)). For injections, mRNA was synthesized from NotI-linearized pCS2+ plasmids (see Construct Sequences, Supplementary Materials) using SP6 mMessage mMachine kits (Invitrogen AM1340). 1 nl of injection mix containing mRNA and phenol red tracer was injected through the chorion at the one-cell stage (injection mixes in Fig. 2 and Supp. Fig. 1 did not contain phenol red). For experiments in Figs. 3, 5, 6, and 7, and Supp. Figs. 5, 6, 7, 8, 9, and 10, 20 pg *GFP* mRNA (modified from (Lim et al., 2009)) was included in the same injection mix as *bOpto* mRNA. For all experiments, embryos were incubated at 28-28.5 °C immediately after injection. Between 1.5 - 2 hpf, healthy, fertilized embryos were transferred to 6-well dishes used in experiments, wrapped in aluminum foil to prevent light exposure (or immediately exposed to light for phenotyping at 1 dpf, see below and Fig. 2, Supp. Fig. 1, Fig. 3A, and Supp. Fig. 3), and returned to 28-28.5 °C (Saul, Rogers et al., 2023). At 1 dpf, embryos allocated for phenotyping were scored on Leica M80 stereoscopes. Representative phenotype images were acquired on Leica M80 stereoscopes with Flexacam C1 or C5 camera attachments (Fig. 2, Supp. Fig. 1, Fig. 3, Supp. Fig. 3). The mRNA amounts injected for each optogenetic tool (except Fig. 2 and Supp. Fig 1) are listed below:

<u>bOpto-FGF</u> (Addgene #232639): 3.5 pg

<u>bOpto-BMP (combined in injection mix)</u>

bOpto-Bmpr1aa (Addgene # 207614): 23.4 pg

bOpto-Acvr1l (Addgene # 207615): 23.4 pg

bOpto-Bmpr2a (Addgene # 207616): 40.2 pg

<u>bOpto-2A-Nodal</u> (Addgene #232640): 10 pg

In Fig. 2 and Supp. Fig. 1, for bOpto-2A-Nodal 2.5, 10, and 40 pg mRNA were injected. For bOpto-Nodal (Sako et al., 2016) experiments, injections mixes were made containing 2.5, 10, and 40 ng/ul of each component, and 1 nl was injected for a total of 2.5, 10, and 40 pg of each mRNA.

To ensure mRNA function, we included phenotype controls in all experiments. From each experiment, a subset of embryos was exposed to dark or 455 nm light starting at ∼2 hpf and phenotypes were scored at 1 dpf. We only proceeded with experiments in which these controls demonstrated that the appropriate amount of functional mRNA had been delivered. The following criteria were implemented for inclusion:

#### bOpto-FGF and -2A-Nodal experiments

Proceed if at least 60% of injected, 455 nm-exposed embryos are severely deformed or lysed. In addition to the criteria above, do not proceed if the dark injected condition exhibits more than 20% of the following phenotypes: mild to severe deformity, lysed.

#### bOpto-BMP experiments

Proceed if, in the injected, 455 nm-exposed condition, 1) there are V2 or higher embryos present, and 2) at least 60% of embryos show V1-V4 phenotypes (Kishimoto et al., 1997; Nguyen et al., 1998; Schmid et al., 2000). In addition to the criteria above, do not proceed if the dark injected condition exhibits more than 20% of the following phenotypes: V1-V4, mild-severe deformity, lysed.

In addition, rare experiments in which uninjected embryos in both dark and light conditions had high numbers of defects (over ∼30%) were excluded.

### Safe embryo handling

To protect embryos from light in “dark” conditions, 6-well dishes containing embryos were wrapped in aluminum foil. To handle light-sensitive embryos, in a windowless room with the door closed we used a ∼600 nm red light source (Supp. Fig. 2C) (A19 9W Equivalent 60W, E26 Red LED Colored Light Bulb, UNILAMP, Cat. No. B0C7YZ4KSY). In experiments requiring manipulation or visualization of light-sensitive embryos on a dissecting scope, the scope base/white light source was covered with red gel filter paper to limit wavelength emission (#E106, Rosco, Cat. No. 110084014805-E106). In experiments requiring fixation, embryos were fixed either under these red light conditions, or, for some light exposed embryos, in the LED incubator in which they were exposed to light.

### Whole embryo light exposure

Embryos were exposed to 455 nm light using the “LED incubators” described in (Saul, Rogers et al., 2023). To create a 495+ nm light source (18.51 W/m^2^, Fig. 3, Supp. Figs. 3, 5, and 11) a broadband (400 - 2200 nm) Quartz-Tungsten Halogen white light source (Thor Labs, Cat. No. QTH10/M) was used as the base emitter. A 2” x 2” square glass 495 nm long-pass filter (Thor Labs, Cat. No: FGL495S) was mounted to the source via a custom 3D printed adapter (design modified from: Thor Labs, Cat. No. SM2FH) to restrict emission wavelengths to 495+ nm. Light source emission spectra were measured over the 200-1000 nm range using a compact spectrometer (Thor Labs, Cat. No. CCS200) with a cosine corrector (Thor Labs, Cat. No. CCSB1)

Desired irradiance was confirmed prior to all experiments. Power at the sample plane across the 350-1100 nm range was measured with a digital optical power meter (Thor Labs, Cat. No. PM100D) coupled to a microscope slide sensor head (Thor Labs, Cat. No. S170C). Measured power was converted to irradiance by dividing the power by the sensor head’s active detector area. For the light irradiance experiments (Fig. 7), neutral density filters (Norman Pack of 3 Neutral Density (ND) 810551 Filters (5”), B & H Foto & Electronics Corp., Cat. No. NONDS5) were taped over the light output of the LED incubators (Saul, Rogers et al., 2023). Combinations of ND filters and adjustments to the LED power settings were tuned to achieve the desired irradiances.

### Spatial activation experiments

For spatial activation experiments (Fig. 8), embryos were dechorionated using pronase (Sigma Aldrich, Cat. No. 11459643001) at the 1 cell stage before being injected with *bOpto-FGF* (3.5 pg) + *nlsKaede* (25 pg) mRNA or *bOpto-2A-Nodal (10 pg)* + *nlsKaede* (25 pg) mRNA. Post-dechorionation and injection, embryos were incubated at 28-28.5 °C on agarose-coated petri dishes in the dark. Upon reaching 50% epiboly, embryos were carefully moved under a safe handling light source to a fresh agarose-coated dish which had wells cast into the agarose to maintain the embryos at a stable orientation (animal pole up) during light exposure (Zaucker et al., 2021).

To restrict light to a small (170 x 170 µm) square area within the embryo, an upright laser scanning confocal microscope (Zeiss LSM 800) with a digital micromirror device (DMD) (Mightex, Polygon 1000) was used as the illumination source. The light path consisted of: the emission from two LEDs (405 nm [Mightex, Part No. BLS-LCS-0405-03-22], 455 nm [Mightex, Part No. BLS-LCS-0470-14-22]) passing through beam combiners in series (Mightex Part No. LCS-BC25-0435, LCS-BC25-0515) to the DMD. The reflected light is directed to the confocal through an infinity port expander (Mightex, Part No. IPX-CHASSIS, IPX-BS-90R-10T-UF2), and passes through a W N-Achroplan 10x/0.3 objective before reaching the sample plane. A 640 nm laser was used to position the sample in the center of the field of view. A consistent focal plane for illumination was identified by finding the apex of the animal pole and then moving the z-position 125 µm down towards the center of the embryo. Using PolyScan4 (Mightex), a 170 x 170 µm region of interest (ROI) was defined in the center of the field-of-view. Stimulation to the ROI (e.g. LED power, duration) was optimized for each bOpto tool empirically. For bOpto-FGF, a 3 minute exposure period was used with the 405 nm LED set to 75% (443.3 W/m^2^) and the 455 nm LED set to 1% (9.7 W/m^2^). For bOpto-2A-Nodal, a 30 minute exposure period was used with three consecutive blocks: 1) 3 minutes of 405 nm + 455 nm, 2) 24 minutes of 455 nm only, 3) 3 minutes of 405 nm + 455 nm. Restricting 405 nm exposure to the first and last 3-minute block allowed sufficient Kaede photoconversion while minimizing any potential phototoxicity. LED power was set to 100% (578.2 W/m^2^) for 405 nm and 100% (2367.1 W/m^2^) for 455 nm.

One embryo at a time was subjected to the exposure paradigm above. After exposure, embryos were immediately fixed for downstream processing. Time-matched unexposed and uninjected embryos were also fixed as controls.

### Hybridization Chain Reaction fluorescence *in situ* hybridization (HCR-FISH, mRNA detection)

Our hybridization chain reaction fluorescence *in situ* hybridization (HCR-FISH) protocol is based on (Choi et al., 2018). Briefly, embryos were fixed in 4% formaldehyde at 4 °C overnight. After the overnight fixation, the embryos were washed 3x with 1x PBST (1x PBS + 0.1% Tween 20) to stop fixation. Embryos were manually dechorionated in 1x PBST. The dechorionated embryos were washed with 100% MeOH 3x, then stored overnight or longer at -20 °C.

Embryos were rehydrated briefly with 50% MeOH in PBST followed by 4x washes using 1x PBST. The residual PBST was carefully removed without disturbing the embryos. The embryos were pre-hybridized at 37 °C with shaking at 300 rpm for 30 minutes in pre-warmed hybridization buffer (Molecular Instruments). The HCR was multiplexed for simultaneous detection of three different genes per pathway. Three different initiators (B1, B2, and B3) were used in combination with three different pathway specific gene probes (Supp. Tables 11 & 12). The probe solution was prepared by adding equal volumes of 1 uM probe solution to the pre-warmed hybridization buffer (Molecular Instruments). Following pre-hybridization, embryos were incubated with the probe hybridization solution at 37 °C for ≥12 hours with shaking at 300 rpm. After hybridization, excess probe was removed by washing with probe wash buffer (Molecular Instruments) for 1 hour, followed by washing the embryos with 5x SSCT (5x Saline-sodium citrate + 0.1% Tween 20) for not more than 20 minutes.

Next, the pre-amplification was done by adding pre-warmed amplification buffer (Molecular Instruments) and incubating embryos at room temperature for 30 minutes. The amplification solution was prepared by snap cooling 30 pmol each of hairpin 1 and hairpin 2 for three HCR amplifiers (B1-488, B2-546, and B3-647) in separate PCR tubes and combining them with amplification buffer. The embryos were incubated with the amplification solution overnight at room temperature with gentle rocking.

After amplification the embryos were washed briefly for 10 minutes with 5x SSCT and then for a total of an hour. Embryos were then washed briefly with 1x PBST and stained with the nuclear stain DAPI (1:5000) for 2 hours at room temperature. The embryos were then washed with 5x PBST at room temperature over 1 hour, then washed briefly 3x with 1x PBS. Embryos were mounted using 1% LMP (low melting point agarose) and imaged within two weeks after processing.

### Hybridization Chain Reaction Immunofluorescence (HCR-IF, protein detection)

Our hybridization chain reaction immunofluorescence (HCR-IF) protocol is based on (Schwarzkopf et al., 2021). This protocol utilizes HCR-based signal amplification to detect protein. In brief, the embryos were fixed with 4% formaldehyde in 1x PBS overnight at 4 °C. The fixation was stopped by washing the embryos 4x with 1x PBST. Embryos were manually dechorionated in 1x PBST. Then embryos were washed with 100% MeOH 3x at room temperature to remove residual PBST. Embryos were permeabilized in 100% MeOH at -20 °C overnight or longer. The embryos were rehydrated by washing 4x using 1x PBST. Then the embryos were blocked with antibody buffer (Molecular Instruments) for either ∼4 hours at 4 °C or 1-4 hours at room temperature. The following primary antibodies were used: 1:100 rabbit anti-pSmad1/5/9 primary antibody (Cell Signaling Technology #13820S), 1:5000 rabbit anti-pSmad2/3 primary antibody (Cell Signaling Technology #8828S) and 1:5000 mouse anti-ppERK1/2 primary antibody (Sigma-Aldrich #M8159). Embryos were incubated in primary antibody in antibody buffer overnight at 4 °C. The embryos were washed with 1x PBST for ≥2 hours with gentle rocking. Next, embryos were incubated in 1:1000 donkey-anti rabbit-B1 secondary antibody (Molecular Instruments) against rabbit anti-pSmad5, or 1:500 donkey-anti rabbit-B1 secondary antibody (Molecular Instruments) against rabbit anti-pSmad2, or 1:500 donkey-anti mouse-B5 secondary antibody (Molecular Instruments) against mouse anti-ppERK for 3 hours at room temperature. After incubation in secondary antibody, the embryos were washed with 1x PBST for >30 minutes. After a brief rinse with 5x SSCT embryos were incubated in amplification buffer (Molecular Instruments) for 30 minutes. The amplifier solution was prepared by snap cooling hairpin 1 and hairpin 2 for corresponding HCR amplifiers (B1-647 or B5-647) and combining them after snap cooling in amplification buffer. Embryos were incubated in HCR amplification solution overnight at room temperature with gentle rocking, then washed with 5x SSCT for 40 minutes and rinsed briefly with 1x PBST (For pSmad2 HCR-IF experiments in Figs. 5, 6, 7, and 8 and Supp. Figs. 6, 7, 8, 9, and 10, amplification occurred at 4 °C. We found this led to better signal to noise ratio for pSmad2). Embryos were then incubated in 1:5000 DAPI for 2.5 hours and washed with 1x PBST for 1 hour, followed by a 1x PBS for 15 minutes and stored in PBS. Note that GFP survives the MeOH incubation and HCR-IF protocol and was imaged directly.

### Confocal imaging

Embryos that underwent HCR-FISH or HCR-IF were transferred to melted 1-1.2% low melting agarose (LMA) (in embryo medium) using a glass pipette. Embryos were then immediately removed from the LMA, mounted on a plastic petri dish, and oriented with the animal cap up using a stainless-steel needle. LMA was allowed to solidify and 1x PBS was added to the petri dish before imaging the embryos.

HCR-IF and HCR-FISH-processed embryos were imaged using a Zeiss LSM 800 upright confocal laser scanning microscope. The light path consisted of the emission from one or a combination of laser lines passing a W N-Achroplan 10x/0.3 objective before reaching the sample plane. Reflected light was collected with 2 GaAsP-PMT and an Airyscan63 detectors. The details of the laser lines and emission detection ranges used for all experiments are given in Supplementary Table 1. Images were acquired with 0.7x zoom in bidirectional scanning mode with 16-bit depth, with a resolution of 512×512 pixels using ZEN microscopy software. For all experiments, z-stacks were acquired with a 9 μm interval between slices, except for spatial activation experiments in which a 10 μm interval was used. Remaining imaging settings (e.g., laser power, gain, digital gain, line averaging, pixel dwell time) were consistently maintained for all samples within a biological replicate (where there existed an internal dark control) for each experiment and bOpto tool.

### Image Processing

#### Visualization Pipeline

Images from HCR-FISH gene expression experiments (Fig. 4) were processed using a custom FIJI-based pipeline (Supplementary Materials, Codes folder). In brief, for each sample, a maximum intensity z-projection was generated, and lookup table for each acquired channel was adjusted to the defined minimum and maximum values for that experiment.

HCR-IF images from wavelength dependence (Fig. 3), pathway specificity (Fig. 5), on/off kinetics (Fig. 6, Supp. Fig. 7), irradiance dependence (Fig. 7, Supp. Fig. 9), and spatial activation (Fig. 8) experiments, and HCR-FISH images from circadian experiments (Supp. Fig. 11) were processed using a custom MATLAB-based pipeline (Supplementary Materials, Codes folder). The user is prompted to supply lookup table bounds for visualization; these settings were kept consistent within each experiment and pathway. Maximum intensity z-projections were then generated for each sample and channel. Additionally, the user is prompted to identify a threshold for nuclear segmentation based on the DAPI or Kaede Green channels. Afterwards, a 3D mask defining the nuclear region is generated (hereby termed “DAPI region”) and applied voxelwise to the effector signal channel image volume. A “nuclear” maximum intensity projection is then generated.

#### Quantification Pipeline

For all HCR-IF experiments needing downstream effector signal normalization and quantification (wavelength dependence (Fig. 3, Supp. Fig. 5), pathway specificity (Fig. 5, Supp. Fig. 6), on/off kinetics (Fig. 6, Supp. Figs. 7 & 8), irradiance dependence (Fig. 7, Supp. Figs. 9 & 10)), the MATLAB-pipeline then prompts the user to specify a threshold for the GFP channel. Like nuclear thresholding, a 3D mask defining the “GFP region” is generated. The intersection between the DAPI and GFP regions was calculated as the “DAPI+GFP region”. The effector signal channel image volume is divided voxelwise by the GFP channel image volume, creating a “normalized signal” data volume. The following median values are then calculated for the imaged volume: 1) raw effector signal within the DAPI region, 2) GFP-normalized effector signal within the DAPI region, 3) raw effector signal within the GFP region, 4) GFP-normalized effector signal within the GFP region, 5) raw effector signal within the DAPI+GFP region, and 6) GFP-normalized effector signal within the DAPI+GFP region. Maximum z-projections of masks are exported for quality control assessment. Nuclear and GFP thresholds were kept consistent for all samples within a biological replicate (where there existed an internal dark control) for each experiment and bOpto tool. The median GFP-normalized effector signal within the DAPI+GFP region was used in all downstream analyses.

Statistics were performed using JMP (v17.0.0) and graphs were generated using GraphPad Prism (v10.2.3). Fold changes are defined as the considered value divided by the reference value.

#### Quality Control

Output data was then assessed for quality by 1-2 authors before additional processing. If any of the following criteria were met through screening maximum intensity projections, that sample was thrown out: 1) damage was apparent in the DAPI channel spanning 1/3 of the embryo, 2) any part of the embryo was out of frame, 3) a portion of an errant second embryo was in frame, 4) the embryo was mounted on an angle (>40 degrees off axis), 5) the z-stack did not encompass the entire embryo. If the following criterion was met through screening maximum intensity projections, that entire exposure group within the biological replicate was removed: saturation-induced “streaking” present in any one channel. If the following criterion was met through screening masks, that sample was removed: <5% of the embryo was covered by the appropriate GFP mask, indicating a sample did not receive enough mRNA for analysis.

Any embryo imaged that passed all inclusion criteria was defined as a technical replicate within an exposure condition and biological replicate for the appropriate experiment. Sufficient “trials” were run for each experiment such that 3 biological replicates were acquired for all assayed conditions within each experiment. GFP-normalized signal for all technical replicates within an exposure condition, biological replicate, and bOpto tool were averaged together and processed as follows for the corresponding experiments:

#### Wavelength Dependence

For each bOpto tool, the mean GFP-normalized signal for all samples within each wavelength (Dark, 495+ nm, 455 nm) and biological replicate was fit by a linear mixed-effects model (Fixed Effect: Wavelength, Random Effect: Biological Replicate; Personality: Standard Least Squares, Method: REML). Predicted least square means and standard error were calculated for each wavelength. Where warranted by a significant (*p* < 0.05) fixed effect, *post hoc* comparisons were performed between all exposure combinations and corrected for multiple comparisons with Tukey’s HSD method where *p* < 0.05 was considered to be statistically significant (Supp. Tables 2-4).

#### Pathway Dependence

For each bOpto tool, the mean GFP-normalized signal for all samples within each pathway effector (ppERK, pSmad5, pSmad2), exposure condition (Dark, 455 nm) and biological replicate was fit by a linear mixed-effects model (Fixed Effect: Pathway, Exposure, Pathway*Exposure, Random Effect: Biological Replicate; Personality: Standard Least Squares, Method: REML). Predicted least square means and standard error were calculated for each wavelength. Where warranted by a significant (*p* < 0.05) fixed interaction effect (Pathway*Exposure), *post hoc* contrast testing was performed between the model-predicted least square means difference of 455 nm exposed minus dark within each pathway, and outputs were adjusted with a Bonferroni correction to account for multiple comparisons. *p* < 0.05 was considered to be statistically significant (Supp. Table 5).

#### On/Off Kinetics

For each bOpto tool, the mean GFP-normalized signal for all 455 nm exposed samples within each time point (FGF: 0, 2, 8, 15, 22, 30, 35, 50, 60, 70, and 85 min; BMP/Nodal: 0, 2, 8, 18, 30, 35, 50, 60, 70, 85, and 120 min) and biological replicate was subtracted by its time-matched dark control. The dark-subtracted, normalized data was then analyzed by a one-way ANOVA with a fixed effect of time. Where warranted by a significant (*p* < 0.05) fixed effect, *post hoc* comparisons were performed between each time point and time = 0 min with Dunnett’s method for multiple comparison correction where *p* < 0.05 was considered to be statistically significant (Supp. Tables 6-7).

Next, models were fit to the dark-subtracted, GFP-normalized dataset to calculate estimates for on and off kinetic parameters. One model was generated for the “On” portion of the data (time <= 30) and a second model was fit to the “Off” portion of the data (time >= 30). We chose to fit a three-parameter logistic regression to each data subset. The equation for this model is defined as:

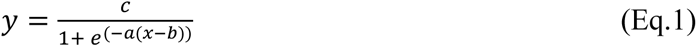

where y is the dark-subtracted normalized signal [AU], x is time [min], a is the growth rate [min^−1^], b is the inflection point [min], and c is the upper asymptote [AU].

This curve fit has two underlying assumptions: 1) the general behavior of the bOpto tools will follow a tri-phasic lag-growth/decay-plateau behavior, and 2) the lower asymptote will be 0 (i.e., for “ON” the dark subtracted, normalized signal begins at zero; for “OFF” the dark subtracted, normalized signal will eventually return to zero at some point in time). R^2^ was reported as the goodness of fit for each model. The predicted value of each factor (a, b, c) and its corresponding 95% confidence interval is also reported (Supp. Tables 8-9).

Additionally for the model fitting the “on” data, we used the predicted equation to solve for time at various points along the curve to more easily describe the activation kinetics. (Supplementary Materials) We took the value for the upper asymptote, c, and calculated three different percentages (20%, 50%, and 90%) of that value. We then used the curve fit equation to predict the time it takes from the start to reach those values. These conceptually represent: tON_20_ predicting initial activation, tON_50_ predicting half-maximum, and tON_90_ predicting saturation. Additionally, for the model fitting the “off” data, we performed an analogous analysis with one modification: we calculated the time from light removal to reach three different percentage *decreases* of the upper asymptote (20%, 50%, and 90% decreases). These conceptually represent: tOFF_20_ predicted time to initial deactivation, tOFF_50_ predicted time to half-maximum, and tOFF_90_ predicted approximate time to return to baseline. The predicted value of each tON and tOFF factor and its corresponding 95% confidence interval was reported in (Supp. Tables 8-9).

#### Irradiance Dependence

For each bOpto tool, the mean GFP-normalized signal for all samples within each irradiance (0, 0.02, 0.05, 0.15, 0.46, 1.39, 4.17, 12.50, 50.00 W/m^2^) and biological replicate was subtracted by its round-matched dark control (Supp. Table 10). Data were fit with the same three-parameter logistic regression model used in the on/off kinetics experiments. (Eq. 1) The predicted value of each factor (a, b, c) and its corresponding 95% confidence interval was reported. From the model, three values were calculated approximating the irradiance at which a percentage (20%, 50%, and 90%) of the curve’s upper asymptote is reached: I_20_ predicting initial response, I_50_ predicting half-maximum, and I_90_ predicting saturation. The predicted value of each factor and its corresponding 95% confidence interval was reported in (Supp. Table 10).

---

**Supplementary Figure 1:**
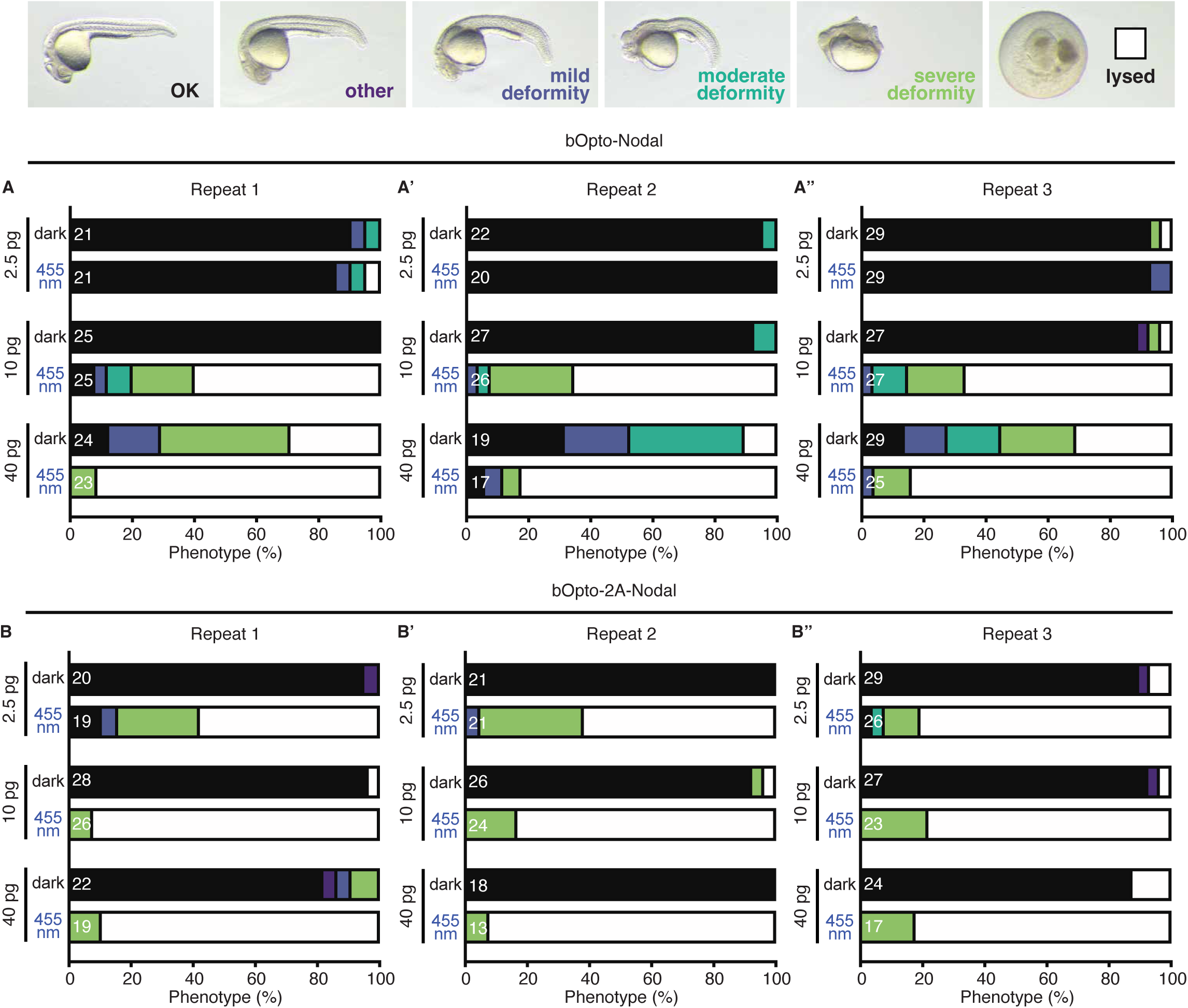
bOpto-Nodal & bOpto-2A-Nodal mRNA dosage test biological replicates. Zebrafish embryos injected with the indicated amounts of “*bOpto-Nodal*” mRNA from Sako et al. 2016 **(A-A’’)** or *bOpto-2A-Nodal* mRNA **(B-B’’)** were exposed to dark or 455 nm light (50 W/m^2^) starting 1.5-2 hours post-fertilization. Note that *bOpto-Nodal* is comprised of 2 transcripts; each transcript was injected at the indicated amount. Phenotypes were scored at 1 day post-fertilization. Developmental defects and lysis are consistent with ectopic Nodal signaling. Three biological replicates were performed and are shown here. White numbers indicate the total number of embryos scored in each condition for that experiment. Figure 2 shows combined data. Phenotype legend images shown here are the same images shown in Figure 2.

**Supplementary Figure 2:**
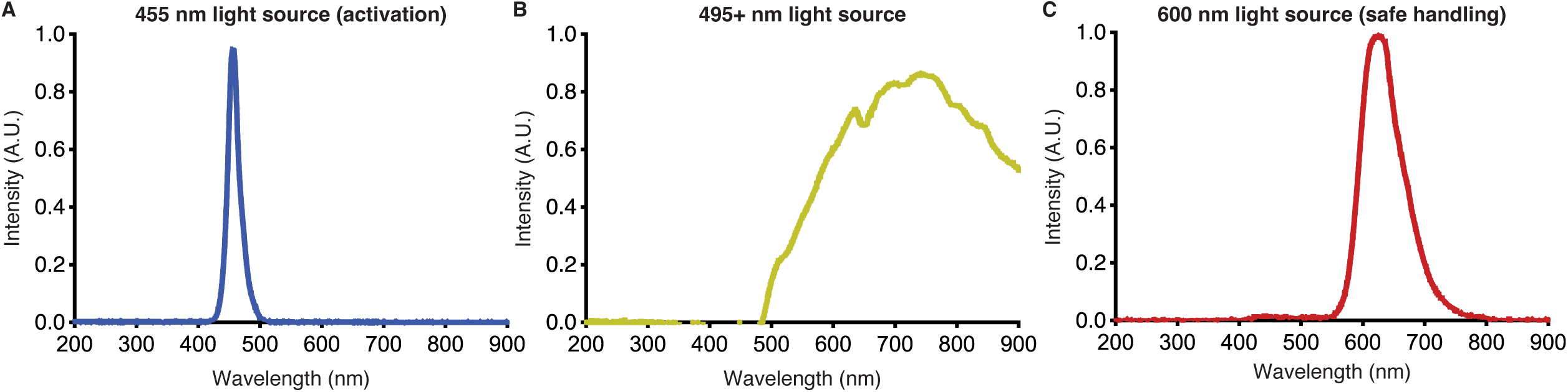
Light source spectra. Measured spectra for **A)** the 455 nm light used in uniform blue light exposure experiments (all figures except Fig. 8), **B)** the 495+ nm used in Figs. 3 and Supp. Figs. 3, 5, and 11, and **C)** the 600 nm light source used during handling of embryos expressing bOpto constructs.

**Supplementary Figure 3:**
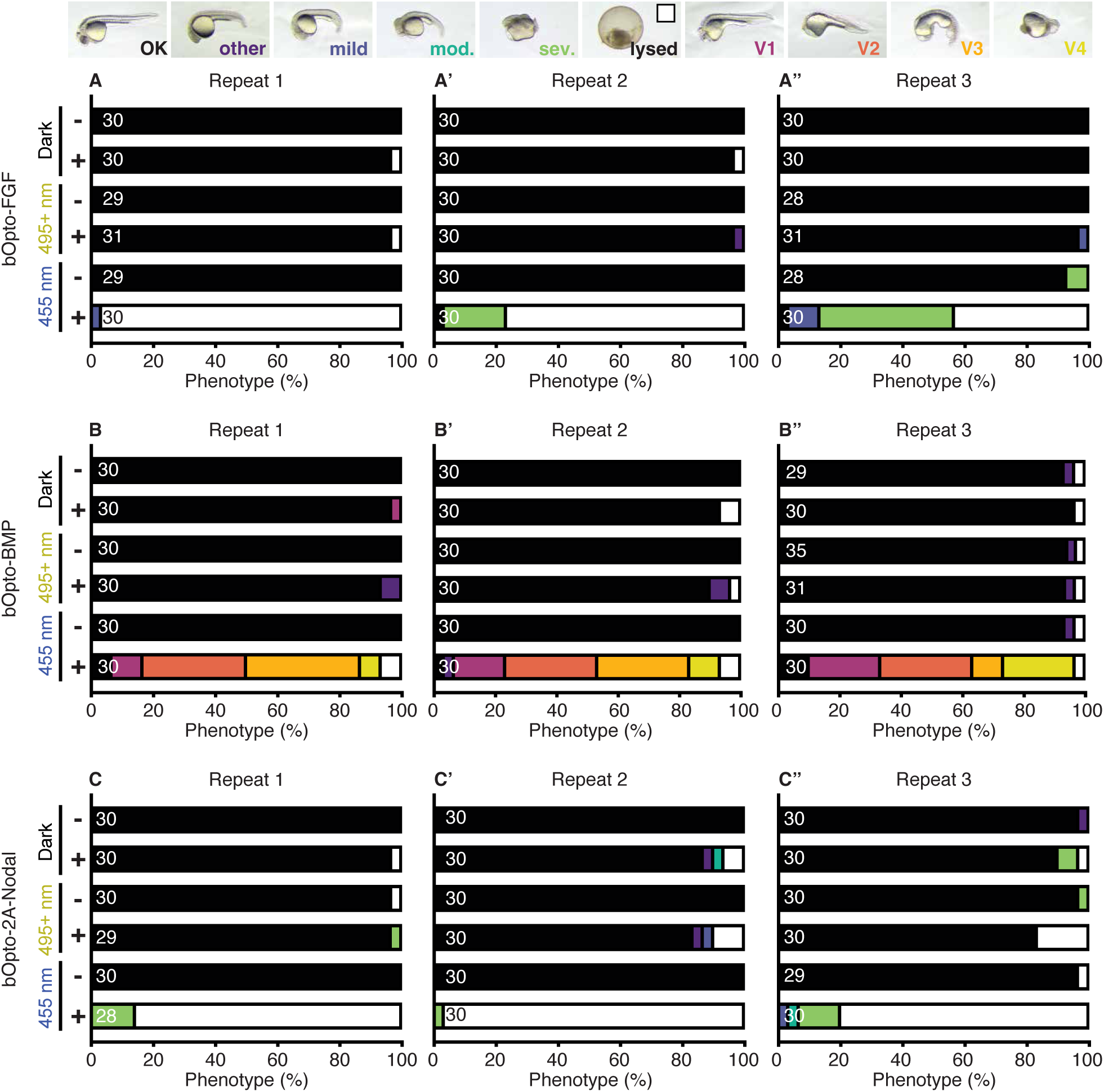
Wavelength-dependent activation of FGF, BMP, & Nodal signaling experiment repeats. **A-C”)** Uninjected embryos (-) and embryos injected (+) at the one-cell stage with the *bOp-to-FGF* (A-A”), *bOpto-BMP* (B-B”), or *bOpto-2A-Nodal* (C-C”) mRNA were exposed to dark, 495+ nm light (18.51 W/m^2^), or 455 nm light (50 W/m^2^) starting 1.5-2 hours post-fertilization. Phenotypes were scored at 1 day post-fertilization. Biological repeats are shown. Numbers indicate the total number of embryos scored in each condition for that experiment. Figure 3A shows the combined data from the three biological repeats. Phenotype legend images shown here are the same images shown in Figure 3A.

**Supplementary Figure 4:**
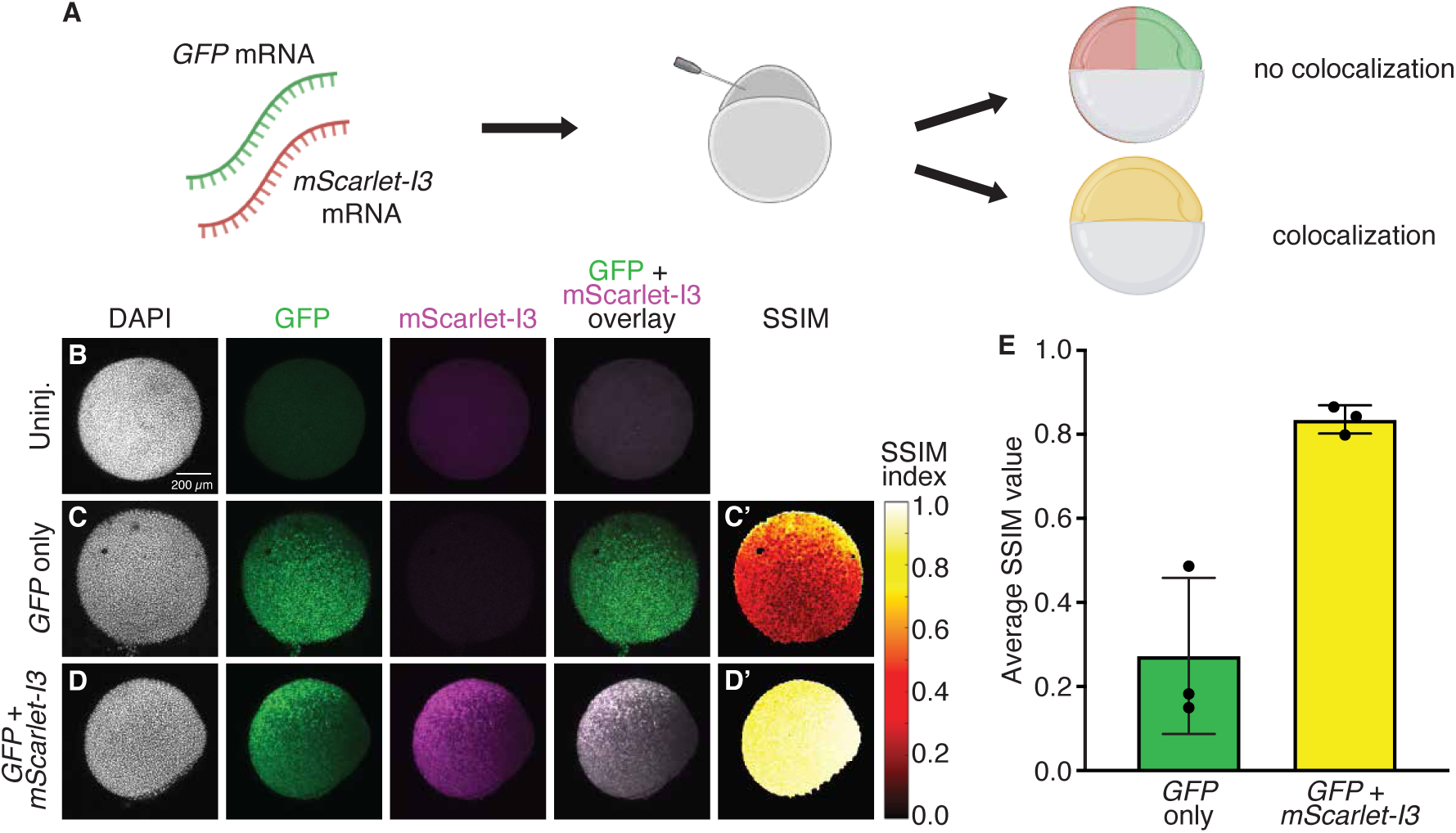
Co-localization of fluorescent protein signals from co-injected mRNAs. **A)** Zebrafish embryos were injected at the one-cell stage with 20 pg *GFP* mRNA only or co-injected with 20 pg *GFP* + 40 pg *mScarlet-I3* mRNA. Embryos were fixed at shield stage, DAPI stained, and imaged to determine the extent of overlap between fluorescent protein signals. Three biological replicates were perfomed with 2-4 technical replicates per trial. **B,C,D)** DAPI, GFP, mScarlet-I3 signal and overlays are shown for representative uninjected (B), GFP-only (C), and GFP + mScarlet-I3 (D) images. **C’,D’)** Structural similarity index (SSIM) analysis was applied to *GFP*-only (C’) and *GFP* + *mScarlet-I3* (D’) embryos to quantify fluorescent signal overlap (Wang et al., 2004). SSIM compares the pixel-wise similarity of two images (GFP and mScarlet-I3 here). Higher values indicate greater similarity. **E)** SSIM quantification. Average SSIM was caculated for each embryo. Dots indicate average of one biological replicate. Scale bar is 200 µm.

**Supplementary Figure 5:**
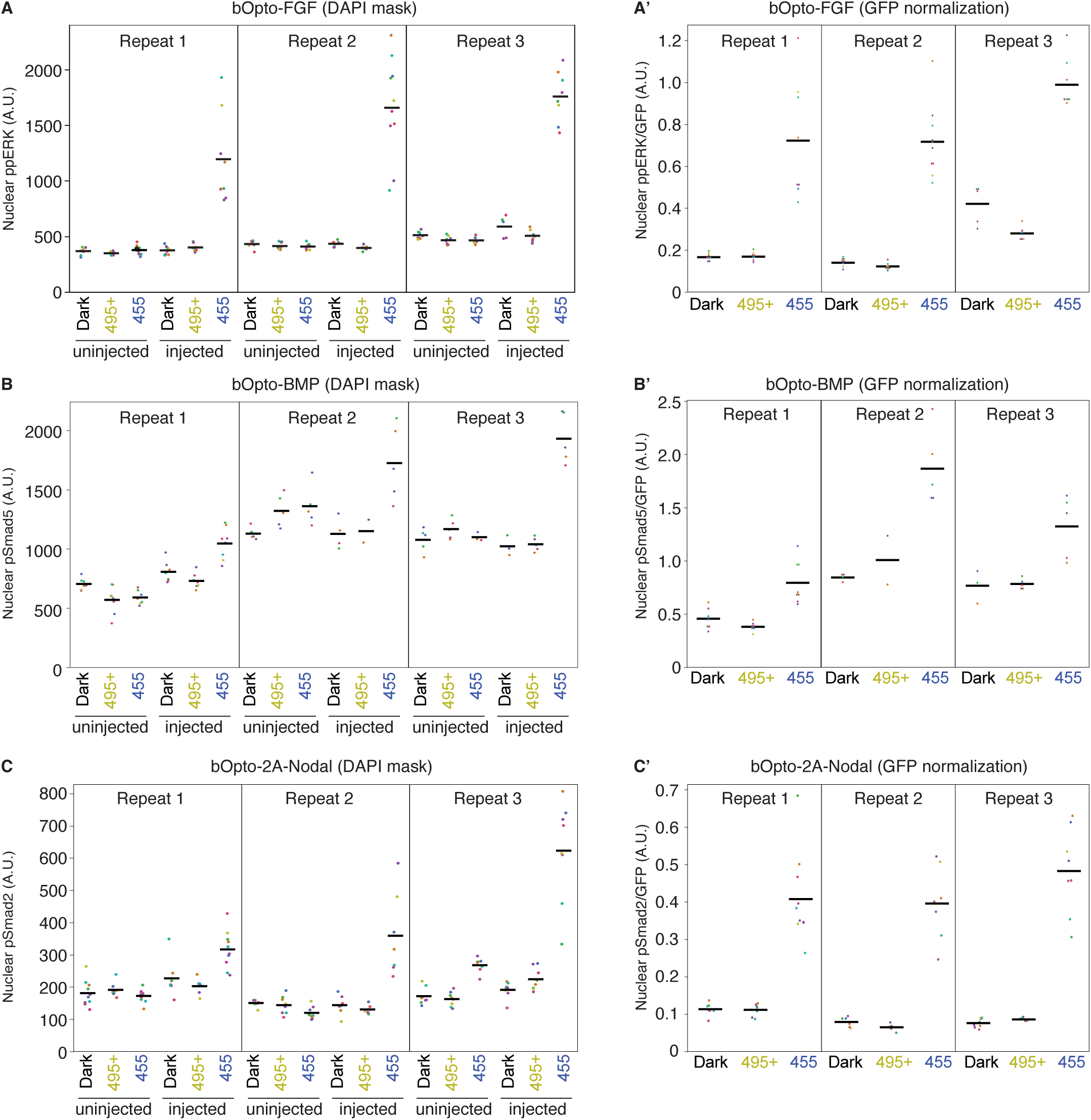
Wavelength-dependent activation of FGF, BMP, & Nodal signaling quantification. Embryos injected at the one-cell stage with the indicated mRNA (+ *GFP* mRNA) were exposed to dark, 495+ nm light (18.51 W/m^2^), or 455 nm light (50 W/m^2^) starting at early gastrulation (50% epiboly - shield) for 30 min. HCR-IF was used to detect activated signaling effectors (ppERK1/2, pSmad1/5/9, & pSmad2/3 reflect FGF, BMP, and Nodal signaling, respectively). **A,B,C)** Raw phosphorylated signaling effector intensity was measured in each DAPI-positive nuclear pixel. Each dot represents the median nuclear pixel intensity in one embryo. Black lines represent the mean nuclear intensity of all embryos in the indicated condition. **A’, B’, C’)** Phosphorylated signaling effector intensity in each DAPI + GFP-positive pixel was divided by the corresponding GFP intensity. Each dot represents the median GFP-normalized nuclear pixel signal in one embryo. Black lines represent the mean GFP-normalized nuclear signal of all embryos in the indicated condition.

**Supplementary Figure 6:**
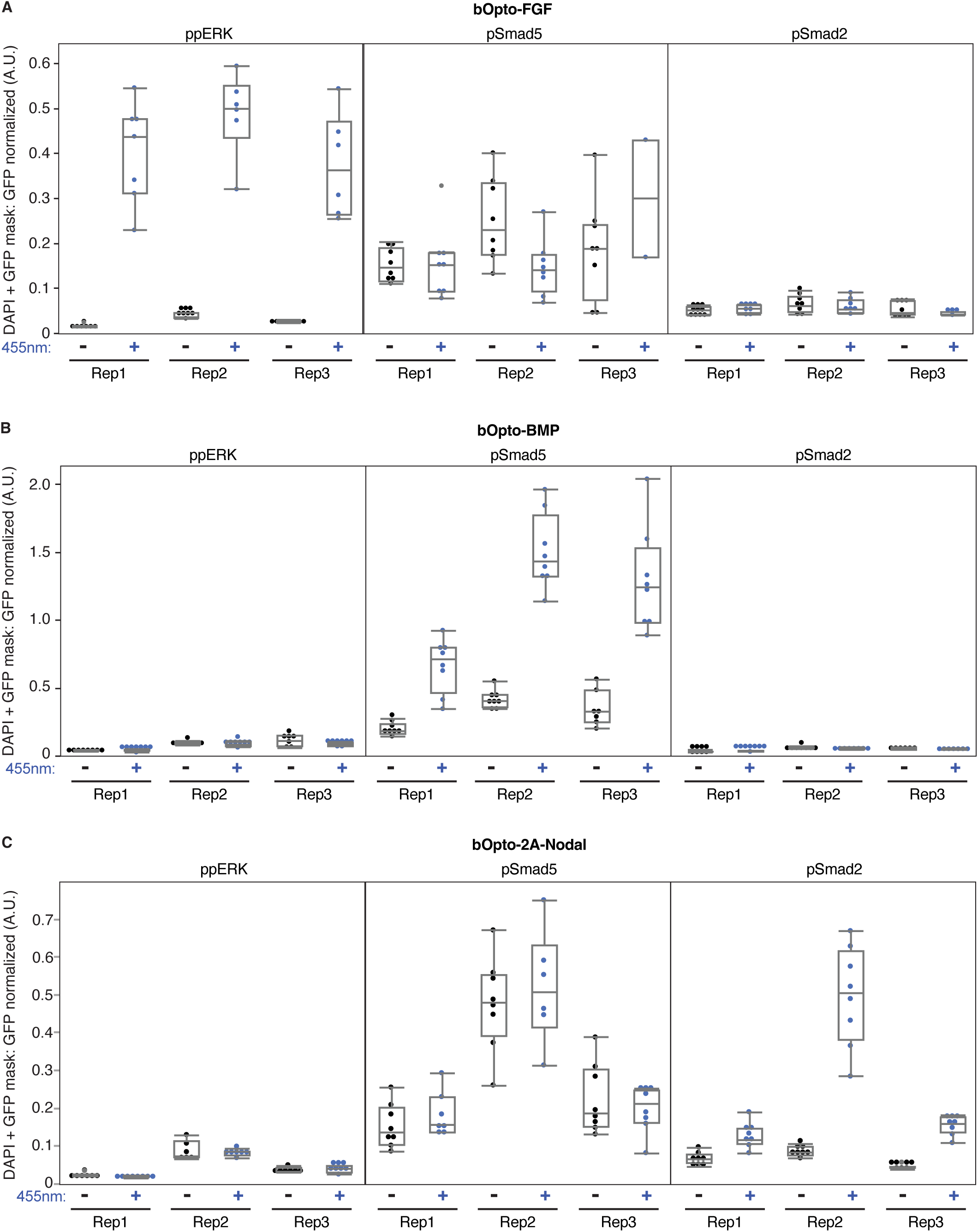
Quantification of pathway specificity experiments. Embryos injected at the one-cell stage with mRNA encoding *GFP* & *bOpto-FGF* **(A)**, *bOpto-BMP* **(B)**, or *bOpto-2A-Nodal* **(C)** were exposed to dark or 455 nm light (50 W/m^2^) for 30 minutes starting at gastrulation (50% epiboly - shield). HCR-IF was used to detect phosphorylated signaling effectors (ppERK1/2, pSmad1/5/9, & pSmad2/3 reflect FGF, BMP, & Nodal signaling, respectively). Raw phosphorylated signaling effector intensity was measured in each DAPI + GFP-positive pixel and was divided by the corresponding GFP intensity. Each dot represents the median GFP-normalized nuclear pixel signal in one embryo.

**Supplementary Figure 7:**
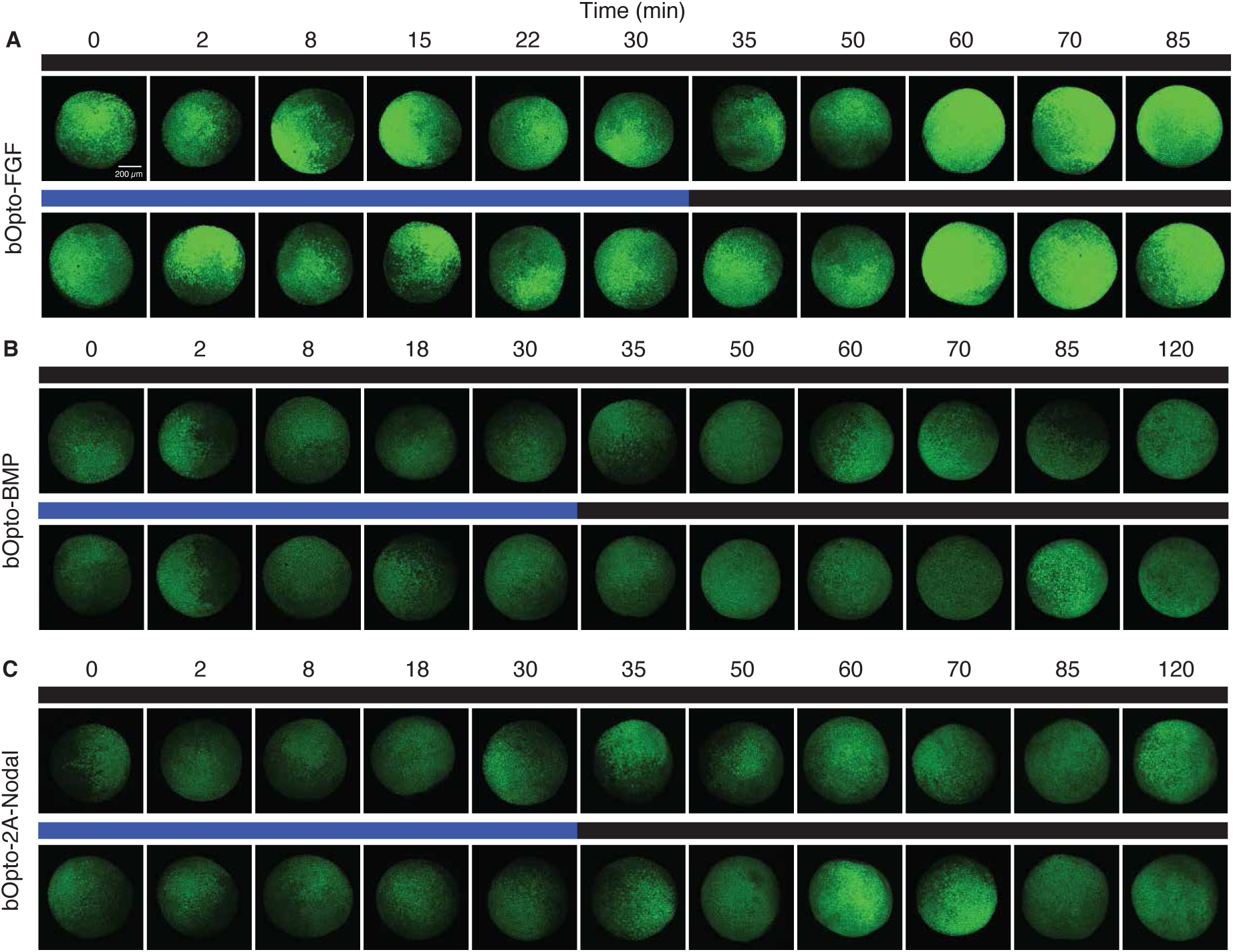
Co-injected GFP signal used for quantification of on / off kinetics. **A-C)** Embryos were injected at the one-cell stage with mRNA encoding *GFP* and either *bOp-to-FGF* (A), *bOpto-BMP* (B), or *bOpto-2A-Nodal* (C). Starting at early gastrulation (50% epiboly - shield), embryos were exposed to 455 nm light (50 W/m^2^) for 30 min and fixed at different time points during and after exposure. HCR-IF staining used to detect activated signaling effectors is shown in Fig. 6. HCR-IF signal was normalized voxel-wise based on co-injected GFP intensity shown here (see Supp. Fig. 8). GFP images correspond to representative HCR-IF images shown in Fig. 6. Scale bar is 200 µm.

**Supplementary Figure 8:**
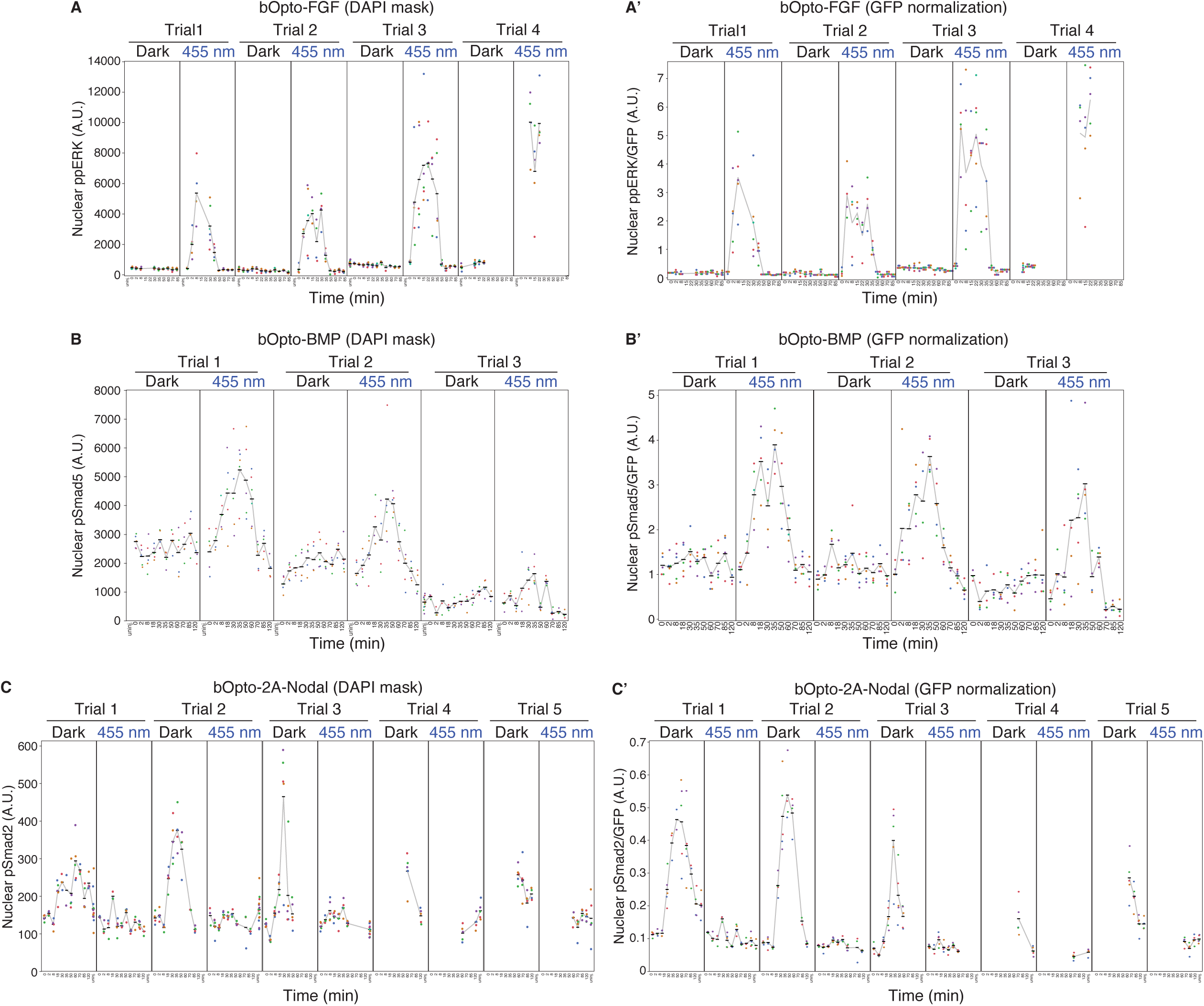
Quantification of optogenetic signaling activator toolkit on/off kinetics. Embryos were injected at the one-cell stage with mRNA encoding *GFP* and either *bOpto-FGF* (A,A’), *bOpto-BMP* (B,B’), or *bOpto-2A-Nodal* (C,C’). Starting at early gastrulation (50% epiboly - shield), embryos were exposed to 455 nm light (50 W/m^2^) for 30 min and fixed during and after exposure. Three full repeats were performed across 3-5 trials. HCR-IF staining used to detect phosphorylated signaling effectors is shown in Fig. 6 (ppERK1/2, pSmad1/5/9, and pSmad2/3 reflect FGF, BMP, and Nodal signaling, respectively). **A,B,C)** Raw phosphorylated signaling effector intensity was measured in each DAPI-positive nuclear pixel. Each dot represents the median nuclear pixel intensity in one embryo. Black lines represent the mean nuclear intensity of all embryos in the indicated condition. **A’,B’, C’)** Phosphorylated effector intensity in each DAPI + GFP-positive pixel was divided by the corresponding GFP intensity. Each dot represents the median GFP-normalized nuclear pixel signal in one embryo. Black lines represent the mean GFP-normalized nuclear signal of all embryos in the indicated condition.

**Supplementary Figure 9:**
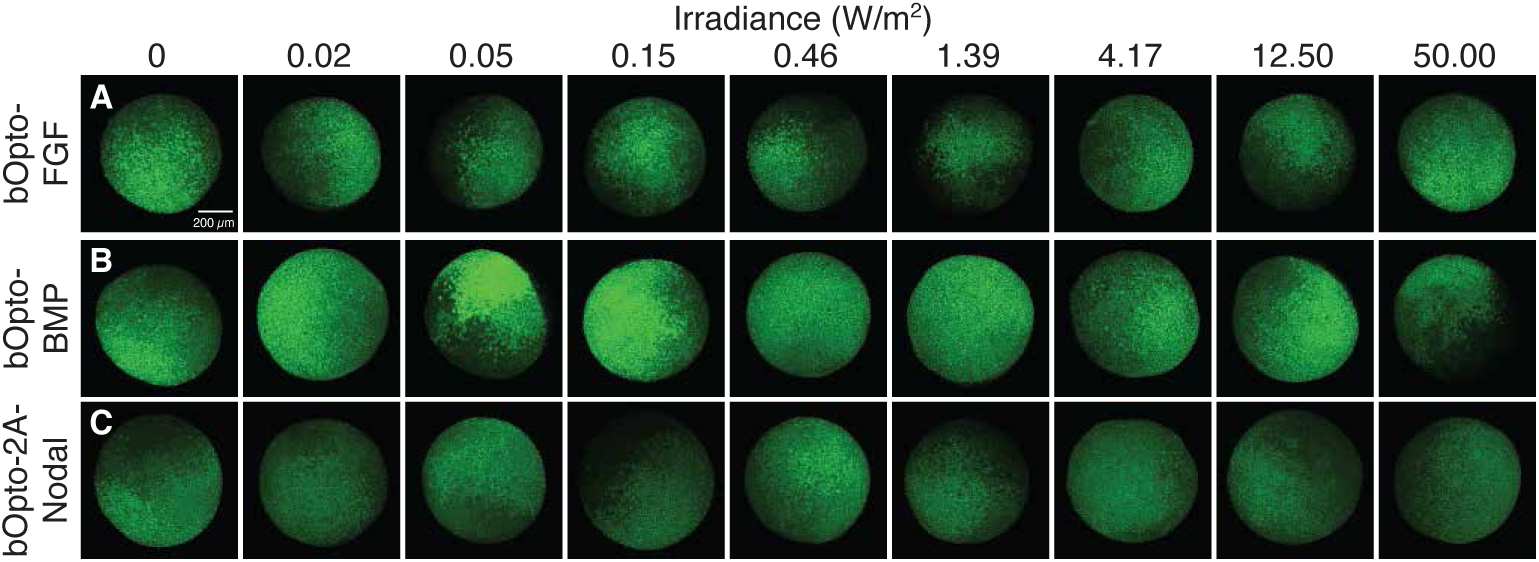
Co-injected GFP signal used for quantification of irradiance dependence HCR-IF. **A-C)** Embryos were injected at the one-cell stage with mRNA encoding *GFP* and either *bOpto-FGF* (A), *bOpto-BMP* (B), or *bOp-to-2A-Nodal* (C). Starting at early gastrulation (50% epiboly - shield), embryos were exposed to 455 nm light at the indicated irradiances for 5 min (bOpto-FGF) or 25 min (bOpto-BMP and -2A-Nodal). HCR-IF staining used to detect phosphorylated signaling effectors is shown in Fig. 7. HCR-IF signal was normalized based on co-injected GFP intensity shown here; GFP images correspond to representative images shown in Fig. 7. Scale bar is 200 µm.

**Supplementary Figure 10:**
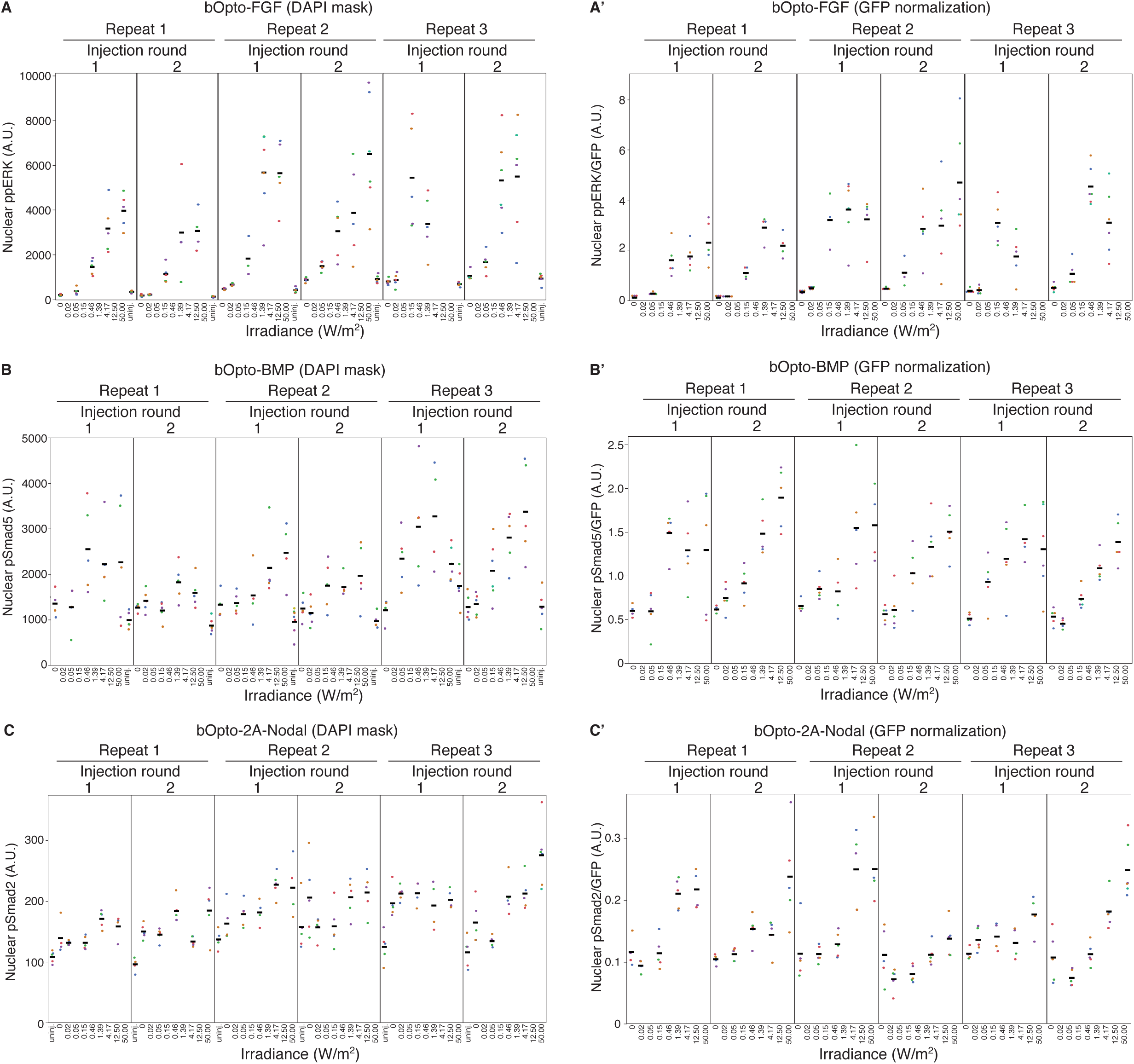
Quantification of light irradiance dependence. Embryos were injected at the one-cell stage with mRNA encoding *GFP* and either *bOpto-FGF* (A,A’), *bOpto-BMP* (B,B’), or *bOpto-2A-Nodal* (C,C’). At early gastrulation (50% epiboly - shield), embryos were exposed to 455 nm light at the indicated irradiance for 5 min (bOpto-FGF) or 25 min (bOpto-BMP and -2A-Nodal). HCR-IF was used to detect phosphorylated signaling effectors (ppERK1/2, pSmad1/5/9, & pSmad2/3 reflect FGF, BMP, & Nodal signaling, respectively). **A,B,C)** Raw phosphorylated signaling effector intensity was measured in each DAPI-positive nuclear pixel. Each dot represents the median nuclear pixel intensity in one embryo. Black lines represent the mean nuclear intensity of all embryos in the indicated condition. **A’,B’, C’)** Phosphorylated effector intensity in each DAPI + GFP-positive pixel was divided by the corresponding GFP intensity. Each dot represents the median GFP-normalized nuclear pixel signal in one embryo. Black lines represent the mean GFP-normalized nuclear signal of all embryos in the indicated condition.

**Supplementary Figure 11:**
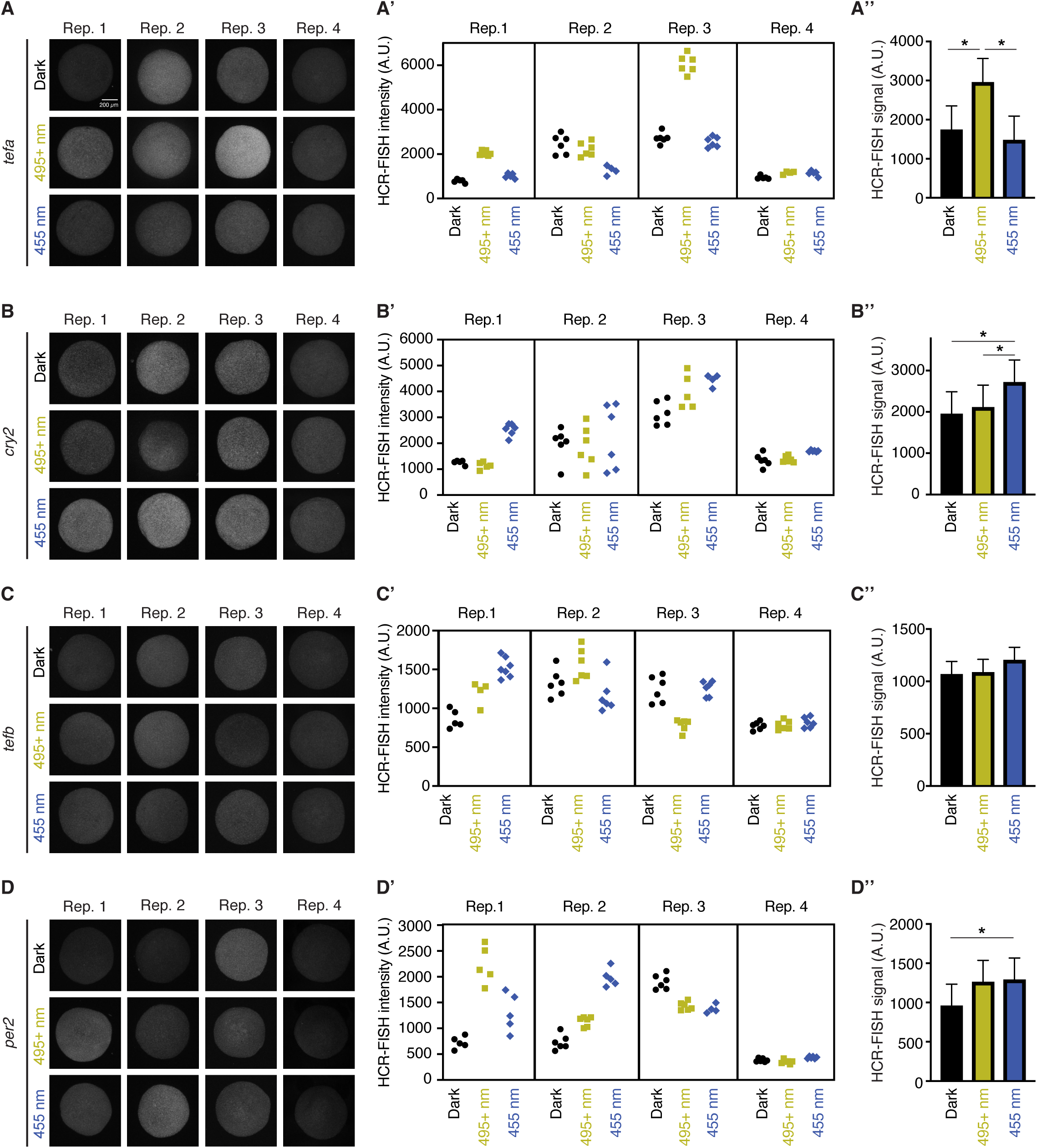
Circadian gene responses to light exposure in gasturation stage embryos. **A-D)** Uninjected AB wild type zebrafish embryos were reared in the dark. At early gastrulation (shield stage), a subset of embryos were exposed to 495+ nm light (18.51 W/m^2^) or 455 nm light (50 W/m^2^) for 30 min, then dark for 20 min. A third set of embryos were maintained in the dark throughout the experiment. HCR-FISH was performed for the indicated circadian-related genes. Each panel represents four experimental repeats for the indicated gene. Scale bar is 200 µm. **A’-D’)** Mean HCR-FISH intensities from experiments shown in A-D. Each symbol represents mean intensity from one embryo. **A’’-D’’)** Quantification of experiments shown in A-D. Linear-mixed model-predicted least squared means of HCR-FISH signal +/- SEM (N = 4; LMM - Fixed Effect: Wavelength, Random Effect: Replicate; post hoc Tukey’s HSD where * indicates p < 0.05).

