## Supplementary material for "An optogenetic toolkit for robust activation of FGF, BMP, & Nodal signaling in zebrafish": Pathway_Specific_Activation_Macro.ijm

```
//Fiji based macro used for analyzing pathway specific optogenetic activation of FGF, BMP, and Nodal signaling	
//The following Fiji based macro was used to generate maximum intensity projections (MIPs) with the above give “Min”, “Max” values for each pathway specific signaling effector. 

rename("A");
run("Split Channels");
selectWindow("C1-A");
run("Z Project...", "projection=[Max Intensity]");
run("Grays");
rename("DAPI");
setMinAndMax(min,max);
run("Out [-]");
run("RGB Color");
selectWindow("C1-A");
close();
selectWindow("C2-A");
run("Z Project...", "projection=[Max Intensity]");
run("Yellow");
rename("pSmad2");
setMinAndMax(min,max);
run("RGB Color");
run("Out [-]");
selectWindow("C2-A");
close();
selectWindow("C3-A");
run("Z Project...", "projection=[Max Intensity]");
run("Cyan");
rename("ppERK");
setMinAndMax(min,max);
run("RGB Color");
run("Out [-]");
selectWindow("C3-A");
close();
selectWindow("C4-A");
run("Z Project...", "projection=[Max Intensity]");
run("Magenta");
rename("pSmad1");
setMinAndMax(min,max);
run("RGB Color");
run("Out [-]");
selectWindow("C4-A");
close();
run("Merge Channels...", "c5=pSmad2 c6=ppERK c7=pSmad1 keep");
```
