## Supplementary material for "An optogenetic toolkit for robust activation of FGF, BMP, & Nodal signaling in zebrafish": Iannucci_Maria_Thomas_Supp_Materials.pdf

**SUPPLEMENTARY MATERIALS**

**Supplementary Schematics**

Supplementary Schematic 1: On/off kinetics of optogenetic signaling activator toolkit I.

**Supplementary Tables**

Supplementary Table 1: Summary of Imaging Parameters.

Supplementary Table 2: Wavelength-dependent activation of FGF, BMP, and Nodal signaling I.

Supplementary Table 3: Wavelength-dependent activation of FGF, BMP, and Nodal signaling II.

Supplementary Table 4: Wavelength-dependent activation of FGF, BMP, and Nodal signaling III.

Supplementary Table 5: Pathway specific optogenetic activation of FGF, BMP, and Nodal signaling.

Supplementary Table 6: On/off kinetics of optogenetic signaling activator toolkit I.

Supplementary Table 7: On/off kinetics of optogenetic signaling activator toolkit II.

Supplementary Table 8: On/off kinetics of optogenetic signaling activator toolkit III.

Supplementary Table 9: On/off kinetics of optogenetic signaling activator toolkit IV.

Supplementary Table 10: Irradiance sensitivity of optogenetic toolkit I.

Supplementary Table 11: List of pathway-specific target genes and corresponding HCR initiators and amplifiers.

Supplementary Table 12: List of pathway-specific target genes and corresponding HCR probes.

**Supplementary Information**

Supplementary Information 1: Construct sequences.

**References**

**Supplementary Schematic 1: On/off kinetics of optogenetic signaling activator toolkit I.**
Visualization of tON & tOFF parameters to describe on/off kinetics in Fig. 6.

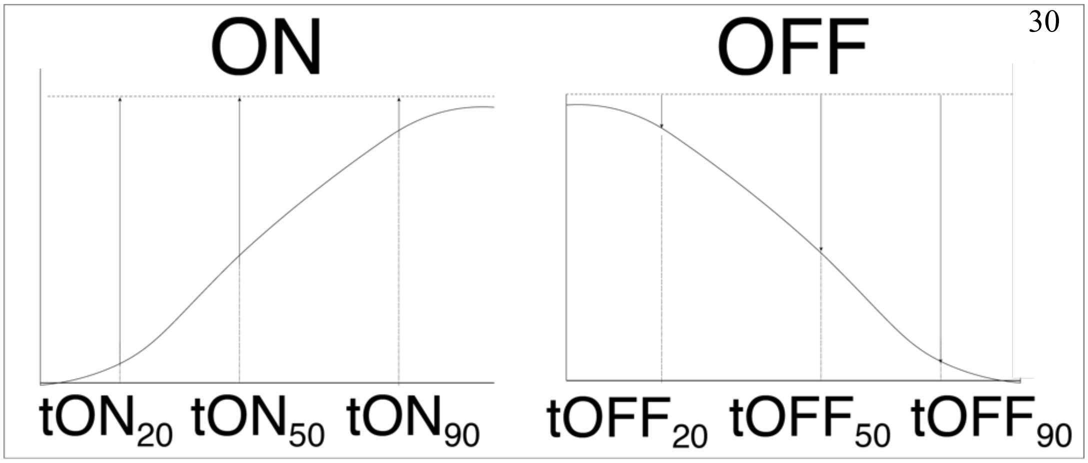

**Supplementary Table 1: Summary of Imaging Parameters.** Fluorophores used in HCR-IF, HCR-FISH and fluorescent proteins detected with corresponding imaging conditions.

| **Fluorophore/Protein** | **Fluorophore/Fluorescent protein** | **Laser Line (nm)** | **Emission wavelength (nm)** | **Signal** | **Experiments** |
|---|---|---|---|---|---|
| Alexa-647 | Fluorophore | 640 | 553-700 | pSmad5, pSmad2, ppERK1/2, *sizzled, bambia, spry4, il17rd, noto, lft1* | Time Course, Intensity Dependent, Wavelength, Gene Expression, Spatial Activation, Pathway Specificity |
| Alexa-546 | Fluorophore | 561 | 553-700 | pSmad2, *gata2a, dusp6, gsc* | Intensity Dependent, Gene Expression |
| Kaede-Red | Fluorescent protein | 561 | 576-616 | Nuclear-Kaede-Red | Spatial Activation |
| Alexa-488 | Fluorophore | 488 | 480-560 | *sizzled, bambia, spry4, il17rd, noto, lft1* | Gene Expression |
| GFP | Fluorescent protein | 488 | 480-560 | GFP | Time Course, Intensity Dependence, Wavelength |
| Kaede-Green | Fluorescent protein | 488 | 410-546 | Nuclear-Kaede-Green | Spatial Activation |
| DAPI | Fluorophore | 405 | 400-489 | Nuclei | Used in all experiments except Spatial experiment |

**Supplementary Table 2: Wavelength-dependent activation of FGF, BMP, and Nodal signaling I.** Fixed effects tests from Linear Mixed-effects Model (LMM) shown in Fig. 3. Fixed Effect: Wavelength, Random Effect: Biological Replicate; Personality: Standard Least Squares, Method: REML. $p < 0.05$ considered to be significant.

| | **Source** | **Nparm** | **DF** | **DFDen** | **F Ratio** | **Prob > F** |
|---|---|---|---|---|---|---|
| **FGF** | Exposure | 2 | 2 | 65.01 | 183.68 | <.0001 |
| **BMP** | Exposure | 2 | 2 | 41.05 | 35.78 | <.0001 |
| **Nodal** | Exposure | 2 | 2 | 65.08 | 188.71 | <.0001 |

**Supplementary Table 3: Wavelength-dependent activation of FGF, BMP, and Nodal signaling II.** Least square means estimates from Linear Mixed-effects Model (LMM) shown in Fig. 3 and Supp. Table 2. Fixed Effect: Wavelength, Random Effect: Biological Replicate; Personality: Standard Least Squares, Method: REML.

| | Exposure | Estimate | Std Error | DF | Lower 95% | Upper 95% |
|---|---|---|---|---|---|---|
| **FGF** | Dark | 0.24 | 0.08 | 2.39 | -0.05 | 0.53 |
| | Yellow | 0.19 | 0.08 | 2.33 | -0.10 | 0.49 |
| | Blue | 0.81 | 0.08 | 2.27 | 0.52 | 1.11 |
| **BMP** | Dark | 0.72 | 0.22 | 2.24 | -0.14 | 1.60 |
| | Yellow | 0.74 | 0.22 | 2.24 | -0.11 | 1.59 |
| | Blue | 1.30 | 0.22 | 2.15 | 0.43 | 2.17 |
| **Nodal** | Dark | 0.09 | 0.02 | 10.26 | 0.05 | 0.12 |
| | Yellow | 0.09 | 0.02 | 9.57 | 0.05 | 0.12 |
| | Blue | 0.43 | 0.02 | 8.31 | 0.39 | 0.46 |

**Supplementary Table 4: Wavelength-dependent activation of FGF, BMP, and Nodal signaling III.** Results of *post hoc* pairwise comparisons of Least square means and standard errors estimated in Supp. Table 2-3. Comparisons were performed with Tukey's HSD test where $p < 0.05$ is considered to be significant and denoted with * in Fig. 3.

| | Exposure | Minus Exposure | Difference | Std Error | t Ratio | Prob>\|t\| | Lower 95% | Upper 95% |
|---|---|---|---|---|---|---|---|---|
| **FGF** | Dark | Yellow | 0.04 | 0.04 | 1.17 | 0.47 | -0.05 | 0.14 |
| | Blue | Dark | 0.58 | 0.04 | 15.48 | <.0001 | 0.49 | 0.66 |
| | Blue | Yellow | 0.62 | 0.04 | 17.14 | <.0001 | 0.53 | 0.71 |
| **BMP** | Dark | Yellow | -0.02 | 0.09 | -0.28 | 0.96 | -0.23 | 0.19 |
| | Blue | Dark | 0.59 | 0.08 | 7.32 | <.0001 | 0.39 | 0.78 |
| | Blue | Yellow | 0.56 | 0.08 | 6.97 | <.0001 | 0.37 | 0.76 |
| **Nodal** | Dark | Yellow | 0.00 | 0.02 | 0.02 | 1.00 | -0.05 | 0.05 |
| | Blue | Dark | 0.34 | 0.02 | 16.56 | <.0001 | 0.29 | 0.39 |
| | Blue | Yellow | 0.34 | 0.02 | 16.82 | <.0001 | 0.29 | 0.39 |

**Supplementary Table 5: Pathway-specific optogenetic activation of FGF, BMP, and Nodal signaling.**

| **bOpto-FGF** | | | | | |
|---|---|---|---|---|---|
| **Fixed Effect Test** | | | | | |
| **Source** | **Nparm** | **DF** | **DFDen** | **F Ratio** | **Prob > F** |
| Pathway | 2 | 2 | 121.5 | 71.372 | <.0001* |
| Light | 1 | 1 | 122.3 | 97.5002 | <.0001* |
| Pathway*Light | 2 | 2 | 121.4 | 125.9479 | <.0001* |
| **LS Means Estimates** | | | | | |
| **Level** | **Least Sq Mean** | **Std Error** | | | |
| ppERK,Blue | 0.42062635 | 0.01771607 | | | |
| ppERK,Dark | 0.02685416 | 0.01647561 | | | |
| pSmad5,Blue | 0.16655686 | 0.01840752 | | | |
| pSmad5,Dark | 0.19642751 | 0.01621398 | | | |
| pSmad2,Blue | 0.05416893 | 0.01781414 | | | |
| pSmad2,Dark | 0.05594617 | 0.01576753 | | | |
| **Post hoc Contrast Testing** | | | | | |
| ppERK,Blue | 1 | 0 | 0 | | |
| ppERK,Dark | -1 | 0 | 0 | | |
| pSmad5,Blue | 0 | 1 | 0 | | |
| pSmad5,Dark | 0 | -1 | 0 | | |
| pSmad2,Blue | 0 | 0 | 1 | | |
| pSmad2,Dark | 0 | 0 | -1 | | |
| Estimate | 0.3938 | -0.03 | -0.002 | | |
| Std Error | 0.0211 | 0.0215 | 0.0206 | | |
| t Ratio | 18.659 | -1.391 | -0.086 | | |
| Prob>\|t\| | 2.00E-37 | 0.1668 | 0.9313 | | |
| SS | . | . | . | | |
| Lower 95% | 0.352 | -0.072 | -0.043 | | |
| Upper 95% | 0.4356 | 0.0126 | 0.039 | | |
| **Bonferroni-adjusted P value** | 0.0000 | 0.5004 | 2.7939 | | |
| Paper formatted p-value | < 0.0001 | 0.5004 | >0.9999 | | |
| Fold Change | 15.7 | | | | |

| | | | | | |
|---|---|---|---|---|---|
| **bOpto-BMP** | | | | | |
| **Fixed Effect Test** | | | | | |
| **Source** | **Nparm** | **DF** | **DFDen** | **F Ratio** | **Prob > F** |
| Pathway | 2 | 2 | 131 | 220.0304 | <.0001* |
| Light | 1 | 1 | 131 | 80.7804 | <.0001* |
| Pathway*Light | 2 | 2 | 131 | 85.5949 | <.0001* |
| **LS Means Estimates** | | | | | |
| **Level** | **Least Sq Mean** | **Std Error** | | | |
| ppERK,Blue | 0.0744056 | 0.06934978 | | | |
| ppERK,Dark | 0.0841817 | 0.0697715 | | | |
| pSmad5,Blue | 1.1528356 | 0.06934978 | | | |
| pSmad5,Dark | 0.3221164 | 0.0697715 | | | |
| pSmad2,Blue | 0.0509066 | 0.0697715 | | | |
| pSmad2,Dark | 0.0546684 | 0.07024428 | | | |
| **Post hoc Contrast Testing** | | | | | |
| ppERK,Blue | 1 | 0 | 0 | | |
| ppERK,Dark | -1 | 0 | 0 | | |
| pSmad5,Blue | 0 | 1 | 0 | | |
| pSmad5,Dark | 0 | -1 | 0 | | |
| pSmad2,Blue | 0 | 0 | 1 | | |
| pSmad2,Dark | 0 | 0 | -1 | | |
| Estimate | -0.01 | 0.8307 | -0.004 | | |
| Std Error | 0.0521 | 0.0521 | 0.0532 | | |
| t Ratio | -0.188 | 15.948 | -0.071 | | |
| Prob>\|t\| | 0.8514 | 2.00E-32 | 0.9438 | | |
| SS | . | . | . | | |
| Lower 95% | -0.113 | 0.7277 | -0.109 | | |
| Upper 95% | 0.0933 | 0.9338 | 0.1016 | | |
| **Bonferroni-adjusted P value** | 2.5542 | 0.0000 | 2.8314 | | |
| Paper formatted | >0.9999 | <0.0001 | >0.9999 | | |
| Fold Change | | 3.6 | | | |
| **bOpto-Nodal** | | | | | |

| **Fixed Effect Test** | | | | | |
|---|---|---|---|---|---|
| **Source** | **Nparm** | **DF** | **DFDen** | **F Ratio** | **Prob > F** |
| Pathway | 2 | 2 | 128 | 72.682 | <.0001* |
| Light | 1 | 1 | 128 | 17.1768 | <.0001* |
| Pathway*Light | 2 | 2 | 128 | 16.4851 | <.0001* |
| **LS Means Estimates** | | | | | |
| **Level** | **Least Sq Mean** | **Std Error** | | | |
| ppERK,Blue | 0.0489442 | 0.0649109 | | | |
| ppERK,Dark | 0.0524716 | 0.0653385 | | | |
| pSmad5,Blue | 0.292529 | 0.0651871 | | | |
| pSmad5,Dark | 0.2842524 | 0.0647898 | | | |
| pSmad2,Blue | 0.258035 | 0.0647898 | | | |
| pSmad2,Dark | 0.0653179 | 0.0647898 | | | |
| **Post hoc Contrast Testing** | | | | | |
| ppERK,Blue | 1 | 0 | 0 | | |
| ppERK,Dark | -1 | 0 | 0 | | |
| pSmad5,Blue | 0 | 1 | 0 | | |
| pSmad5,Dark | 0 | -1 | 0 | | |
| pSmad2,Blue | 0 | 0 | 1 | | |
| pSmad2,Dark | 0 | 0 | -1 | | |
| Estimate | -0.004 | 0.0083 | 0.1927 | | |
| Std Error | 0.0282 | 0.0276 | 0.0267 | | |
| t Ratio | -0.125 | 0.2998 | 7.2291 | | |
| Prob>\|t\| | 0.9008 | 0.7648 | 4.00E-11 | | |
| SS | . | . | . | | |
| Lower 95% | -0.059 | -0.046 | 0.14 | | |
| Upper 95% | 0.0523 | 0.0629 | 0.2455 | | |
| **Bonferroni-adjusted P value** | 2.7024 | 2.2944 | 0.0000 | | |
| Paper Formatted | >0.9999 | >0.9999 | <0.0001 | | |
| Fold Change | | | 4.0 | | |

**Supplementary Table 6: On/off kinetics of optogenetic signaling activator toolkit I.** Fixed effects tests from One-way ANOVA shown in Fig. 6. Fixed Effect: Time. $p < 0.05$ was considered to be significant.

| Pathway | Source | DF | Sum of Squares | Mean Square | F Ratio | Prob > F |
|---|---|---|---|---|---|---|
| **FGF** | Time | 10 | 89.31 | 8.93 | 7.06 | <.0001* |
| | Error | 23 | 29.11 | 1.27 | | |
| | C. Total | 33 | 118.42 | | | |
| **BMP** | Time | 10 | 25.62 | 2.56 | 13.75 | <.0001* |
| | Error | 22 | 4.10 | 0.19 | | |
| | C. Total | 32 | 29.72 | | | |
| **Nodal** | Time | 10 | 0.51 | 0.05 | 7.84 | <.0001* |
| | Error | 22 | 0.14 | 0.01 | | |
| | C. Total | 32 | 0.65 | | | |

**Supplementary Table 7: On/off kinetics of optogenetic signaling activator toolkit II.** LSD Threshold Matrix of pairwise *post hoc* comparisons to time = 0 min (Fig. 6 and Supp. Table 6). Comparisons were performed with Dunnett's Method for multiple comparison where $p < 0.05$ is considered to be significant and denoted with * in Fig. 6.

| FGF | | | BMP | | | Nodal | | |
|---|---|---|---|---|---|---|---|---|
| **Time** | **Abs(Dif)-LSD** | **p-Value** | **Time** | **Abs(Dif)-LSD** | **p-Value** | **Time** | **Abs(Dif)-LSD** | **p-Value** |
| 0 | -2.70 | 1 | 0 | -1.04 | 1 | 0 | -0.19 | 1 |
| 2 | 0.52 | 0.0143* | 2 | -0.22 | 0.1713 | 2 | -0.19 | 1 |
| 8 | 0.75 | 0.0070* | 8 | -0.15 | 0.1153 | 8 | -0.19 | 1 |
| 15 | 0.77 | 0.0075* | 18 | 0.92 | 0.0001* | 18 | -0.08 | 0.4192 |
| 22 | 1.24 | 0.0022* | 30 | 0.50 | 0.0020* | 30 | 0.15 | 0.0003* |
| 30 | -0.11 | 0.0647 | 35 | 1.47 | <.0001* | 35 | 0.13 | 0.0005* |
| 35 | -1.17 | 0.4953 | 50 | 0.30 | 0.0076* | 50 | 0.09 | 0.0023* |
| 50 | -2.67 | 1 | 60 | -0.20 | 0.154 | 60 | 0.01 | 0.0358* |
| 60 | -2.59 | 1 | 70 | -0.94 | 1 | 70 | -0.04 | 0.1815 |
| 70 | -2.67 | 1 | 85 | -0.80 | 0.9917 | 85 | -0.13 | 0.94 |
| 85 | -2.67 | 1 | 120 | -0.89 | 0.9997 | 120 | -0.14 | 0.9539 |

**Supplementary Table 8: On/off kinetics of optogenetic signaling activator toolkit III.**

Parameters predicted with 95% CI from non-linear, three-parameter logistic regression of data in Fig. 6A’’-C’’; Data from Fig.6A’-C’ & Supp. Fig. where time <= 30 min.

| Pathway | Phase | | R2 | Growth Rate | Inflection Point | Asymptote | tON20 | tON50 | tON90 |
|---|---|---|---|---|---|---|---|---|---|
| **FGF** | **ON** | Predictive Y | | | | | 0.67 | 1.68 | 3.08 |
| | | Estimate | 0.490 | 4.05 | 1.14 | 3.35 | 0.79 | 1.14 | 1.68 |
| | | Lower 95% CI | | -20.58 | -4.36 | 2.62 | -6.27 | -4.33 | -2.20 |
| | | Upper 95% CI | | 28.68 | 6.63 | 4.08 | 7.86 | 6.61 | 5.57 |
| **BMP** | **ON** | Predictive Y | | | | | 0.32 | 0.79 | 1.43 |
| | | Estimate | 0.643 | 0.30 | 7.34 | 1.59 | 2.76 | 7.31 | 14.66 |
| | | Lower 95% CI | | -0.09 | 2.23 | 1.12 | -2.94 | 2.99 | 3.42 |
| | | Upper 95% CI | | 0.69 | 12.45 | 2.06 | 8.47 | 11.62 | 25.91 |
| **Nodal** | **ON** | Predictive Y | | | | | 0.07 | 0.17 | 0.31 |
| | | Estimate | 0.966 | 0.38 | 19.78 | 0.35 | 16.12 | 19.78 | 25.59 |
| | | Lower 95% CI | | -0.41 | 15.08 | 0.27 | 12.25 | 16.13 | 14.04 |
| | | Upper 95% CI | | 1.16 | 24.48 | 0.42 | 19.99 | 23.43 | 37.14 |

**Supplementary Table 9: On/off kinetics of optogenetic signaling activator toolkit IV.**

Parameters predicted with 95% CI from non-linear, three-parameter logistic regression of data in Fig. 6A’’’-C’’’; Data from Fig. 6A’-C’ where time >= 30 min.

| Pathway | Phase | | R2 | Growth Rate | Inflection Point | Asymptote | tOFF20 | tOFF50 | tOFF90 |
|---|---|---|---|---|---|---|---|---|---|
| **FGF** | **OFF** | Predictive Y | | | | | 2.26 | 1.41 | 0.28 |
| | | Estimate | 0.791 | -0.47 | 35.45 | 2.82 | 2.51 | 5.45 | 10.10 |
| | | Lower 95% CI | | -16.60 | 18.99 | -16.10 | -42.62 | -6.77 | -147.52 |
| | | Upper 95% CI | | 15.65 | 51.92 | 21.74 | 47.70 | 17.68 | 167.72 |
| **BMP** | **OFF** | Predictive Y | | | | | 1.47 | 0.92 | 0.18 |
| | | Estimate | 0.761 | -0.20 | 54.68 | 1.84 | 17.91 | 24.69 | 35.63 |
| | | Lower 95% CI | | -0.44 | 47.69 | 1.35 | 8.46 | 18.71 | 22.54 |
| | | Upper 95% CI | | 0.03 | 61.68 | 2.33 | 27.36 | 30.67 | 48.72 |
| **Nodal** | **OFF** | Predictive Y | | | | | 0.31 | 0.19 | 0.04 |
| | | Estimate | 0.626 | -0.06 | 62.67 | 0.39 | 10.63 | 32.67 | 67.60 |
| | | Lower 95% CI | | -0.14 | 36.80 | 0.15 | -4.34 | 22.22 | 33.56 |
| | | Upper 95% CI | | 0.01 | 88.54 | 0.62 | 25.61 | 43.12 | 101.63 |

**Supplementary Table 10: Irradiance sensitivity of optogenetic toolkit I.** Parameters predicted with 95% CI from non-linear, three-parameter logistic regression of data in Fig. 7B'-D'. Dosage parameters (D20 & D90) are calculated from corresponding Irradiance parameters (I20, I90) by multiplying by time of exposure in seconds for each tool (FGF: 5 min, BMP/Nodal: 25 min). Fold increase is a ratio between the D20 & D90 of each tool & FGF's D20 and D90, respectively.

| **Pathway** | | **R2** | **Growth Rate** | **Inflection Point** | **Asymptote** | | | | **Dosage** | | **Fold Increase over FGF** | |
|---|---|---|---|---|---|---|---|---|---|---|---|---|
| | | | | | | **I20** | **I50** | **I90** | **D20** | **D90** | **D20** | **D90** |
| **FGF** | Predictive Y | | | | | 0.52 | 1.29 | 2.33 | | | | |
| | Estimate | 0.654 | 34.49 | 0.10 | 2.59 | 0.06 | 0.10 | 0.16 | 17.61 | 29.58 | 1.00 | 1.00 |
| | Lower 95% CI | | -1.30 | 0.04 | 2.15 | 0.00 | 0.04 | 0.07 | 0.00 | 13.24 | | |
| | Upper 95% CI | | 70.27 | 0.16 | 3.02 | 0.12 | 0.15 | 0.25 | 36.04 | 45.92 | | |
| **BMP** | Predictive Y | | | | | 0.17 | 0.42 | 0.76 | | | | |
| | Estimate | 0.736 | 5.58 | 0.30 | 0.85 | 0.05 | 0.30 | 0.69 | 80.35 | 1032.58 | 5 | 35 |
| | Lower 95% CI | | 1.40 | 0.14 | 0.73 | -0.12 | 0.15 | 0.30 | -182.09 | 445.85 | | |
| | Upper 95% CI | | 9.76 | 0.46 | 0.96 | 0.23 | 0.44 | 1.08 | 342.79 | 1619.30 | | |
| **Nodal** | Predictive Y | | | | | 0.03 | 0.07 | 0.12 | | | | |
| | Estimate | 0.578 | 0.20 | 11.00 | 0.13 | 3.90 | 11.00 | 22.25 | 5845.05 | 33376.65 | 332 | 1128 |
| | Lower 95% CI | | 0.05 | 4.57 | 0.09 | -0.27 | 5.54 | 6.12 | -406.04 | 9182.01 | | |
| | Upper 95% CI | | 0.34 | 17.42 | 0.18 | 8.06 | 16.45 | 38.38 | 12096.15 | 57571.22 | | |

**Supplementary Table 11: List of pathway-specific target genes and corresponding HCR initiators and amplifiers.** Used to generate images in Fig. 4.

| Pathway | Gene | Initiator | Amplifier |
|---|---|---|---|
| BMP | *bambia* | B1 | 647 |
| | *gata2a* | B2 | 546 |
| | *szl* | B3 | 488 |
| FGF | *spry4* | B1 | 647 |
| | *dusp6* | B2 | 546 |
| | *Il17rd* | B3 | 488 |
| Nodal | *noto* | B1 | 647 |
| | *gsc* | B2 | 546 |
| | *lft1* | B3 | 488 |

**Supplementary Table 12: List of pathway-specific target genes and corresponding HCR probes.** Used to generate images in Fig. 4.

| Sequence name | Sequence |
|---|---|
| B1_bambia_26_Dla50 | GAGGAGGGCAGCAAACGGaaCATAAGCTCCACCCTTTTCATATTC |
| B1_bambia_26_Dla50 | AGAGGGAAGGAGGTGGGGCTTGCAGtaGAAGAGTCTTCCTTTACG |
| B1_bambia_26_Dla50 | GAGGAGGGCAGCAAACGGaaCCTTGAAATGTCTTTGCTCTCGTTG |
| B1_bambia_26_Dla50 | TTTTTGAGAAAACACAATGAATTGTtaGAAGAGTCTTCCTTTACG |
| B1_bambia_26_Dla50 | GAGGAGGGCAGCAAACGGaaAACATTACGACAGCAAACTACACAG |
| B1_bambia_26_Dla50 | ACAATAAAATAACACGCCGATAATAtaGAAGAGTCTTCCTTTACG |
| B1_bambia_26_Dla50 | GAGGAGGGCAGCAAACGGaaTGAGTAGTCATACGAACTCCAGCTT |
| B1_bambia_26_Dla50 | CCCTTTTGTGCCCTAACAAAGAAGCtaGAAGAGTCTTCCTTTACG |
| B1_bambia_26_Dla50 | GAGGAGGGCAGCAAACGGaaCAGAGATAGGAGACGCTCACCCCCG |
| B1_bambia_26_Dla50 | GTGCCCCGTGTACATCCCCCAGTGTtaGAAGAGTCTTCCTTTACG |
| B1_bambia_26_Dla50 | GAGGAGGGCAGCAAACGGaaCGCAGCTTATCGCAGCCCAGACAAC |
| B1_bambia_26_Dla50 | CCTCCTCCAGTGCACAAATCCGTCTtaGAAGAGTCTTCCTTTACG |
| B1_bambia_26_Dla50 | GAGGAGGGCAGCAAACGGaaACTCCAAGTCCAACTTAGCCACGTG |
| B1_bambia_26_Dla50 | TCTCATGTCCCGTTACCGGCACCATtaGAAGAGTCTTCCTTTACG |
| B1_bambia_26_Dla50 | GAGGAGGGCAGCAAACGGaaGCTGTAATGCAGGCGAGAAAGCATC |
| B1_bambia_26_Dla50 | TTTCTTGGCATGGTGGTGTCCGTGAtaGAAGAGTCTTCCTTTACG |
| B1_bambia_26_Dla50 | GAGGAGGGCAGCAAACGGaaTCGCTACGGAGCATTCGCAACGCCA |
| B1_bambia_26_Dla50 | TGGCGCTGTGCCTGGAGACGCTTGTtaGAAGAGTCTTCCTTTACG |
| B1_bambia_26_Dla50 | GAGGAGGGCAGCAAACGGaaCCGCGATGGGAACCGCTATCACCGC |
| B1_bambia_26_Dla50 | TAATCAGCAGAACCAGGATAAGCCCtaGAAGAGTCTTCCTTTACG |
| B1_bambia_26_Dla50 | GAGGAGGGCAGCAAACGGaaTAACTCTTGCACCCTTGTGATCAGG |
| B1_bambia_26_Dla50 | CCGGAACCACACCTCTTTAGCAGACtaGAAGAGTCTTCCTTTACG |
| B1_bambia_26_Dla50 | GAGGAGGGCAGCAAACGGaaGAGTCCCCTCTGGGGTGTGTGAGGT |
| B1_bambia_26_Dla50 | TGATTGGAGCTGTGGTATCGGTCTGtaGAAGAGTCTTCCTTTACG |
| B1_bambia_26_Dla50 | GAGGAGGGCAGCAAACGGaaCATGGCAGCACTCCACAGGAGAGGA |
| B1_bambia_26_Dla50 | GCAAACCCCTGTAGTTACACATATCtaGAAGAGTCTTCCTTTACG |
| B1_bambia_26_Dla50 | GAGGAGGGCAGCAAACGGaaAGAGCACACGTCTGCAGAGTTTAAA |
| B1_bambia_26_Dla50 | TCCACTTGAAATGTCCACATTTTTAtaGAAGAGTCTTCCTTTACG |
| B1_bambia_26_Dla50 | GAGGAGGGCAGCAAACGGaaGAGTTTGTGTTAAGAGGGTCCAGGA |
| B1_bambia_26_Dla50 | GAATCCACGCAGCCGTGTGTTAAAGtaGAAGAGTCTTCCTTTACG |
| B1_bambia_26_Dla50 | GAGGAGGGCAGCAAACGGaaTACACATGTATCCGGTGGCAACGCA |
| B1_bambia_26_Dla50 | TAGTAAAGCAAGCGTTGAGCTCTGAtaGAAGAGTCTTCCTTTACG |
| B1_bambia_26_Dla50 | GAGGAGGGCAGCAAACGGaaTCCTTTCGTGAGAAGAACAGCCATC |
| B1_bambia_26_Dla50 | CGGTGCGTCACAGTAGCACCTGATCtaGAAGAGTCTTCCTTTACG |
| B1_bambia_26_Dla50 | GAGGAGGGCAGCAAACGGaaACCAGGCGATCCATTTACGGCTGCT |
| B1_bambia_26_Dla50 | CAAAGTTCCAGCTGAAACCACAGAGtaGAAGAGTCTTCCTTTACG |
| B1_bambia_26_Dla50 | GAGGAGGGCAGCAAACGGaaCCTCCGAATGGTTTGAGAGGACTCG |
| B1_bambia_26_Dla50 | CCTTGTCTTGATACCCCGTGGCAAGtaGAAGAGTCTTCCTTTACG |
| B1_bambia_26_Dla50 | GAGGAGGGCAGCAAACGGaaCCCAGGGATCTGTGATCTACACTGT |
| B1_bambia_26_Dla50 | CATATGTCCTACATTTATTACTGCAtaGAAGAGTCTTCCTTTACG |

| Sequence name | Sequence |
|---|---|
| B2_gata2a_42_Dla100 | CCTCGTAAATCCTCATCAaaTGTGGTTCGGCCCAGGCGGGAGAGC |
| B2_gata2a_42_Dla100 | CTGTGGAGGGGATGCTGGGTTGTTCaaATCATCCAGTAAACCGCC |
| B2_gata2a_42_Dla100 | CCTCGTAAATCCTCATCAaaGGTGAATGGGGGTCGGCGTGGGCAG |

| | |
|---|---|
| B2_gata2a_42_Dla100 | CGGAATGGTGCGGGTGGCTGAATGTaaATCATCCAGTAAACCGCC |
| B2_gata2a_42_Dla100 | CCTCGTAAATCCTCATCAaaGTGGCCCATGTGTGGCATGTGGCTT |
| B2_gata2a_42_Dla100 | GTGTCCGGAATGGCTGAACGGTGGCaaATCATCCAGTAAACCGCC |
| B2_gata2a_42_Dla100 | CCTCGTAAATCCTCATCAaaGTCTTGTCCTGCATGCACTTGGAGA |
| B2_gata2a_42_Dla100 | AGGGCGCTGGCGCTGCCAAACGGAGaaATCATCCAGTAAACCGCC |
| B2_gata2a_42_Dla100 | CCTCGTAAATCCTCATCAaaTCCTCTTGGACTTGCTGGACATCTT |
| B2_gata2a_42_Dla100 | CCTCGAAACCCTCACCAGATCGTTTaaATCATCCAGTAAACCGCC |
| B2_gata2a_42_Dla100 | CCTCGTAAATCCTCATCAaaCATGGTCAGTGGCCTGTTGACGTTG |
| B2_gata2a_42_Dla100 | GTTGCGTGTTTGGATGCCCTCCTTCaaATCATCCAGTAAACCGCC |
| B2_gata2a_42_Dla100 | CCTCGTAAATCCTCATCAaaCACACCGGGTCACCGTTCCCGTTGC |
| B2_gata2a_42_Dla100 | AGTTTGTAGTAGAGCCCGCAGGCGTaaATCATCCAGTAAACCGCC |
| B2_gata2a_42_Dla100 | CCTCGTAAATCCTCATCAaaAGTTCGCGCAGCAGGTGCCAGCTCG |
| B2_gata2a_42_Dla100 | GCCACAGGGTGGTCGTGGTTGTCTGaaATCATCCAGTAAACCGCC |
| B2_gata2a_42_Dla100 | CCTCGTAAATCCTCATCAaaGATAAGGGGTCTGTTCTGGCCGTTC |
| B2_gata2a_42_Dla100 | CGCCGCAGACAGTCTGCGCTTTGGCaaATCATCCAGTAAACCGCC |
| B2_gata2a_42_Dla100 | CCTCGTAAATCCTCATCAaaAGGTAGTGGCCCGTTCCATCCCGCC |
| B2_gata2a_42_Dla100 | TTGTGGTACAGGCCGCACGCGTTGCaaATCATCCAGTAAACCGCC |
| B2_gata2a_42_Dla100 | CCTCGTAAATCCTCATCAaaTCACACATTCACGCCCCTCTGAGCA |
| B2_gata2a_42_Dla100 | ACAGCGGGGTGGAGGTGGCTCCACAaaATCATCCAGTAAACCGCC |
| B2_gata2a_42_Dla100 | CCTCGTAAATCCTCATCAaaGAAGCTGGAAGCGGATCCGCTGAGG |
| B2_gata2a_42_Dla100 | TCGCGTCTTGCTTTTACATTTAGGTaaATCATCCAGTAAACCGCC |
| B2_gata2a_42_Dla100 | CCTCGTAAATCCTCATCAaaTAGTCGTGTGGCGCGGGCAGGGAGT |
| B2_gata2a_42_Dla100 | GTACCGGGATGAAACAGGCCGCCGCaaATCATCCAGTAAACCGCC |
| B2_gata2a_42_Dla100 | CCTCGTAAATCCTCATCAaaTGGAGGGTGTCTGTGCACTCATAGC |
| B2_gata2a_42_Dla100 | TTGGGTAGGTCGGGATGGGATGATGaaATCATCCAGTAAACCGCC |
| B2_gata2a_42_Dla100 | CCTCGTAAATCCTCATCAaaCATTTTCATGCCATCTGCGATGGAC |
| B2_gata2a_42_Dla100 | GCTTCCCCGAAGAGGACTACATCCCaaATCATCCAGTAAACCGCC |
| B2_gata2a_42_Dla100 | CCTCGTAAATCCTCATCAaaATTCTTGCAGTGGTGGATGTCGGAG |
| B2_gata2a_42_Dla100 | TGGTACTTGATGGACTCCTTTTCGTaaATCATCCAGTAAACCGCC |
| B2_gata2a_42_Dla100 | CCTCGTAAATCCTCATCAaaTGGAGTGGGTTGCGGATGTGAGGGA |
| B2_gata2a_42_Dla100 | GGGGGAAATTATAAAGGGGATGCGGaaATCATCCAGTAAACCGCC |
| B2_gata2a_42_Dla100 | CCTCGTAAATCCTCATCAaaGCACGGATAAGCGGCGCTGGCGGGG |
| B2_gata2a_42_Dla100 | AACTGGAGCCGTGCTTGAACTGCTGaaATCATCCAGTAAACCGCC |
| B2_gata2a_42_Dla100 | CCTCGTAAATCCTCATCAaaGCCCAGGCGTTGTGGTGGGCGGCGG |
| B2_gata2a_42_Dla100 | AGGCCGGGCTTGCTGAAGTGGCTGAaaATCATCCAGTAAACCGCC |
| B2_gata2a_42_Dla100 | CCTCGTAAATCCTCATCAaaGAATACCGGGACTGTGTATGAGGTG |
| B2_gata2a_42_Dla100 | GCGCTGCCTTCCCGCTGTCCAGCCAaaATCATCCAGTAAACCGCC |

107

| **Sequence name** | **Sequence** |
|---|---|
| B3_szl_23_Dla100 | GTCCCTGCCTCTATATCTttATCTTTACAAATTACATCTGAATAG |
| B3_szl_23_Dla100 | ACATACAATACACTTATACATCTGAttCCACTCAACTTTAACCCG |
| B3_szl_23_Dla100 | GTCCCTGCCTCTATATCTttCAACAGTTGGAGTGCATCTCAAGTC |
| B3_szl_23_Dla100 | ATAATAATAATATTTCATCTCTATGttCCACTCAACTTTAACCCG |
| B3_szl_23_Dla100 | GTCCCTGCCTCTATATCTttTGTGTGTGTGTTAAATCATATCTAG |
| B3_szl_23_Dla100 | GATCAAAGAGCTGTGTGTGTGTGTGttCCACTCAACTTTAACCCG |
| B3_szl_23_Dla100 | GTCCCTGCCTCTATATCTttAATGTGAGGTACAATACCATTTGCG |
| B3_szl_23_Dla100 | ACAATATTGTAAGCAAAAATGAATAttCCACTCAACTTTAACCCG |
| B3_szl_23_Dla100 | GTCCCTGCCTCTATATCTttTCTGCATCTGCAATTCGAACAAAGT |
| B3_szl_23_Dla100 | AAAAACTTGAAATGTTTGTTTTGGTttCCACTCAACTTTAACCCG |

| | |
|---|---|
| B3_szl_23_Dla100 | GTCCCTGCCTCTATATCTttCATAACCACACCCCTCTTATTTGTA |
| B3_szl_23_Dla100 | TATATTCTTAGCCACACCCCTTTTAttCCACTCAACTTTAACCCG |
| B3_szl_23_Dla100 | GTCCCTGCCTCTATATCTttCTCTTATTTGTAGCCACGCCCCTCT |
| B3_szl_23_Dla100 | ACACCTGTCTTATCCATAACCACACttCCACTCAACTTTAACCCG |
| B3_szl_23_Dla100 | GTCCCTGCCTCTATATCTttGGAAGTACGTCAACATACTGAAATA |
| B3_szl_23_Dla100 | CCCCCTGCTTAATATTCAGTTTCAGttCCACTCAACTTTAACCCG |
| B3_szl_23_Dla100 | GTCCCTGCCTCTATATCTttATAAGTTGCGCTTTCTCTCAGCACT |
| B3_szl_23_Dla100 | TTTTTCTATATTTTTATAAATATATttCCACTCAACTTTAACCCG |
| B3_szl_23_Dla100 | GTCCCTGCCTCTATATCTttCGATGTGCAAGTCTTTCTTCAGCCA |
| B3_szl_23_Dla100 | AGTGTTTCCACTTGCGTGTTGCCGTttCCACTCAACTTTAACCCG |
| B3_szl_23_Dla100 | GTCCCTGCCTCTATATCTttACCAGCACGCATTTTCCCGGTGATG |
| B3_szl_23_Dla100 | GAACAGATTAGCGATCTCGATGGATttCCACTCAACTTTAACCCG |
| B3_szl_23_Dla100 | GTCCCTGCCTCTATATCTttAGCGCATGCGCACAGCGGAGGTTGA |
| B3_szl_23_Dla100 | TATAACTGCGCACGGCCGGGTCTGAttCCACTCAACTTTAACCCG |
| B3_szl_23_Dla100 | GTCCCTGCCTCTATATCTttTGTCATAGGGCATTAGCGGCCCGCG |
| B3_szl_23_Dla100 | GCATCCAGCGCTGCAGCAGACTAGCttCCACTCAACTTTAACCCG |
| B3_szl_23_Dla100 | GTCCCTGCCTCTATATCTttAAACTCCGGCTCGGAAGACGGCACA |
| B3_szl_23_Dla100 | GATCAGCTCCACAGCGCCCTCTACTttCCACTCAACTTTAACCCG |
| B3_szl_23_Dla100 | GTCCCTGCCTCTATATCTttAAGTCGTTGAGGCAGAGCGAGTCCA |
| B3_szl_23_Dla100 | CGACGCATGATCTTCACCTTCACCGttCCACTCAACTTTAACCCG |
| B3_szl_23_Dla100 | GTCCCTGCCTCTATATCTttCTGATGGACAGCTCTGACATGCTGG |
| B3_szl_23_Dla100 | GGGTTTTGAGTGACGGAGACTCCTGttCCACTCAACTTTAACCCG |
| B3_szl_23_Dla100 | GTCCCTGCCTCTATATCTttGTGTTTGGGCAGCGGCGTCAGACAG |
| B3_szl_23_Dla100 | AGGGAACTCTTTGGAGAAAGCACTGttCCACTCAACTTTAACCCG |
| B3_szl_23_Dla100 | GTCCCTGCCTCTATATCTttAGTGATTCAGGCCAGGAGTGTCCGT |
| B3_szl_23_Dla100 | TCCTGCTCCGGGAATCGCTCACAATttCCACTCAACTTTAACCCG |
| B3_szl_23_Dla100 | GTCCCTGCCTCTATATCTttTTACTGCCAAACACACACTCCGGCA |
| B3_szl_23_Dla100 | AGGCTAGAACGGGACTGCAGCTCTCttCCACTCAACTTTAACCCG |
| B3_szl_23_Dla100 | GTCCCTGCCTCTATATCTttGGCGATGAGCGAGCAGACGAAGGCC |
| B3_szl_23_Dla100 | CTGGATGAACCTGTCGAGGCATACAttCCACTCAACTTTAACCCG |

| **Sequence name** | **Sequence** |
|---|---|
| B1_spry4_55_Dla200 | GAGGAGGGCAGCAAACGGaaGCAATGGGGACTCGGAATCCTTCAG |
| B1_spry4_55_Dla200 | CTGCCACGAGGATACCTGGCGTCTTtaGAAGAGTCTTCCTTTACG |
| B1_spry4_55_Dla200 | GAGGAGGGCAGCAAACGGaaTCAAACAACACAAGAAATAAAAGCT |
| B1_spry4_55_Dla200 | CACGCAATCACCTCCCATGTTTGCAtaGAAGAGTCTTCCTTTACG |
| B1_spry4_55_Dla200 | GAGGAGGGCAGCAAACGGaaAGAGATGAAGTCTGGTTATTTTGAT |
| B1_spry4_55_Dla200 | GAACAAGCTCGTAAATGCACTGTGCtaGAAGAGTCTTCCTTTACG |
| B1_spry4_55_Dla200 | GAGGAGGGCAGCAAACGGaaTGCACTTTCTGTCTTCCGGATCCGG |
| B1_spry4_55_Dla200 | GACATGTGTTATCAGCTTGGAATTCtaGAAGAGTCTTCCTTTACG |
| B1_spry4_55_Dla200 | GAGGAGGGCAGCAAACGGaaGAATGGGTGGTTTCTCAAAGAGAAA |
| B1_spry4_55_Dla200 | GGGATTGCCAAAGTGGAGAACGCTTtaGAAGAGTCTTCCTTTACG |
| B1_spry4_55_Dla200 | GAGGAGGGCAGCAAACGGaaTCTGTGTTTGCAAGGTCCTTTAGTA |
| B1_spry4_55_Dla200 | AACAGAGATCTTTTGGAAATGCAGCtaGAAGAGTCTTCCTTTACG |
| B1_spry4_55_Dla200 | GAGGAGGGCAGCAAACGGaaAAGAACTTCTTTGCAGTACGTGAAA |
| B1_spry4_55_Dla200 | GCTAATACGTAGAACCATCGTGAGGtaGAAGAGTCTTCCTTTACG |
| B1_spry4_55_Dla200 | GAGGAGGGCAGCAAACGGaaCTACTGATTTCAACACCTTAAGTAC |
| B1_spry4_55_Dla200 | TCATTTGTGCCCTTCAGTCAAACATtaGAAGAGTCTTCCTTTACG |

| | |
|---|---|
| B1_spry4_55_Dla200 | GAGGAGGGCAGCAAACGGaaATCATGAGGCTTGTTTTCTGGCTG |
| B1_spry4_55_Dla200 | GATTTTGGGAGGAAGGTCCTGCAAAtaGAAGAGTCTTCCTTTACG |
| B1_spry4_55_Dla200 | GAGGAGGGCAGCAAACGGaaCTGGGTGCTTTTGCATCGGCATCCT |
| B1_spry4_55_Dla200 | GGCCTTGATTTCTGCCACCTTGCAGtaGAAGAGTCTTCCTTTACG |
| B1_spry4_55_Dla200 | GAGGAGGGCAGCAAACGGaaGAAAGCTTGGCACAGCCAGTGGCAG |
| B1_spry4_55_Dla200 | CGGCTGACGCCGTCGTAGCATTTCTtaGAAGAGTCTTCCTTTACG |
| B1_spry4_55_Dla200 | GAGGAGGGCAGCAAACGGaaATGAGACGGCTGCCATGAAGGACCA |
| B1_spry4_55_Dla200 | AATAGCACACCAGGCAGGGCAAAACtaGAAGAGTCTTCCTTTACG |
| B1_spry4_55_Dla200 | GAGGAGGGCAGCAAACGGaaGCATGGTTTGTCCGCGCAGGAGCCT |
| B1_spry4_55_Dla200 | TGCGCAACAGTTCGAGTGCGAACAGtaGAAGAGTCTTCCTTTACG |
| B1_spry4_55_Dla200 | GAGGAGGGCAGCAAACGGaaACCCCCTTGACTAGACACATGCAAG |
| B1_spry4_55_Dla200 | TCGTCCTCATCAGTGCAGTGGTAGAtaGAAGAGTCTTCCTTTACG |
| B1_spry4_55_Dla200 | GAGGAGGGCAGCAAACGGaaACAGACACTCTTGGTTGCAAACCCA |
| B1_spry4_55_Dla200 | CCGAGTCCACTAGGTTTTGTGCGGAtaGAAGAGTCTTCCTTTACG |
| B1_spry4_55_Dla200 | GAGGAGGGCAGCAAACGGaaTTCGGTGCATCGGCATTTCCCGCAT |
| B1_spry4_55_Dla200 | AGAAGGCAAGGTTCGGGGCAGCGTGtaGAAGAGTCTTCCTTTACG |
| B1_spry4_55_Dla200 | GAGGAGGGCAGCAAACGGaaTCTTCAGGCAAGGCTGCCAGTGTCT |
| B1_spry4_55_Dla200 | TCGCACAGCAGCACATGCTTTTTCTtaGAAGAGTCTTCCTTTACG |
| B1_spry4_55_Dla200 | GAGGAGGGCAGCAAACGGaaGCTCCGCAGCTAGAGTCCTGCCGTG |
| B1_spry4_55_Dla200 | TGTTCTTGGAGCTCAGTATCTTGGGtaGAAGAGTCTTCCTTTACG |
| B1_spry4_55_Dla200 | GAGGAGGGCAGCAAACGGaaAGGAGTAGGAGCTGCGTGATCCAGC |
| B1_spry4_55_Dla200 | ATTCCCGGTCGTATACGGATCCACAtaGAAGAGTCTTCCTTTACG |
| B1_spry4_55_Dla200 | GAGGAGGGCAGCAAACGGaaCTGCTGATGGAGCTAGGCCGACCGC |
| B1_spry4_55_Dla200 | CGCTGATCTGAAGACGTGCTACTGCtaGAAGAGTCTTCCTTTACG |

| **Sequence name** | **Sequence** |
|---|---|
| B2_dusp6_49_Dla100 | CCTCGTAAATCCTCATCAaaGTGAAGTACAGTGGCTGGGTTGGGG |
| B2_dusp6_49_Dla100 | TGAAAAACGTTGTGATTGGTTGGTGaaATCATCCAGTAAACCGCC |
| B2_dusp6_49_Dla100 | CCTCGTAAATCCTCATCAaaGTCCTAACGTGCGCTCAAAGTCCAA |
| B2_dusp6_49_Dla100 | GCACACGGTTATCACACGGACTCTTaaATCATCCAGTAAACCGCC |
| B2_dusp6_49_Dla100 | CCTCGTAAATCCTCATCAaaGATGTTCGACTTCTTCATTTTGACA |
| B2_dusp6_49_Dla100 | TTGACCCATGAAGTTAAAGTTGGGCaaATCATCCAGTAAACCGCC |
| B2_dusp6_49_Dla100 | CCTCGTAAATCCTCATCAaaAGCTTCTGCATGAGGTACGCCACTG |
| B2_dusp6_49_Dla100 | TCATAAGCATCGTTCATGGACAGGTaaATCATCCAGTAAACCGCC |
| B2_dusp6_49_Dla100 | CCTCGTAAATCCTCATCAaaGGCAGTGAACAAGCACGCCACACTT |
| B2_dusp6_49_Dla100 | CAGTGACAGAACGACTGATGCCTGCaaATCATCCAGTAAACCGCC |
| B2_dusp6_49_Dla100 | CCTCGTAAATCCTCATCAaaGGCTTCAGGGAAAAACTGTGAGAGG |
| B2_dusp6_49_Dla100 | TCCACGGGCCTCATCAATAAAGCTGaaATCATCCAGTAAACCGCC |
| B2_dusp6_49_Dla100 | CCTCGTAAATCCTCATCAaaTGCTTGTACTTAAACTCCCCGGCAT |
| B2_dusp6_49_Dla100 | TGGCTCCAGTGATCAGAGATGGGAAaaATCATCCAGTAAACCGCC |
| B2_dusp6_49_Dla100 | CCTCGTAAATCCTCATCAaaCGTTCAAGATGTACTTGATGCCAAA |
| B2_dusp6_49_Dla100 | CGAACATGTTGGGGAGATTAGGGGTaaATCATCCAGTAAACCGCC |
| B2_dusp6_49_Dla100 | CCTCGTAAATCCTCATCAaaCTTAGCACAGCCCAGATACAGATGT |
| B2_dusp6_49_Dla100 | CTCCAGGATATCCAGGTTTGTGGAGaaATCATCCAGTAAACCGCC |
| B2_dusp6_49_Dla100 | CCTCGTAAATCCTCATCAaaGGGTTGGAAAGGGGGCTGCCATCTG |
| B2_dusp6_49_Dla100 | AGGATCTCCACGGGGAATGAGGGCTaaATCATCCAGTAAACCGCC |
| B2_dusp6_49_Dla100 | CCTCGTAAATCCTCATCAaaTATCCGACTCGATGTCCGAGGAGTC |
| B2_dusp6_49_Dla100 | CAGTTGCACTGCTTGGGTCTCGGTCaaATCATCCAGTAAACCGCC |
| B2_dusp6_49_Dla100 | CCTCGTAAATCCTCATCAaaGACCTGGGAGGTTGGGGAACTGCTG |

| | |
|---|---|
| B2_dusp6_49_Dla100 | GCTGATTCTGAGCCCTCCGAGACCCaaATCATCCAGTAAACCGCC |
| B2_dusp6_49_Dla100 | CCTCGTAAATCCTCATCAaaATTGCGGGAAAATCAGTTTGAAATT |
| B2_dusp6_49_Dla100 | GAGGAACCGTCGAGGTTCGTCTCACaaATCATCCAGTAAACCGCC |
| B2_dusp6_49_Dla100 | CCTCGTAAATCCTCATCAaaTGTAGCCCTCGTCCTTCATTCTCCT |
| B2_dusp6_49_Dla100 | TGAAGCCACCCTCGAGATAGAAAGCaaATCATCCAGTAAACCGCC |
| B2_dusp6_49_Dla100 | CCTCGTAAATCCTCATCAaaGATGTTTTCATTCCACTCGCGGCTG |
| B2_dusp6_49_Dla100 | CAGTAAACCCAACACGGAGCCGCCGaaATCATCCAGTAAACCGCC |
| B2_dusp6_49_Dla100 | CCTCGTAAATCCTCATCAaaTTGCATCTCCGCGCGAACCTTTCCC |
| B2_dusp6_49_Dla100 | TCGTCGTACAACACGATCGTGTCCGaaATCATCCAGTAAACCGCC |
| B2_dusp6_49_Dla100 | CCTCGTAAATCCTCATCAaaTGGGCAGGTTGCCTTTCTTGAGTCG |
| B2_dusp6_49_Dla100 | CTTCCCCGTTAGAAAGCAGAGACTTaaATCATCCAGTAAACCGCC |
| B2_dusp6_49_Dla100 | CCTCGTAAATCCTCATCAaaAATGGCCGTTTCGACGTGCGACGAC |
| B2_dusp6_49_Dla100 | GAGCATGAGGCTCGGGATGGCCACGaaATCATCCAGTAAACCGCC |
| B2_dusp6_49_Dla100 | CCTCGTAAATCCTCATCAaaACGAGCAAACAGTCTCTGCGCGTTT |
| B2_dusp6_49_Dla100 | TACAGCTCTTGCGCTCGGCAGTCCAaaATCATCCAGTAAACCGCC |
| B2_dusp6_49_Dla100 | CCTCGTAAATCCTCATCAaaTGCTTATGGCCATGACCGAATCGAT |
| B2_dusp6_49_Dla100 | GCTGCTCCTTCAGCCACTCTACCGTaaATCATCCAGTAAACCGCC |

| **Sequence name** | **Sequence** |
|---|---|
| B3_il17rd_52_Dla100 | GTCCCTGCCTCTATATCTttCTGAATCTACTTTTTCAGGAATGGT |
| B3_il17rd_52_Dla100 | TGAGTTTGACTTCATTTAGCACCAAttCCACTCAACTTTAACCCG |
| B3_il17rd_52_Dla100 | GTCCCTGCCTCTATATCTttTTCCTTCTCCAGCCAATCAGGCTCC |
| B3_il17rd_52_Dla100 | TTTATTAGGCAGCGGCGGAGGCATGttCCACTCAACTTTAACCCG |
| B3_il17rd_52_Dla100 | GTCCCTGCCTCTATATCTttGCAACATATAACGAGCGCCCCGATT |
| B3_il17rd_52_Dla100 | GTGACGTACTGATGCATGTTATAAAttCCACTCAACTTTAACCCG |
| B3_il17rd_52_Dla100 | GTCCCTGCCTCTATATCTttTCGGCGGCTGTGGTTCACGATCGGT |
| B3_il17rd_52_Dla100 | TGCAGAAGTAGTTGCGTTTGCTAACttCCACTCAACTTTAACCCG |
| B3_il17rd_52_Dla100 | GTCCCTGCCTCTATATCTttGAAGAGTTGCGGCAGCTGATCCATC |
| B3_il17rd_52_Dla100 | ACTCAGTTGGCGCGAATGCAATCGGttCCACTCAACTTTAACCCG |
| B3_il17rd_52_Dla100 | GTCCCTGCCTCTATATCTttGTCGGGACATCGGTTTCATGGGAAT |
| B3_il17rd_52_Dla100 | TTAAACTTCGGCGCCAGGCTGAGTGttCCACTCAACTTTAACCCG |
| B3_il17rd_52_Dla100 | GTCCCTGCCTCTATATCTttGGTCGGAGGATTTCTGGTGCACTTC |
| B3_il17rd_52_Dla100 | CGAAATACACAGACATGAAGCGCGAttCCACTCAACTTTAACCCG |
| B3_il17rd_52_Dla100 | GTCCCTGCCTCTATATCTttCACGATGAACAGATCTCTGCTGCTG |
| B3_il17rd_52_Dla100 | GAGTTTCTCTGAGATGATGGCGGAGttCCACTCAACTTTAACCCG |
| B3_il17rd_52_Dla100 | GTCCCTGCCTCTATATCTttTTTTTCTCTTTGGATGTGGCTTTGC |
| B3_il17rd_52_Dla100 | CTGGAGTCGCTAGCGCTCGGCTCTCttCCACTCAACTTTAACCCG |
| B3_il17rd_52_Dla100 | GTCCCTGCCTCTATATCTttTCAGGCCTTTGGAGCAGACGGTGAT |
| B3_il17rd_52_Dla100 | TGCGGTGGCGTTTCTCCACGAAGTGttCCACTCAACTTTAACCCG |
| B3_il17rd_52_Dla100 | GTCCCTGCCTCTATATCTttCAACCAGGACATCTGACCCTCCTTA |
| B3_il17rd_52_Dla100 | GAAGTGGGCTTCGTCGATGCGCCGGttCCACTCAACTTTAACCCG |
| B3_il17rd_52_Dla100 | GTCCCTGCCTCTATATCTttGAAACCTCGCAGCCGCAGAAGTCCT |
| B3_il17rd_52_Dla100 | ATCTCCAGGTGCTCCCAGAGGTCCAttCCACTCAACTTTAACCCG |
| B3_il17rd_52_Dla100 | GTCCCTGCCTCTATATCTttCGAGGTGTTTGGCTCCGTCTCTGCT |
| B3_il17rd_52_Dla100 | GGAAGAAGGCGAAGCTCTGGATGACttCCACTCAACTTTAACCCG |
| B3_il17rd_52_Dla100 | GTCCCTGCCTCTATATCTttCCAGGGTCTGTCTGCGCTCAGAGCC |
| B3_il17rd_52_Dla100 | GTAACAGATGAAGATTTTGGGTCTGttCCACTCAACTTTAACCCG |
| B3_il17rd_52_Dla100 | GTCCCTGCCTCTATATCTttTCGTCCAGGTGAGAGTAGATGTTTT |
| B3_il17rd_52_Dla100 | GTCTGCGATGAAGACTCCGAGCTCTttCCACTCAACTTTAACCCG |

| | |
|---|---|
| B3_il17rd_52_Dla100 | GTCCCTGCCTCTATATCTttAGAGTGTGGCGAAGGCTGACATGAT |
| B3_il17rd_52_Dla100 | GCTGTTTCTTGCGGCACATGACGGTttCCACTCAACTTTAACCCG |
| B3_il17rd_52_Dla100 | GTCCCTGCCTCTATATCTttGATCGGCCCGGCCCACGGGGAGTGA |
| B3_il17rd_52_Dla100 | CAGCGGGACGGTGATGGCCATGGCAttCCACTCAACTTTAACCCG |
| B3_il17rd_52_Dla100 | GTCCCTGCCTCTATATCTttCTGGTGTTGTTGCTGTCATCACGAA |
| B3_il17rd_52_Dla100 | TGGCTGACGTGATACTGTGTCTGTCttCCACTCAACTTTAACCCG |
| B3_il17rd_52_Dla100 | GTCCCTGCCTCTATATCTttCTTGTAAAACACAAGTAGTTTGGG |
| B3_il17rd_52_Dla100 | CGATTGCATACGTTCCTGGAGTGACttCCACTCAACTTTAACCCG |
| B3_il17rd_52_Dla100 | GTCCCTGCCTCTATATCTttTCTGAACGGTCCTTCCTGTCTGAGT |
| B3_il17rd_52_Dla100 | GTTTTGCTCAGGTTTGCAGCGCTTGttCCACTCAACTTTAACCCG |

112

| **Sequence name** | **Sequence** |
|---|---|
| B1_noto_16_Dla100 | GAGGAGGGCAGCAAACGGaaTTCATCTATATTCAAACATTTTGAC |
| B1_noto_16_Dla100 | TTGATTTTAAAAATATATGTCCTTTtaGAAGAGTCTTCCTTTACG |
| B1_noto_16_Dla100 | GAGGAGGGCAGCAAACGGaaGGCGTGTGTAAGTGACGCCTTAAAT |
| B1_noto_16_Dla100 | AAACAAAGATACCCTTATAAGAGCAtaGAAGAGTCTTCCTTTACG |
| B1_noto_16_Dla100 | GAGGAGGGCAGCAAACGGaaTAGAGAGGAGTAACAATCTGGGGAA |
| B1_noto_16_Dla100 | TTTCTATTAATTAAATATTTGTACAtaGAAGAGTCTTCCTTTACG |
| B1_noto_16_Dla100 | GAGGAGGGCAGCAAACGGaaTGTGAGAAAGTCTCAGTCTTGCAGT |
| B1_noto_16_Dla100 | CATGGAAAAGTATGTACAAAAACACtaGAAGAGTCTTCCTTTACG |
| B1_noto_16_Dla100 | GAGGAGGGCAGCAAACGGaaTCTTCTGTGAAATCCCTCTCCTCAT |
| B1_noto_16_Dla100 | TCGTCAATGTCAATGTCTACATCAGtaGAAGAGTCTTCCTTTACG |
| B1_noto_16_Dla100 | GAGGAGGGCAGCAAACGGaaGTGGAACGGTCAGTCCCAGTTTGGC |
| B1_noto_16_Dla100 | CTCTGCCCTGGGATCCAGGGCTTTTtaGAAGAGTCTTCCTTTACG |
| B1_noto_16_Dla100 | GAGGAGGGCAGCAAACGGaaCTTTCTCCATTTGATGCGCCTGTTC |
| B1_noto_16_Dla100 | TTTGGCTTGTTGTTGCTCAAGACTCtaGAAGAGTCTTCCTTTACG |
| B1_noto_16_Dla100 | GAGGAGGGCAGCAAACGGaaAGTTGGAGAGCAGATGCCAACAGAA |
| B1_noto_16_Dla100 | AACCAGACTTTAACCTGAGCTTCAGtaGAAGAGTCTTCCTTTACG |
| B1_noto_16_Dla100 | GAGGAGGGCAGCAAACGGaaGCGCGAACTCTTTCTCCAGTCTGGA |
| B1_noto_16_Dla100 | GTTCAGATCCCACCATGTATTGCTGtaGAAGAGTCTTCCTTTACG |
| B1_noto_16_Dla100 | GAGGAGGGCAGCAAACGGaaTCGCTTTGATTTCCCAGATTTGTGT |
| B1_noto_16_Dla100 | CTGATCGTTGGTAAAACTTGTACGCtaGAAGAGTCTTCCTTTACG |
| B1_noto_16_Dla100 | GAGGAGGGCAGCAAACGGaaAGTACTGCATCTTGTGCGTACACAG |
| B1_noto_16_Dla100 | TAATGAGAATGAACTGCTGCTTTCGtaGAAGAGTCTTCCTTTACG |
| B1_noto_16_Dla100 | GAGGAGGGCAGCAAACGGaaGCGGATAGCAGAACACGGGATAGCC |
| B1_noto_16_Dla100 | CACGACATGTTGTTTGGAAGTTGTAtaGAAGAGTCTTCCTTTACG |
| B1_noto_16_Dla100 | GAGGAGGGCAGCAAACGGaaCGCGAAGTGTGGCATCTGCGAGTAA |
| B1_noto_16_Dla100 | CTGAGTTTGCATGATGCTTTGGCTGtaGAAGAGTCTTCCTTTACG |
| B1_noto_16_Dla100 | GAGGAGGGCAGCAAACGGaaGTTTGGTTGGTTATGCTCCGGTACG |
| B1_noto_16_Dla100 | AGCGGGAGCGCAGAGCTGGAGACAGtaGAAGAGTCTTCCTTTACG |
| B1_noto_16_Dla100 | GAGGAGGGCAGCAAACGGaaCAGGCCTCGCGAGCAGAGCGTCTAT |
| B1_noto_16_Dla100 | TTGTTCGTTCTCTCATCTCCGCCTGtaGAAGAGTCTTCCTTTACG |
| B1_noto_16_Dla100 | GAGGAGGGCAGCAAACGGaaAGAAGAGGTGGCATAGTCCTGATGC |
| B1_noto_16_Dla100 | GAATGATTTCCCGGTGCTCGGTTTTtaGAAGAGTCTTCCTTTACG |

113

| **Sequence name** | **Sequence** |
|---|---|
| B2_gsc_22_Dla20 | CCTCGTAAATCCTCATCAaaCCTGTTTTCAGGCGACATTAAACTT |
| B2_gsc_22_Dla20 | TTTTTAGATATTACTTTAATATTTGaaATCATCCAGTAAACCGCC |
| B2_gsc_22_Dla20 | CCTCGTAAATCCTCATCAaaTGCCATCGTACATGTCTTCAGCTAC |

| | |
|---|---|
| B2_gsc_22_Dla20 | ATTAATTAATGTCCGAATGTATCGTaaATCATCCAGTAAACCGCC |
| B2_gsc_22_Dla20 | CCTCGTAAATCCTCATCAaaATCTTATCACGACAAGACTTTAAAA |
| B2_gsc_22_Dla20 | CCATTCCAGAACATCAGATTTAGGTaaATCATCCAGTAAACCGCC |
| B2_gsc_22_Dla20 | CCTCGTAAATCCTCATCAaaTATGGCTAGAATCCACGTTATTTG |
| B2_gsc_22_Dla20 | TTTGCAGTATACACCAGGTAATATTaaATCATCCAGTAAACCGCC |
| B2_gsc_22_Dla20 | CCTCGTAAATCCTCATCAaaTATCATTGTTAAAGGTAACATTTAC |
| B2_gsc_22_Dla20 | GTAAATTTAAATTAATATAAATAACaaATCATCCAGTAAACCGCC |
| B2_gsc_22_Dla20 | CCTCGTAAATCCTCATCAaaATTTACTCCAACTCACATACTTTAC |
| B2_gsc_22_Dla20 | ACATCTGTGCAACAGCAAGACAACAaaATCATCCAGTAAACCGCC |
| B2_gsc_22_Dla20 | CCTCGTAAATCCTCATCAaaGTGCAAGATTTCCCGTTCTCGTGTT |
| B2_gsc_22_Dla20 | ATGTACAATAAAGTCCGAATTATATaaATCATCCAGTAAACCGCC |
| B2_gsc_22_Dla20 | CCTCGTAAATCCTCATCAaaTTTTGCCCTCCTCAATTTTCTCTGA |
| B2_gsc_22_Dla20 | TATATCAGCTGTCAGAATCCACGTCaaATCATCCAGTAAACCGCC |
| B2_gsc_22_Dla20 | CCTCGTAAATCCTCATCAaaTGAGTTTTCTGATTCCTCTGACGAC |
| B2_gsc_22_Dla20 | TGTTTTCGTGGATTTGTTCCATTTCaaATCATCCAGTAAACCGCC |
| B2_gsc_22_Dla20 | CCTCGTAAATCCTCATCAaaCTGTTTTTGAACCAAACCTCTACCT |
| B2_gsc_22_Dla20 | CTTTTCTGTCTTCTCCATTTTGCTCaaATCATCCAGTAAACCGCC |
| B2_gsc_22_Dla20 | CCTCGTAAATCCTCATCAaaTGTAGCTGGTTGAGCAGCTGTAGCT |
| B2_gsc_22_Dla20 | GTTCGGTGTCGTCTCTTGCGCCGACaaATCATCCAGTAAACCGCC |
| B2_gsc_22_Dla20 | CCTCGTAAATCCTCATCAaaACGACATCATTTGATGTGGGACTGG |
| B2_gsc_22_Dla20 | TTCTGGACAAGGTGCCCACGTTCATaaATCATCCAGTAAACCGCC |
| B2_gsc_22_Dla20 | CCTCGTAAATCCTCATCAaaACCTGTATGAATACACGGACACTGT |
| B2_gsc_22_Dla20 | AATAAGCACAGAGCCGGCGCTGTCAaaATCATCCAGTAAACCGCC |
| B2_gsc_22_Dla20 | CCTCGTAAATCCTCATCAaaCAAGCTGGTCCAGTCGGCCCCTGGA |
| B2_gsc_22_Dla20 | GAACCAAGGGTTGGTATTGCGCCACaaATCATCCAGTAAACCGCC |
| B2_gsc_22_Dla20 | CCTCGTAAATCCTCATCAaaAGCCTATCCTTCCATTCACCGATTG |
| B2_gsc_22_Dla20 | GAAGTTGTCCATAGTAGTAGTTGTTaaATCATCCAGTAAACCGCC |
| B2_gsc_22_Dla20 | CCTCGTAAATCCTCATCAaaATACAGTCCATTAAAATCGCCAGCT |
| B2_gsc_22_Dla20 | GTTCGGCGCTGGAGGTCCTGTGTGTaaATCATCCAGTAAACCGCC |
| B2_gsc_22_Dla20 | CCTCGTAAATCCTCATCAaaAACACAACCGGGGCATTCCGGTGGA |
| B2_gsc_22_Dla20 | GTGTACAAGGATTCCGTCAAGTTGGaaATCATCCAGTAAACCGCC |
| B2_gsc_22_Dla20 | CCTCGTAAATCCTCATCAaaCGGCCAAGATGCTGTCGATACTAAA |
| B2_gsc_22_Dla20 | GAACCGAGTCTTTGCAGCTGGGTCTaaATCATCCAGTAAACCGCC |
| B2_gsc_22_Dla20 | CCTCGTAAATCCTCATCAaaGTGTGAGATTTGTTGCCAACGGTAA |
| B2_gsc_22_Dla20 | CCCAGCGGGCATCACAAGCGAAAAGaaATCATCCAGTAAACCGCC |
| B2_gsc_22_Dla20 | CCTCGTAAATCCTCATCAaaACGAGTGTCCTTCATAGTGACAAAA |
| B2_gsc_22_Dla20 | CGTGTTATTTTAGTCCTTTTAAAAAaaATCATCCAGTAAACCGCC |

114

| **Sequence name** | **Sequence** |
|---|---|
| B3_lft1_22_Dla100 | GTCCCTGCCTCTATATCTttCTACAAATCAATGGCATCATATACA |
| B3_lft1_22_Dla100 | CCGTGCTATATGCTCAAAATAAAACttCCACTCAACTTTAACCCG |
| B3_lft1_22_Dla100 | GTCCCTGCCTCTATATCTttTCTATTTACAAGTCTATACAAAGTG |
| B3_lft1_22_Dla100 | GCAAAAATAAACGCATATCAGATTAttCCACTCAACTTTAACCCG |
| B3_lft1_22_Dla100 | GTCCCTGCCTCTATATCTttAACACCCATTCTAAATCTTATAAGT |
| B3_lft1_22_Dla100 | GAGGTATCTAGTAAGGGTTAAATTGttCCACTCAACTTTAACCCG |
| B3_lft1_22_Dla100 | GTCCCTGCCTCTATATCTttTGGTCGATACAAACAGGCTATTTAT |
| B3_lft1_22_Dla100 | GGTAGTATAGTGCGTCATGAAGATAttCCACTCAACTTTAACCCG |
| B3_lft1_22_Dla100 | GTCCCTGCCTCTATATCTttCATTTTTCCACAATCATGTTTGGGA |
| B3_lft1_22_Dla100 | ACTGAAATATTGTCCATTGCGCATCttCCACTCAACTTTAACCCG |

| | |
|---|---|
| B3_lft1_22_Dla100 | GTCCCTGCCTCTATATCTttGCGCGCTCTCGACGACCGCGCATTT |
| B3_lft1_22_Dla100 | TTTTTACTAGATACATCATCGGTAGttCCACTCAACTTTAACCCG |
| B3_lft1_22_Dla100 | GTCCCTGCCTCTATATCTttCTGCCGGCAGCCGCCTTTACACCTG |
| B3_lft1_22_Dla100 | CTCTCCGTAGCCGTAGTTGCGCTTTttCCACTCAACTTTAACCCG |
| B3_lft1_22_Dla100 | GTCCCTGCCTCTATATCTttCAGTACTGTGTCCAAGTCAGAGCTC |
| B3_lft1_22_Dla100 | GCCTGGTAACCGGACGGCTCGATGAttCCACTCAACTTTAACCCG |
| B3_lft1_22_Dla100 | GTCCCTGCCTCTATATCTttACATTTCACGGTCTTTGTTGTTTTC |
| B3_lft1_22_Dla100 | AATTGATGAAGTACTGTTCCCTGCAttCCACTCAACTTTAACCCG |
| B3_lft1_22_Dla100 | GTCCCTGCCTCTATATCTttGTTGAGTGTGTAAAGCACCAGCTCT |
| B3_lft1_22_Dla100 | GTCTCCACTAGACCCAAACTCCTCTttCCACTCAACTTTAACCCG |
| B3_lft1_22_Dla100 | GTCCCTGCCTCTATATCTttTGAGTTGTGAAGTGGACACACTTGG |
| B3_lft1_22_Dla100 | TTTCCCAGAGTGTTGTCGTCTGGGTttCCACTCAACTTTAACCCG |
| B3_lft1_22_Dla100 | GTCCCTGCCTCTATATCTttCGCCCTCGATCCACACCTCAAGGTG |
| B3_lft1_22_Dla100 | TCTCTGCCGCGTAACTGCCGGGTCTttCCACTCAACTTTAACCCG |
| B3_lft1_22_Dla100 | GTCCCTGCCTCTATATCTttATATTGCACCGCCTGGGTGACATCA |
| B3_lft1_22_Dla100 | GGGCATCTCCATTCGGCTCCTAGACttCCACTCAACTTTAACCCG |
| B3_lft1_22_Dla100 | GTCCCTGCCTCTATATCTttATCAACCTGGAATCCACTAGTGAAG |
| B3_lft1_22_Dla100 | CTCTTCCAGCCAGTTTCGTGAATGGttCCACTCAACTTTAACCCG |
| B3_lft1_22_Dla100 | GTCCCTGCCTCTATATCTttCCCAGTAGATGCTCACTCGTGCGTT |
| B3_lft1_22_Dla100 | GGTTTGACCCGTCTTTCTGAGGTTCttCCACTCAACTTTAACCCG |
| B3_lft1_22_Dla100 | GTCCCTGCCTCTATATCTttTATGGAGCGCTTGTGTGGGGCCTTC |
| B3_lft1_22_Dla100 | GACCGGTCTGTGGCCCTTTCTCTCCttCCACTCAACTTTAACCCG |
| B3_lft1_22_Dla100 | GTCCCTGCCTCTATATCTttTCACTGTTCTCGGGGATTCTTGATG |
| B3_lft1_22_Dla100 | TAGAGCTTCAGTTCTGCCATGGTCAttCCACTCAACTTTAACCCG |
| B3_lft1_22_Dla100 | GTCCCTGCCTCTATATCTttTGTCAGAATATACGAATTCACCAGA |
| B3_lft1_22_Dla100 | TTTCAAAGACCACACGCTGACGTGTttCCACTCAACTTTAACCCG |
| B3_lft1_22_Dla100 | GTCCCTGCCTCTATATCTttGATGCCGGCCAAACTGGGAAGCGAG |
| B3_lft1_22_Dla100 | GTCTGCATTTCCAGGAATTCCCCTCttCCACTCAACTTTAACCCG |
| B3_lft1_22_Dla100 | GTCCCTGCCTCTATATCTttATGGAGATATATTTGTTCTTTACGT |
| B3_lft1_22_Dla100 | CGTTTTCTTGAGTGGTGAAGTTTCAttCCACTCAACTTTAACCCG |

**Supplementary Information 1: Construct sequences.** All constructs used here are in the ampicillin-resistant pCS2+ vector backbone and can be linearized for mRNA synthesis using NotI.

Key: **Myr**, **linker**, **LOV**, **putative kinase domain**, **HA tag**

*bOpto-FGF* (Addgene #232639)

atggggagtagcaagagcaagcctaaggaccccagccagcgcggtggaggaggttctggaggcggtggaagtggtggcggaggtag
cctggccaaaatgcacagctctgccaagaaaagcgacttcaacagccagctggccgtccacaagctggccaagagcatccccctgcgca
gacaggtaacagtgtctgtggactccagctcatctatgcattcgggtgggatgttggtccgtccatcccgtctgtcctccagtggctccccaat
gctctcaggggtctccgaatacgagcttccccaggacccacgctgggaggtgcaacgagacaggctggttctcgggaaacctcttggcga
aggctgctttggacaggtgatgatggccgaggcgatggggatggataaagaaaaacccaatagaatcaccaaagtggccgtcaagatgct
caaatcggatgccacggagaaagacctgtcagacctgatctctgagatggagatgatgaagattattggcaaacacaagaacatcatcaac
ctgctgggagcctgcacacaagacggtccgctgtacgtcatcgtggagtttgctgctaaagggaacctgcgggagtatctgcgcgtacggc
gtccaccagggatggagtactgttataaccctgaccaggtgccagtggagaacatgtccattaaagacctggtgtcctgcgcatatcaggtg
gcccgaggaatggagtatctcgcatccaagaagtgtattcatcgagacctggctgctcggaatgtgctggtgacggaagataacgttatgaa
gatcgcagactttggcctggccagagacatccatcatattgattactacaagaagaccaccaatggtcgtttgccggtgaaatggatggctcc
cgaagctctgtttgaccgcatatacacccatcaaagtgacgtctggtcttttggggtgctgctgtgggagatcttcactctgggggctctccg
taccccggcgtccccgtcgaggagctctttaagctgctgaaggaaggacaccgcatggaccgaccctccacatgcacacatgagctgtat
atgatgatgagggattgttggcacgccgtcccgtctcagagacccacttttaaacagctggtggaggatctggaccgcacccctctccatgac
gtccaatcaggagtatctggacctgtccgtatctctggaccagttttctccaaacttcccggacactcgcagctccacctgctcctcaggtgaa
gactcagtgttttctcatgacgccggagccgatgagccctgtttgcccaaattcccaccccatcccaaccgaggagtggcctttaaaaagcg
cggtggaggaggttctggaggcggtggaagtggtggcggaggtagccctgactacagtctcgtgaaggctctgcaaatggcacaacaga
attttgtcattacagacgcctccctcccagacaaccctatcgtctacgccagtagagggtttctgacactgacaggctattctctcgaccagatc
ctgggcaggaactgcaggtttctgcaagggccagaaacagacccaagagctgtggataagatcaggaatgccatcaccaaaggcgttgat
accagtgtctgtctgctgaattatagacaggatggcacaaccttctggaatctcttcttcgtggctggactcagagattctaagggcaatattgt
caactacgtcggagtgcagtcaaaggtgagcgaagattatgccaagctgctggtcaacgagcagaacattgagtacaaaggtgtgcgcac
cagtaacatgctgcgcagaaagcccgggtctagttatccgtacgacgtaccagactacgcataa

*bOpto-Bmpr1aa* (Addgene # 207614):

atggggagtagcaagagcaagcctaaggaccccagccagcgcggtggaggaggttctggaggcggtggaagtggtggcggaggtag
ctacaggtataagtggcagacagagaggcagcgctaccacagagacctggagcaagacgaggcctttatcccagcaggagaatccctga
aagacctcatcaaccagtctcagacctcaggcagtggctctggactccctctgctggtgcagcgcactatagcaaagcagatccagacagt
gcgaatgatcggaaaaggacgatatggagaagtgtggcttggtcggtggagaggagagaaggtagcagtgaaggtgttctttacccgaga
ggaggccagctggttcagagagacagagatctatcaaaccgtgctcatgagacatgagaacatactcggctttatcgctgctgatataaatg
gcacaggagcctctacgcagctgtacctgatcacagactaccatgagaatggctctctgtatgactatctgaagttcacgactttggacacac
aggctctactcagactggccttctctgcagcctgtggcctgtgtcacctgcatacggagatctacggcacgcagggaaaaccagcgatcgc
tcacagagacctgaagagcaagaacattctcatcaagaaaaacggcacctgctgcatcgctgacctcggccttgctgtgaaattcaacagtg
acacaaatgaagtggacctcccattaagcacacgtatgggaaccaggcgctacatggctccagaagtgttggacgagactctgaataaga
atcatttccaggcctacatcatggcagacatctacagctatgggctggttatttgggaaatggccagacgctgtgtcactggagggattgttga
ggagtatcagctgccatattatgagatggtgccttcagacccatcttatgaagacatgttggaggttgtttgtgtcaaaggactgcggcccacc
gtatccaacagatggaacagtgatgagtgcttaagggccatgctaaagctgatgtctgaatgctgggcccacaatcctgcatcacgcttaac
catcctacgagtcaagaagactttagccaaaatggtggaatctcaagacattaaaatcggtggaggaggttctggaggcggtggaagtggt
ggcggaggtagccctgactacagtctcgtgaaggctctgcaaatggcacaacagaattttgtcattacagacgcctccctcccagacaacc
ctatcgtctacgccagtagagggtttctgacactgacaggctattctctcgaccagatcctgggcaggaactgcaggtttctgcaagggcca
gaaacagacccaagagctgtggataagatcaggaatgccatcaccaaaggcgttgataccagtgtctgtctgctgaattatagacaggatg
gcacaaccttctggaatctcttcttcgtggctggactcagagattctaagggcaatattgtcaactacgtcggagtgcagtcaaaggtgagcg
aagattatgccaagctgctggtcaacgagcagaacattgagtacaaaggtgtgcgcaccagtaacatgctgcgcagaaagcccgggtcta
gttatccgtacgacgtaccagactacgcataa

*bOpto-Acvr1l* (Addgene # 207615):

atggggagtagcaagagcaagcctaaggaccccagccagcgcggtggaggaggttctggaggcggtggaagtggtggcggaggtagccgtcgactccatcatgggcgtctggagagactgcacgagtttgacactgaacagggggccatcgatgggcttatcgcgtctaatgtcggagacagcacacttgcggatctgatggatcactcctgcacttcaggcagtggttcaggactgcccttcctggttcagagaacggttgcgcggcagatcagcctggtggagtgtgttggtaaaggacggtacggtgaagtgtggagaggtcaatggcaaggagaaaatgtagccgtgaagatcttttcctctagagatgagaagtcatggtttcgagaaacagaaatttacaacactgttctgctacgacatgaaaatatattaggcttcatggcttctgacatgacctcccgaaactctagcactcagctgtggctgatcacacactatcacgagaatggctctctgtatgactacctgcagcgtgtggctgtggagatggcagatggactgcacatggcggcttcgattgccagcgggctggtgcacctgcacacggagatctttggcacggagggcaaaccggccatcgctcacagagacctgaagagcaagaacatccttgtgaagaaagatttgcagtgctgcatcgctgacctgggtctggcagtaacacacacgcagtctgataatcagcttgatgtgggaaataatcctaaagtgggaaccaaacgctacatggcaccggaggttctagatgagaccattcagacggactgttttgacgcctataagagggtggatatctgggcctttgggttggtgctgtgggagatcgcacgcagaaccatcagcaatggaattgtagaggaatacaagccgcctttctatgacctggttcctaatgatcccagctttgacgacatgaggaaagtggtttgtgtggagcagcaaaggccattcattcccaaccgctggttttcagatcctaccctgtctgctctggtgaagctgatgaaagagtgctggtaccagaacccctcggctcgtctcactgccctgcgcatcaaaaagactctggataaaatccacagttcactggagaagggcaaaaccgactgcggtggaggaggttctggaggcggtggaagtggtggcggaggtagccctgactacagtctcgtgaaggctctgcaaatggcacaacagaattttgtcattacagacgcctccctcccagacaaccctatcgtctacgccagtagagggtttctgacactgacaggctattctctcgaccagatcctgggcaggaactgcaggtttctgcaagggccagaaacagacccaagagctgtggataagatcaggaatgccatcaccaaaggcgttgataccagtgtctgtctgctgaattatagacaggatggcacaaccttctggaatctcttcttcgtggctggactcagagattctaagggcaatattgtcaactacgtcggagtgcagtcaaaggtgagcgaagattatgccaagctgctggtcaacgagcagaacattgagtacaaaggtgtgcgcaccagtaacatgctgcgcagaaagcccgggtctagttatccgtacgacgtaccagactacgcataa *bOpto-Bmpr2a* (Addgene # 207616):

atggggagtagcaagagcaagcctaaggaccccagccagcgcggtggaggaggttctggaggcggtggaagtggtggcggaggtagccgcatgctaagaggacgtggtaaacactcactgcatactttaaatatactggagacggcactttctcctccttctttggatctggacaacctcacactgcaggagctgattggccggggccggtacggtacagtgtatcgtgcttcgctggatgaccgatctgtagcagtgaaagttttcatttctgctaaccgccagcagttcactaatgagcgtatgatttatcgcctcctcctggatcacgagaacatagcgcgcttcctagagagcgaggagcgcgtcggcacagaaggtcggacagagtttctccttctgctggagttttatcctcatggctctctgtgcacgtatctgagcggccggactgtggactggttgagctgctgccgtttggctctgtctgtgaccagagggttggcgtacctacatacagagatacagcgaggggatgtgtataaaccggctgtctcccaccgggatctgaacagcaggaatgtgctggttaagacggacggctcgtgtgtgatcagtgactttggactgtccatgattctgataggaaagaggccgcctggtcatggagaagaggacaacagtgccatcagtgaggtaggtacagtgcggtacatggctccagaagtgctggaaggggctgtgaatctgagggattgtgagtctgcgctgaaacaagtggatgtttatgcactgggtctggtttactgggagaccttcatgcgctgtgcagatcttttcccaggtgaaacagtgccagcttttcagttggcattccaggcagaggtgggcaatcatcctaccatagaggacgtgcaggcgcttgtgtccagagaaaaagaaagacccaaattcccagaagcctggaaagagaatagcctgacggtgcattctctgaaagagacgatggaagactgctgggatcaggacgcagaagcccgactgacagctcagtgtgcggaggagcgactcgccgaactgctcctcatctgggacagagaaaaatcagccagtcctgctcttaaccacagcactgcactgcagacacctaaagtaggttctgtccttgaacactcatccacaaacactgaagaccatgaaggtgctgataaaccactccacaatgacacctcagtggagcaaacctcagcaggaggaactaatcccgcagagaagaaaaagaactgcatcaactatgagtggcagcaggcccaatcacgacagtctggcacagaaagctcctccgctactcctgtctcagagtcctgcaatgccacgcatcctccaccagtggccaccaatctctgtgctcagctgacacgagaagacctggagataccaaaactcgacccgagcgaagtccagaggaacatgagagagagctcggacgagagccttatggaacattcacagaaacagttctgctctcctgaaacgtcagcctcccacggcccgttttatcctctcatgaagatggtttctgaggtttcggggtcacaaggatccagtaggcatggggacacccctataaccatcttacctaagcagcagaacgtccctaagagacccagtagcctcagtctccacgtaaagcctgggaaaacatccacctcgacgtcttcatcgctgcggatgaagttcggaaagcttggaaagtcaaatctgaagaaggttgagatgggtgtggctaaaagcagtgtggtgaatgcgacgcatgaagcccggctgattacagttgccaacaacgatgcggcggcagcaatgaataaatatgcaacagaaccagcagccgagtcagcaggaagttcagccaacgaggacttgaccttcggcctcttaaacaccagtcccgatgagcaggagcctctgctgagaagagaggcgcatcctgacaatgcaaacaacaacaacagcaataacaacaatggtgagggagatggtgatggagagacagagggtggaggagagggaggagagaacaatgagagtgtcggtccgacaggggatgcttcgtcgtcttctacggtggagccggttgtgctgcctggcccgtgttctgcttcaaccgccattcctccacaggcccaaagccagacacagacacacggagaggctctgctcagacagaaccgagtgcgcagaccagagagacccaactctctggatctgtccatcacaacactgccattactaggaggcaggtctgtcggcgatgggacagaggggtcaggggataaaatcaagaagcgggtaaagacgccttacgcacttaagaagtggcgtcctgccagctgggttatcaccactgacacactggatgccgaagtcaacaacaacagtcgtcacggaggaggcctcgggcagaatcagaatcaggctgggaccagcagacctaaatcagcttcggctgtctatctaggcagtcgaggaggatctcgcttctcagatcctaatgactgtgactttggtggaggaggttctggaggcggtggaagtggtggcggaggtagccctgactacagtctcgtgaaggctctgcaaatggcacaacagaattttgtcattacagacgcctccctcccagacaaccctatcgtctacgccagtagagggtttctgacactga

caggctattctctcgaccagatcctgggcaggaactgcaggtttctgcaagggccagaaacagacccaagagctgtggataagatcagga
atgccatcaccaaaggcgttgataccagtgtctgtctgctgaattatagacaggatggcacaaccttctggaatctcttcttcgtggctggactc
agagattctaagggcaatattgtcaactacgtcggagtgcagtcaaaggtgagcgaagattatgccaagctgctggtcaacgagcagaaca
ttgagtacaaaggtgtgcgcaccagtaacatgctgcgcagaaagcccgggtctagttatccgtacgacgtaccagactacgcataa

*bOpto-2A-Nodal* (Addgene #232640)

Key: **Myr**, **linker**, **Acvr1b**, **LOV**, **HA tag**, **P2A**, **Acvr2b**, **FLAG tag**

atggggagtagcaagagcaagcctaaggaccccagccagcgcggtggaggaggttctggaggcggtggaagtggtggcggaggtag
ccaccggcagcgactggatgttgaggatccatcctgtgatcacctgtacttggccaaagacaagaccttacaggatctcatcttcgatctgtc
cacctctggttcagggtctgggctgcccttgttcgttcagaggactgtggccaggacaattgtactgcaggagatcataggaaaaggtcgctt
tggggaagtgtggagggggagatggagaggaggtgatgtggctgtgaagatcttctcatccagggaggaacgttcctggttccgtgaagc
tgagatttaccaaaccatcatgctccgccatgaaaacatcttgggcttcattgctgctgataataaagacaatggcacatggacacagctgtg
gctagtgtcagactaccatgagcatggctcattatttgactatctcaaccactactccgtcacaatcgaggggatgatcaagttatcgctttcag
ccgccagcggtctggcacatctgcacatggagatcctgggcacacaggggaaaccaggcatcgcacatcgtgacctcaaatccaaaaac
atcttggtgaaaaagaatggtacttgcgccatagcagatctagggctcgctgtacggcacgagtccatcactgatactatagacattgcaccc
aatcagagggtgggcactaaaagatatatggccccagaggtactagatgaaacgatcaacatgaagcattttgattctttcaagtgtgctgat
atctatgctttgggactggtatattgggagattgcacgacggtgtaatgctggaggtatccatgaagattatcagctaccctactatgacctggt
gccgtccgacccctccatagaagagatgaggaaggtggtgtgtgaccagaggcttcggcctaatgtgcccaactggtggcagagctatga
ggcgctgagagtgatgggaaagatcatgcgggagtgttggtatgccaacggagcggcgcggcttacagctctgcggattaagaagactct
ttcgcaactcagcgtccaagaagacattaagggtggaggaggttctggaggcggtggaagtggtggcggaggtagccctgactacagtct
cgtgaaggctctgcaaatggcacaacagaattttgtcattacagacgcctccctcccagacaaccctatcgtctacgccagtagagggtttct
gacactgacaggctattctctcgaccagatcctgggcaggaactgcaggtttctgcaagggccagaaacagacccaagagctgtggataa
gatcaggaatgccatcaccaaaggcgttgataccagtgtctgtctgctgaattatagacaggatggcacaaccttctggaatctcttcttcgtg
gctggactcagagattctaagggcaatattgtcaactacgtcggagtgcagtcaaaggtgagcgaagattatgccaagctgctggtcaacg
agcagaacattgagtacaaaggtgtgcgcaccagtaacatgctgcgcagaaagcccgggtctagttatccgtacgacgtaccagactacg
caggaagcggagctacaaacttcagcctgctgaagcaggctggagacgtggaggagaaccctggacctatggggagtagcaagagcaa
gcctaaggaccccagccagcgcggtggaggaggttctggaggcggtggaagtggtggcggaggtagcaaacctccgtacggacatgtg
gacgtcaatgaggatccaggcccatctcctccatctcctctggtgggtctgaagcctctgcagctgctggaggttaaagctcgcggacgctt
cggctgcgtctggaaggctcagatgatcaatgaatatgtagctgtcaagattttccccattcaggataagctgtcgtggcagaacgagcggg
agatgttttccactccgggaatgaaacatgataacctgctgcgcttcatcgctgctgagaaacgcggatctaacctggagatggagttctggc
tcatcactgagtttcatgagcggggctctctgacggactatctgaaggggaacgcagtgagctgggctgatctgtgtgtgatagcggagag
catggcctgtggtctggcgtatctgcatgaagacgtgccgcgctccaaaggagaaggccccaaaccagccatcgcacacagagacttcaa
gagcaagaatgtgatgctgaagatggacctcaccgccgtcattggggattttgggctggcggtgcggtttgagccggggaaaccgccggg
agacacacatggccaggtgggcacgaggaggtacatggccccggaggttctggaaggagccataaacttccagcgggactcctttctgc
ggatagacatgtacgccatgggcctggtgctgtgggagctggtgtcacgctgcaaagctgctgatggtcctgtggacgagtacatgctgcc

gtttgaggaggagatcggtcagcacccgtcgctggaggatctgcaggatgctgtggtccataagaagctgcggccggcgtttaaggactgctggctcaagcattcaggtctgtgtcagatgtgcgagaccatggaggagtgctgggatcatgacgcagaggctcgtctgtcggccggctgtgtgcaggagcgcatctctcagatccgccgcgtcagcagctccacctcagactgcctgttctccatggtgacctcgctcaccaacgtggacctgccgcccaaagagtccagcatcggcggcggcggcagtggcggcggcggctctggcggcggcggctctcctgactacagtctcgtgaaggctctgcaaatggcacaacagaattttgtcattacagacgcctccctcccagacaaccctatcgtctacgccagtagagggtttctgacactgacaggctattctctcgaccagatcctgggcaggaactgcaggtttctgcaagggccagaaacagacccaagagctgtggataagatcaggaatgccatcaccaaaggcgttgataccagtgtctgtctgctgaattatagacaggatggcacaaccttctggaatctcttcttcgtggctggactcagagattctaagggcaatattgtcaactacgtcggagtgcagtcaaaggtgagcgaagattatgccaagctgctggtcaacgagcagaacattgagtacaaaggtgtgcgcaccagtaacatgctgcgcagaaagcccgggtctagtgactacaaggacgacgacgacaagtga

*GFP* (modified from (Lim et al., 2009))

atgagtaaaggagaagaacttttcactggagttgtcccaattcttgttgaattagatggtgatgttaatgggtacaaattttctgtcagtggagag
ggtgaaggtgatgcaacatacggaaaacttacccttaaatttatttgcactactggaaaactacctgttccatggccaacacttgtcactactct
cacttatggtgttcaatgcttttcaagatatccagatcatatgaagcggcacgacttcttcaagagcgccatgcctgagggatacgtgcagga
gaggaccatcttcttcaaggacgacgggaactacaagacacgtgctgaagtcaagtttgagggagacaccctcgtcaacaggatcgagctt
aagggaatcgatttcaaggaggacggaaacatcctcggccacaagttggaatacaactacaactcccacaacgtatacatcatggccgaca
agcaaaagaacggcatcaaagccaacttcaagacccgccacaacatcgaagacggcggcgtgcaactcgctgatcattatcaacaaaata
ctccaattggcgatgaccctgtccttttaccagacaaccattacctgtccacacaatctgccctttcgaaagatcccaacgaaaagagagacc
acatggtccttcttgagtttgtaacggctgctgggattacacatggcatggatgaactatacaaataa

*mScarlet-I3* (modified from (Gadella et al., 2023))

atggacagcacagaggctgtgatcaaggagttcatgagattcaaggtgcacatggagggaagcatgaacggacacgagttcgagatcga
gggagagggagagggaagaccttacgagggaacacagacagctaagctgaaggtgacaaagggaggacctctgcctttcagctggga
catcctgagccctcagttcatgtacggaagcagagctttcatcaagcaccctgctgacatccctgactactggaagcagagcttccctgagg
gattcaagtgggagagagtgatgatcttcgaggacggaggaacagtgagcgtgacacaggacacaagcctggaggacggaacactgat
ctacaaggtgaagctgagaggaggaaacttccctcctgacggacctgtgatgcagaagagaacaatgggatgggaggctagcacagag
agactgtaccctgaggacgtggtgctgaagggagacatcaagatggctctgagactgaaggacggaggaagatacctggctgacttcaa
gacaacatacaaggctaagaagcctgtgcagatgcctggagctttcaacatcgacagaaagctggacatcacaagccacaacgaggacta
cacagtggtggagcagtacgagagaagcgtggctagacacagcacaggaggaagcggaggaagctaa

*nls-Kaede*

Key: **Nuclear localization signal**, **Kaede**

atgcctaagaagaagagaaaggtgatgagtctgattaaaccagaaatgaagatcaagctgcttatggaaggcaatgtaaacgggcaccagtttgttattgagggagatggaaaaggccatccttttgagggaaaacagagtatggaccttgtagtcaaagaaggcgcacctctcccttttgcctacgatatcttgacaacagcattccattatggtaacagggtttttgctaaatacccagaccatataccagactacttcaagcagtcgtttcccaaagggttttcttgggagcgaagcctgatgttcgaggacgggggcgtttgcatcgctacaaatgacataacactgaaaggagacacttttttaacaaagttcgatttgatggcgtaaactttcccccaaatggtcctgttatgcagaagaagactctgaaatgggaggcatccactgagaaaatgtatttgcgtgatggagtgttgacgggcgatattaccatggctctgctgcttaaaggagatgtccattaccgatgtgacttcagaactacttacaaatctaggcaggagggtgtcaagttgccaggatatcactttgtcgatcactgcatcagcatattgaggcatgacaaagactacaacgaggttaagctgtatgagcatgctgttgcccattctggattgccggacaacgtcaagaagagacctgctgctaccaagaaagctggacaggccaagaagaagaaactggactag
